# What we talk about when we talk about color

**DOI:** 10.1101/2020.09.29.319517

**Authors:** Colin R. Twomey, Gareth Roberts, David H. Brainard, Joshua B. Plotkin

## Abstract

Names for colors vary widely across languages, but color categories are remarkably consistent (Berlin & Kay, 1969). Shared mechanisms of color perception help explain consistent partitions of visible light into discrete color vocabularies (Regier et al., 2007). But the mappings from colors to words are not identical across languages, which may reflect communicative needs – how often speakers must refer to objects of different color (Gibson et al., 2017). Here we quantify the communicative needs of colors in 130 different languages by means of a novel inference algorithm. We find that communicative needs are not uniform: some regions of color space exhibit 30-fold greater demand for communication than other regions. The regions of greatest demand correlate with the colors of salient objects, including ripe fruits in primate diets. Our analysis also reveals a hidden diversity in the communicative needs of colors across different languages, which is partly explained by differences in geographic location and the local biogeography of linguistic communities. Accounting for language-specific, non-uniform communicative needs improves predictions for how a language maps colors to words, and how these mappings vary across languages. Our account closes an important gap in the compression theory of color naming, while opening new directions to study cross-cultural variation in the need to communicate different colors and its impact on the cultural evolution of color categories.

## The color word problem

What colors are “green” to an English speaker? Are they the same as what a French speaker calls “vert?” Berlin & Kay [1] and Kay et al. [4] studied this question on a world-wide scale, surveying the color vocabularies of 130 linguistic communities using a standardized set of color stimuli (Fig. 1a). They found that color vocabularies of independent linguistic origin are remarkably consistent in how they partition color space [1]. In languages with two major color terms, one term typically describes white and warm colors (red/yellow) and the other describes black and cool colors (green/blue). If a language has three color terms, there is typically a term for white, a term for red/yellow, and a term for black/green/blue. Languages with yet larger color vocabularies remain largely predictable in how they partition the space of perceivable colors into discrete terms [5–8] (Fig. 1b).

**Figure 1.**
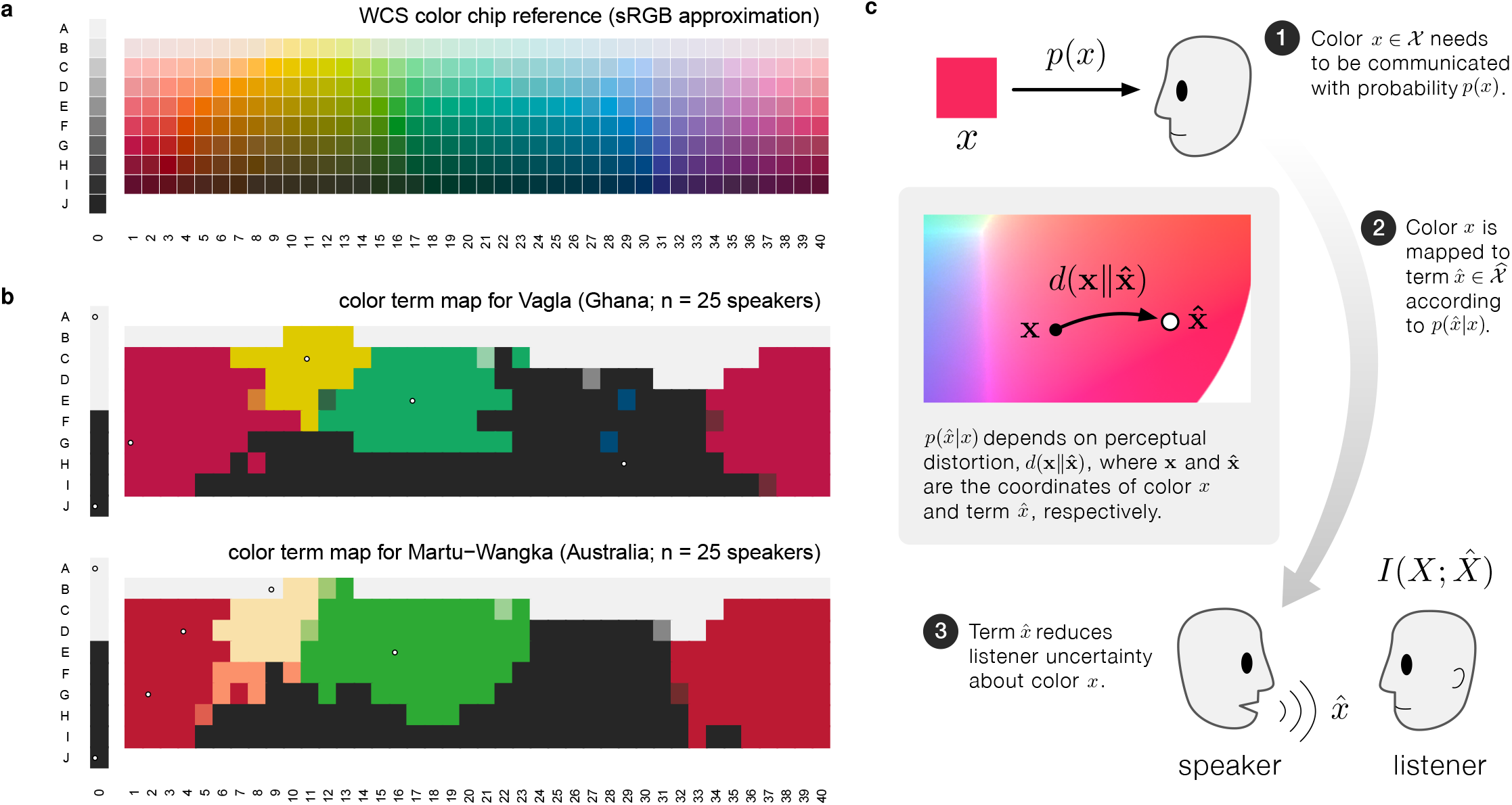
Cross-linguistic patterns in color naming and the rate-distortion hypothesis. Berlin & Kay (B&K) [1] and the World Color Survey (WCS) [4] studied color vocabularies in 130 languages around the world (see Methods: World Color Survey). **(a)** The 330 color chips named by native speakers in the WCS study. **(b)** Empirical color vocabularies for two example languages in the WCS, each with 6 basic color terms. Color chips correspond to panel (a) but they have been colored according to the focal color of the term chosen by the majority of speakers surveyed^†^. The languages Vagla and Martu-Wangka, although linguistically unrelated and separated by nearly 14,000 km, have remarkably similar partitions of colors into basic color terms [4]. **(c)** Schematic diagram of rate-distortion theory applied to color naming. A speaker needs to refer to color *x* with probability *p*(*x*). The speaker uses a probabilistic rule 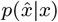 to assign color terms, 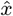, to colors, *x*. This rule depends on the perceptual distortion 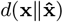 introduced by substituting 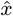 for the true color, *x*, where each term 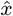 is associated with a coordinate in color space. The choice of the term 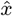 by the speaker reduces the listener’s uncertainty about the true color being referenced, measured on average by the mutual information 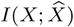. While any probabilistic mapping from colors to terms, 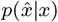, is possible, some mappings are more efficient than others. Rate-distortion theory provides optimal term mappings that allow a listener to glean as much information as possible, for a given level of tolerable distortion and distribution of communicative needs *p*(*x*).

What explains these shared patterns? To talk about color, a language must represent the vast space of human perceivable colors with a comparatively small set of color terms. The compression theory of color naming [2, 9–11] seeks to explain color vocabularies as an efficient mapping from colors to terms, based on the psychophysics of human color perception and the utility, or need, to reference different colors.

Judgements of color appearance by humans with normal color vision are remarkably stable despite genetic variability in photoreceptor spectral sensitivities [12], age-dependent variability in light filtering of the eye [13], and variation in the proportion of different classes of retinal cone photoreceptors [14, 15]. The shared psychophysics of perception therefore provides a common metric for color similarity, and common limits on the gamut of perceivable colors, which each contribute to shared patterns in color naming across languages [1, 2, 16–23].

Recent work [3, 24, 25] has also found that color terms tend to reflect how often speakers need to refer to different colors, with a trend that emphasizes communication about warm hues (red/yellow) over cool hues (blue/green). Shared communicative needs of colors – e.g., emphasizing the colors of greatest importance to ancestral humans, such as those of ripe fruits or dangerous animals [26] – also helps explain shared patterns in color naming across languages.

However, estimating communicative needs is non-trivial. Several approaches have been proposed: using the statistics of surface reflectances in natural scenes [9]; assuming a uniform distribution over highly saturated colors [2]; using a world-wide average of capacity-achieving priors [11, 27]; or extrapolating from English word-frequency corpus data [28]. The aim of all of these approaches is to approximate a single distribution of needs common to all languages worldwide. But unlike perception, needs may vary across cultures – and this variation might explain why color vocabularies, though similar, are far from identical across languages. A complete theory of color naming must explain cross-language variation as well as shared trends. But to date, language-specific communicative needs are unknown.

Here, we seek to close this gap in the compression theory of color naming by providing a new way to directly estimate language-specific communicative needs of colors. Without making strong assumptions about the origins or characteristics of communicative needs, we derive a novel algorithm to solve a natural inverse problem; we infer the most conservative (maximum entropy) distribution of communicative needs across colors consistent with positions of focal colors in a language’s vocabulary – e.g. the “reddest red” and the “greenest green.” Our approach explains focal colors as a natural part of the compression theory of color naming, and it allows us to test predictions for term maps against independent empirical data that was not used in fitting our model.

Applying our method we infer the language-specific communicative needs for 130 languages around the world. We confirm that shared trends in communicative needs across languages are related to the colors of salient objects [3], but we also find substantial variation in communicative needs across languages. This variation is consequential: accounting for variation in needs substantially improves the prediction of color terms in each language. Moreover, this variation in needs across linguistic communities is meaningful: it correlates with differences in geographic location and local biogeography. Our account supports an emerging, unified view of the color word problem that integrates the shared psychophysics of color perception with language-specific communicative needs for colors. We show that this view is consistent with both shared patterns and observed variability across languages.

## Color naming as a compression problem

In the compression model of color naming first introduced by Yendrikhovskij [9] and with recent extensions by Zaslavsky et al. [11], a color in the set of all perceivable colors, 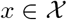, needs to be communicated with some probability, *p*(*x*), to a listener. The speaker cannot be infinitely precise when referring to *x*, and must instead use a term, 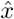, from their shared color vocabulary, 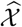. Many colors in 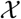 map to the same term, so that a listener hearing 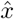 will not know exactly which color *x* was referenced. Color naming is then distilled to the following problem: how do we choose the mapping from colors to color terms? Rate-distortion theory [29–32], the branch of information theory concerned with lossy compression, provides an answer.

Mapping colors to a limited set of terms necessarily introduces imprecision or “distortion” in communication. The amount of distortion depends on a listener’s expectation about what color, *x*, a speaker is referencing when she utters color term 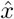. Under the rate-distortion hypothesis, a language’s mapping from colors to terms allows a listener to glean as much information as possible about color *x* from a speaker’s choice of term 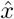 (Fig. 1c).

Each color 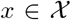 is identified with a unique position, denoted x, in a perceptually uniform color space. Here we use CIE Lab as in Regier et al. [2]. The coordinates corresponding to a color term 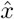 are given by its centroid: the weighted average of all colors a speaker associates with that term, 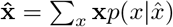. The distortion introduced when a speaker uses 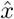 to refer to *x* is simply the squared Euclidean distance between x and 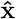 in CIE Lab, denoted 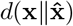. Intuitively, colors that are near 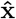 are more likely to be assigned to the term 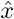 than colors that are far (Fig. 1c).

The mathematics of compression provides optimal ways to represent information for a given level of tolerable distortion. The size of a compressed representation, 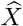, is measured by the amount of information it retains about the uncompressed source, *X*, given by the mutual information 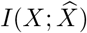. Terms represent colors by specifying the probability of using a particular term 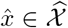 to refer to a given color 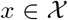, denoted 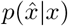. Rate-distortion efficient mappings are choices of the mapping 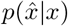 that minimize 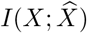 such that the expected distortion, 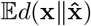, does not exceed a given tolerable level. Efficient mappings and centroid positions can be found for a large class of distortion functions known as Bregman divergences, which includes the CIE Lab measure of perceptual distance (SI Sec. A).

## Communicative needs of colors

Rate-distortion theory provides an efficient mapping from colors to terms that depends on three choices: the distortion function in color space, the degree of distortion tolerated by the language, and the probability *p*(*x*) that each color needs to be referenced during communication, called the “communicative need.” Previous studies have largely focused on communicative needs that are shared across all languages, considering distributions that are either uniform across the WCS color stimuli [7], correlated with the statistics of natural images [9] or the color of salient objects [3], approximated by a world-wide average “capacity achieving prior” [11, 27], or related to linguistic usage [33] as e.g. approximated by the frequencies of words for color in English-language corpus data [28]. As a result, prior studies have drawn conflicting conclusions about whether communicative needs matter for color naming, and little is known about whether communicative needs vary across languages or whether such variation is significant for their color vocabularies. While the potential importance of language-specific communicative needs has been discussed [33], here, for the first time, we resolve these questions by directly estimating the communicative needs of colors for each of the 130 languages in the combined B&K+WCS dataset under the compression theory of color naming.

## Algorithm to infer communicative needs

How can we infer the underlying communicative needs of colors from limited empirical data? Here we derive an algorithm that finds the maximum-entropy estimate of the underlying communicative needs *p*(*x*) consistent with a rate-distortion optimal vocabulary with known centroid coordinates 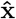 and term frequencies 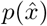, for any Bregman divergence measure of distortion.

The estimate of communicative needs has the form 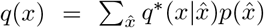, with

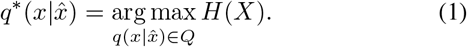

In words, the optimal 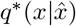 is the choice of 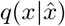 that maximizes the entropy, *H*(*X*), among the the set of conditional probability distributions *Q* whose predicted focal color coordinates match the observed coordinates for each color term. We construct this solution via a novel iterative alternating maximization algorithm (see SI Sec. B for its derivation),

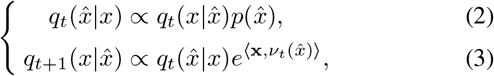

where the vectors 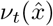 are chosen so that predicted focal color coordinates match observed coordinates (SI Sec. B).

This algorithm provably converges to a unique, globally optimal, maximum-entropy estimate of the true communicative need *p*(*x*) (SI Sec. B.1 and B.2). Remarkably, we can construct this solution knowing only that the observed coordinates 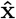 are rate-distortion optimal centroids, without knowledge of the specific distortion measure (SI Sec. B.3; SI Fig. B1).

## Inference from focal colors

Our algorithm infers a language’s communicative needs from knowledge of the centroids associated with its color terms. Berlin & Kay measured the “focal color” of each color term by asking native speakers to choose from among the Munsell stimuli (Fig. 1a) the “best example” of that term. We propose that the measured focal colors are in fact the centroids for each term.^†^ This hypothesis may appear problematic since laboratory experiments suggest focal colors and category centroids are distinct points in color space [34–36]. However, centroids in those studies were calculated under the implicit assumption of uniform communicative needs, leaving open the possibility that focal colors are centroids under the true distribution of non-uniform needs (SI Sec. A.3).

Our approach provides an unbiased inference of language-specific communicative needs that does not make strong assumptions about the form of *p*(*x*) or depend on additional, unmeasured quantities for each WCS language. Prior work on a universal distribution of needs relies on strong assumptions about the form of *p*(*x*) (see SI Sec. B.3, & D), and so applying it to individual languages in the WCS produces implausible inferences (SI Fig. C4ab). Alternatively, there is a prior language-specific approach based on word frequency data, but this approach cannot be applied to the vast majority of languages in the WCS that lack this information (SI Sec. D.1, SI Fig. D10b). Moreover, unlike prior work, our inference of language-specific needs does not rely on knowing the empirical mapping from colors to terms, 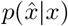, which is the quantity that we ultimately wish to predict from any theory of color naming (SI Fig. D9).

## Different colors, different needs

Our analysis reveals extensive variation in the demand to speak about different regions of color space (Fig. 2a). Averaged over all 130 B&K+WCS languages, the inferred communicative needs emphasize some colors (e.g. bright yellows and reds) up to 36-fold more strongly than others (e.g. blue/green pastels and browns). This conclusion stands in sharp contrast to prior work that assumed a uniform distribution of needs [2] and attributed color naming to the shape of color space alone.

**Figure 2.**
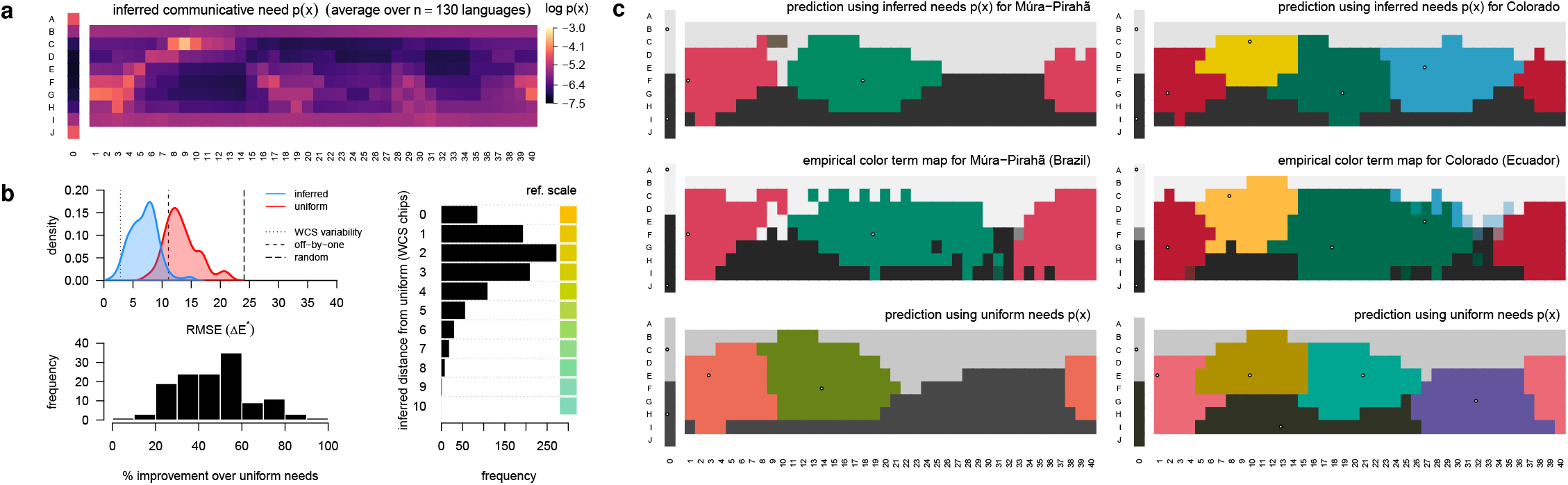
Inferred distributions of communicative needs. **(a)** The mean inferred distribution of communicative need, *p*(*x*), averaged across the WCS and B&K survey data (*n* = 130 languages). Color chips correspond to those shown in Figure 1a. We infer 36-fold variation in communicative need across color chips, with greater demand for communication about yellows and reds, for example, than for blues and greens. **(b)** The color vocabulary of a language predicted by rate-distortion theory better matches the empirical vocabulary when we account for variation in the need to communicate about different colors. (*Top-left*) The error between the predicted and empirical focal color positions across *n* =130 languages, where predictions are rate-distortion optimal vocabularies assuming either a uniform (red) or the inferred (blue) distribution of communicative needs. Root mean square error (RMSE) is measured in units of CIE Lab perceptual distance (denoted ΔE^✶^; Methods: RMSE of focal color predictions). Reference lines show RMSE when empirical focal points are compared to random focal points (*random*), displaced by one WCS column or row (*off-by-one*), and by sampling from participant responses (*WCS variability*) (see SI Sec. C.1). (*Bottom-left*) The relative improvement (reduction in error) using the inferred versus uniform distribution of communicative need. (*Right*) Difference in focal point positions of rate-distortion optimal vocabularies, under inferred versus uniform communicative needs. **(c)** Two example languages, Múra-Pirahã (*left*), and Colorado (*right*), that illustrate how predicted term maps are improved when accounting for non-uniform communicative needs of colors. The region corresponding to each term is colored by the WCS chip closest to the term’s focal point (white points). (*Top row*) The predicted term maps based on the inferred distribution of communicative needs; (*Bottom row*) based on a uniform distribution of communicative need; (*Middle row*) the empirical term maps in the WCS data.

Our ability to predict the color vocabulary of a language is substantially improved once we account for non-uniform communicative needs (Fig. 2b). We find improvement in an absolute sense, as measured by the root mean squared error (RMSE) between predicted and empirically measured focal colors, and also in a relative sense, measured by percent improvement over a uniform distribution of needs. The typical change in predicted focal color once accounting for non-uniform needs is easily perceivable, corresponding to a median change of two WCS color chips (Fig. 2b right). Not only are the predicted focal points in better agreement with the empirical data, once accounting for non-uniform needs, but the entire partitioning of colors into discrete terms is substantially improved, as seen in the example languages Múra-Pirahã and Colorado (Fig. 2c).

We infer communicative needs and predict color terms using data from the first of two experiments in the WCS, which measured focal colors (Fig. 3a). This inference and prediction requires fitting one parameter that controls the “softness” of the partitioning and one hyperparameter to control over-fitting (SI Sec. C). Without any additional fitting, we can then compare the predicted mappings from colors to terms to the empirical term maps measured in the second WCS experiment. For nearly all of the WCS languages analyzed (*n* = 110), the color term maps predicted by rate-distortion theory are significantly improved once accounting for non-uniform communicative needs (improvement in 84% of languages, Fig. 3b). Only 15% of languages show little or no improvement, with an additional 1 outlier, Huave (Huavean, Mexico), that may violate model assumptions in some significant way (see Discussion). The substantial improvement in predicted term maps can be attributed both to universal patterns in communicative needs, shared across languages, and to language-specific variation in needs (Fig. 3c).

**Figure 3.**
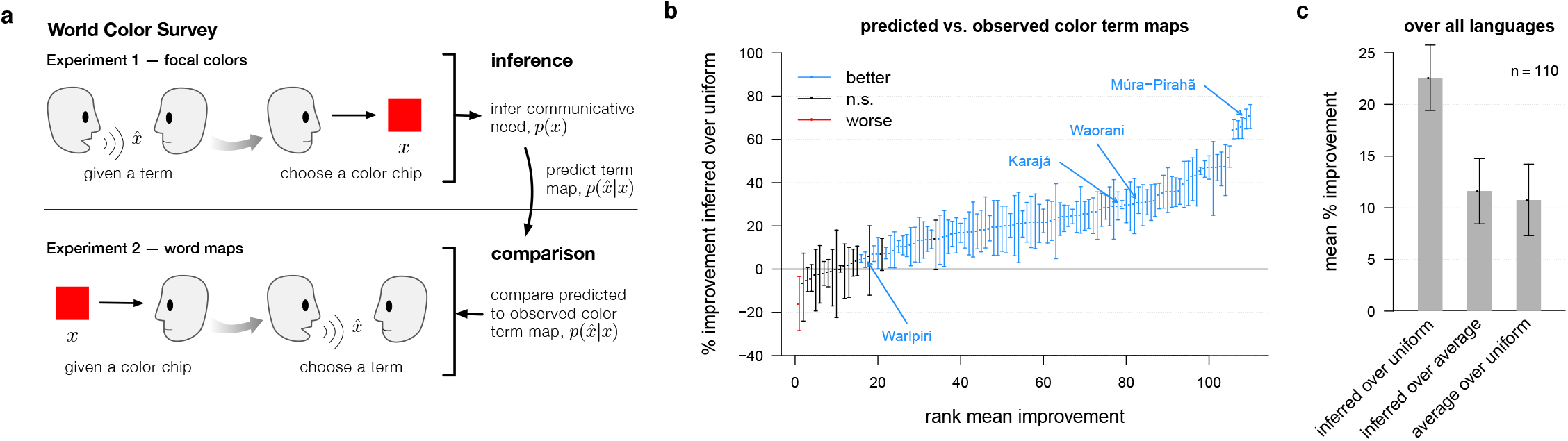
Inference and prediction within the World Color Survey. **(a)** The World Color Survey (WCS) [4] included two separate experiments with native speakers of each language. In this study we used only the WCS focal color experiment to infer the communicative needs of colors, *p*(*x*), and to predict a language’s mapping from colors to terms, 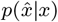. Without any additional fitting we then compared the predicted term maps to the empirical term maps observed in the second WCS experiment. **(b)** Predicted term maps tend to agree with the observed term maps (see Figure 2c; SI Fig. C2c). Moreover, the predicted term maps show better agreement with the empirical data than would predictions assuming a uniform distribution of communicative needs. The panel shows the rank ordered mean percentage improvement in predicted versus observed term maps using the inferred communicative need *p*(*x*) compared to a uniform communicative need, with 95% confidence intervals (bootstrap resampling, see Methods: Measuring distance between distributions over colors). Languages (points) colored black have 95% confidence intervals overlapping 0%; blue indicates significant improvement. Languages that do worse under the inferred distribution of needs (red points) violate model assumptions. **(c)** Over all languages, the mean percentage improvement (and 95% CIs) in predicted vocabularies when using language-specific commutative needs compared to uniform needs (“inferred over uniform”), language-specific versus average needs over all languages (“inferred over average”), and average versus uniform needs (“average over uniform”). Some improvement in predictive accuracy is attributable to commonalities in communicative needs across languages (third comparison), and yet more improvement is attributable to variation in needs among languages (second comparison).

In contrast to prior work on the compression model of color naming [11, 28], no part of our inference procedure uses empirical data on a language’s mapping from colors to terms, 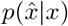.^†^ Nor are our predicted color terms simply an out-of-sample prediction, since the predicted quantities, 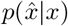, are not used to parameterize the model. And so our analysis is not simply a fit of the compression model to data, but rather an empirical test of its ability to predict color naming from first principles.

## Communicative needs and the colors of salient objects

We can interpret the inferred communicative needs of colors by comparing them to what is known about the colors of salient objects. Prior work [3] suggests a warm-to-cool trend in communicative need, related to the frequency of colors that appear in foreground objects as identified by humans in a large dataset of natural images [37] (Fig. 4a). We find that the same correlation holds, at least when restricting to the middle range of lightness (color chips in rows C–H; two-sided Spearman’s *ρ* = 0.3, *p* < 0.001, *n* = 240). However the pattern of communicative needs is more complex than this warm-cool gradient alone. Pastels that are greenish blue or blue, as well as brownish-greens, need to be communicated less often than dark green or dark blue, for example. Moreover, dark colors in general (e.g. color chips in rows I-J) show a relatively high communicative need under our inference compared to their frequency in foreground objects of natural images (Fig. 4a).

**Figure 4.**
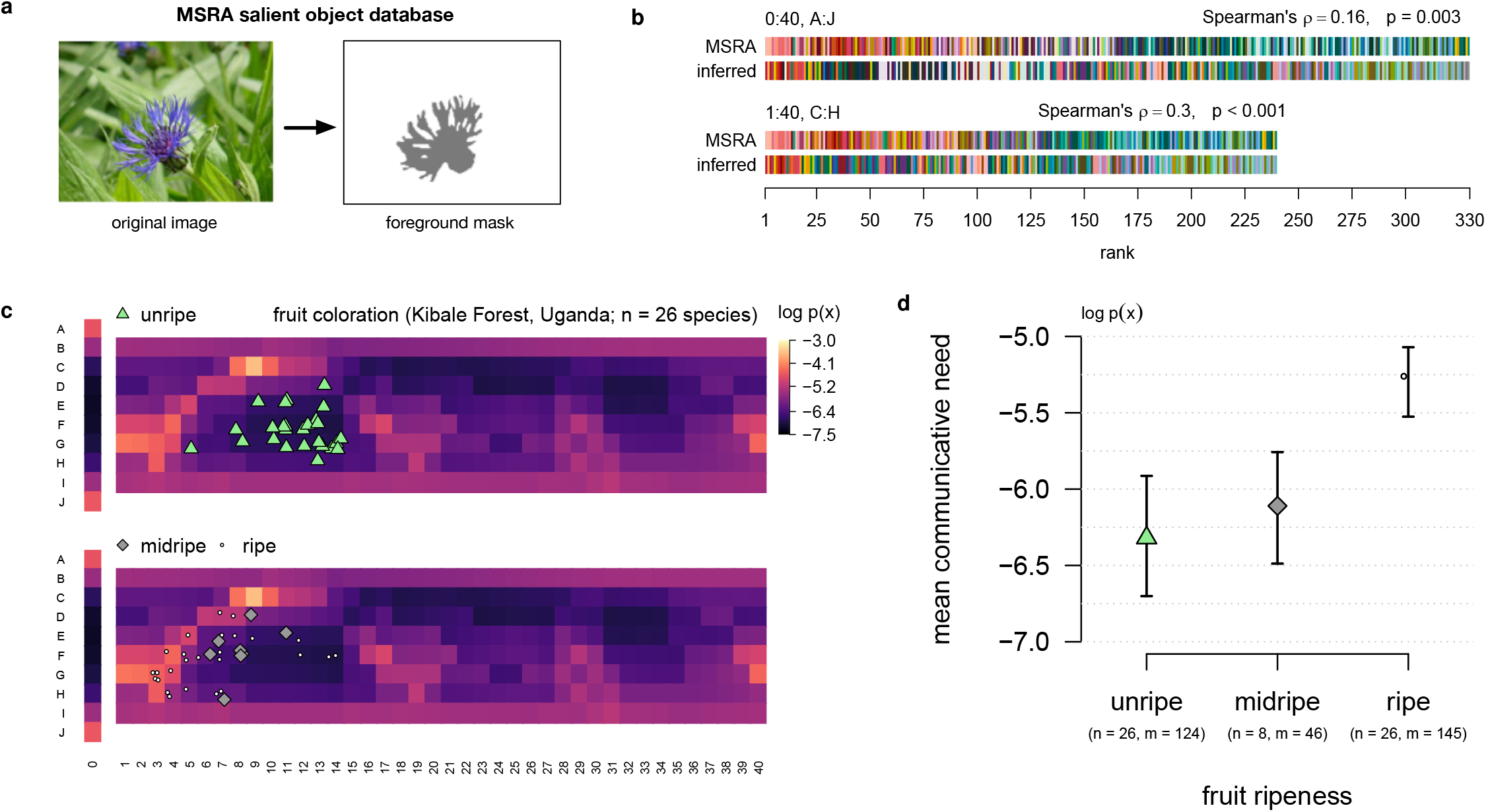
Inferred distributions of communicative needs correlate with the colors of salient objects. **(a)** Human participants in the Microsoft Research Asia (MSRA) salient object study were asked to identify the foreground object in 20,000 images; example foreground mask illustrated in gray. **(b)** WCS color chips ordered by their rank frequency in the foreground of MSRA images (rows “MSRA”; see Gibson et al. [3]), and in the inferred distribution of communicative need (rows “inferred”), averaged across the *n* =130 languages in the B&K+WCS survey data. There is a weak positive correlation between the colors that are considered salient, in the MSRA dataset, and the colors with greatest inferred communicative need, across all WCS color chips (*top*). This relationship is strengthened after removing achromatic chips (WCS column 0, rows B and I) from the comparison (*bottom*). **(c)** Colors of unripe (*top*), midripe, and ripe (*bottom*) fruit in the diets of catarrhine primates, derived from fruit spectral reflectance measurements collected in the Kibale Rainforest, Uganda, by Sumner & Mollon [38, 39]. The colors of ripe fruit tend to correspond with the colors of greatest inferred communicative need. **(d)** Average log-probability in the inferred distribution of communicative need of color corresponding to unripe, midripe, and ripe fruit. *n* denotes the number of fruit species, and *m* the total number of spectral measurements. Error bars show 95% confidence intervals of the means (nonparametric bootstrap by species).

We also compared communicative needs to spectral measurements by Sumner & Mollon [38, 39] of unripe and ripe fruit in the diets of catarrhine primates, which have trichromatic color vision and spectral sensitivities similar to humans. When projected onto the WCS color chips (see SI Fig. C6), unripe, midripe, and ripe fruit occupy distinct regions of perceptual color space (Fig. 4c) corresponding to low, medium, and high values of inferred communicative need, respectively (Fig. 4d). The morphological characteristics of fruit, including color, are known to be adapted to the sensory systems of frugivores that act as their seed dispersers, for vertebrates in general [40–42] and primates in particular [43–45]. And so our results support the hypothesis that shared communicative needs in human cultures emphasize the colors of salient objects that stand out or attract attention in our shared visual system across a typical range of environments^†^.

## Cross-cultural variation

Languages vary considerably in their needs to communicate about different parts of color space (Fig. 5a; SI Fig. E11–E27). The inferred needs for the language Waorani (Ecuador), for example, emphasize white and mid-value blues, while de-emphasizing yellows and greens, relative to the average needs of all B&K+WCS languages. Whereas Martu-Wangka (Australia) emphasizes pinks and mid-value reds, as well as a light greens, while de-emphasizing blues and dark purples (Fig. 5a). In fact, the median distance between language-specific communicative needs and the across-language average needs is nearly as large as the distance between the average needs and uniform needs (9.9 and 11.2, respectively, in units of ΔE^✶^).

**Figure 5.**
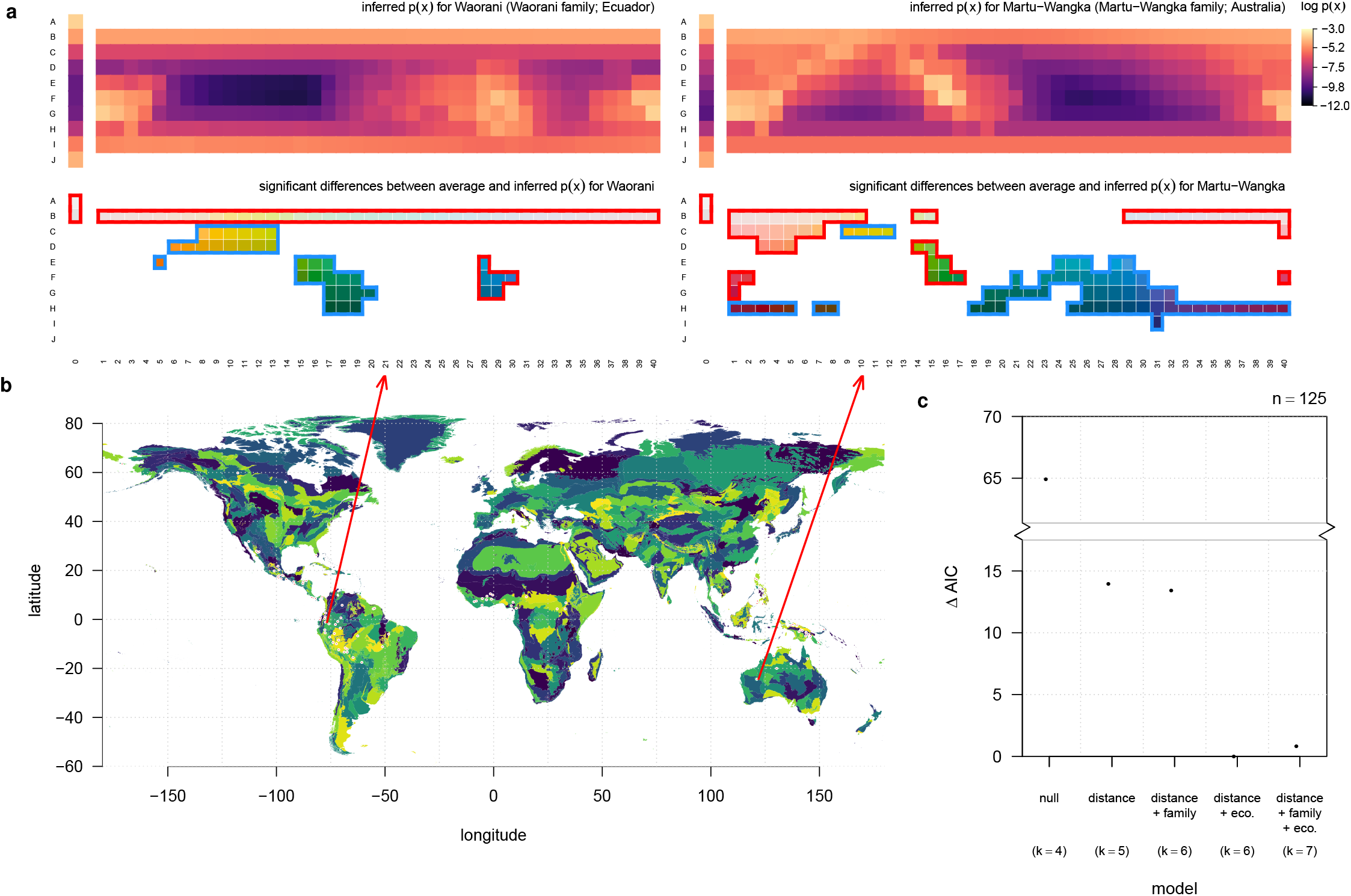
The communicative needs of colors vary across languages, and they are correlated with geographic location and ecological region. **(a)** The inferred distribution of communicative needs for two example languages (*top row*). For each language, many color chips have significantly elevated (red border) or suppressed (blue border) communicative need compared to the across-language average (*bottom row*; deviations that exceed *σ*/2 with 95% confidence are highlighted in red or blue). **(b)** The approximate locations of WCS native language communities (red points) shown on a world map colored by eco-regions [47]. **(c)** Languages spoken in closer proximity to each other and sharing the same eco-region tend to have more similar inferred communicative needs (Type II Wald Chi-square tests; χ^2^ = 20.98, df = 1, *p* < 0.001; and χ^2^ = 12.91, df = 1, *p* < 0.001, respectively), whereas shared language family does not have a significant effect (χ^2^ = 1.022, df = 1, *p* = 0.31). Distance and shared eco-region each substantially improve the fit of generalized linear mixed effects models (GLMMs) predicting the distance between pairs of inferred communicative needs. GLMMs were fit using log-normal link function and a random-effects model designed for regression on distance matrices [49] (see Methods: Correlates of cross-cultural differences in communicative need). *k* denotes the total number of fixed and random effects in each model.

Why do language communities vary in their needs to communicate different colors? Detailed study of this question requires language-specific investigation beyond the scope of the present work. However, we can at least measure how variation in linguistic origin, geographic location, and local biogeography (Fig. 5b) relate to differences in communicative needs. We quantified these factors for pairs of languages by determining: (1) whether or not they belong to the same linguistic family in *glottolog* [46]; (2) the geodesic distance between communities of native speakers; and (3) whether or not language communities share the same “ecoregion,” a measure of biogeography [47] that delineates boundaries between terrestrial biodiversity patterns [48]. Our statistical analysis also controls for differences in the number of color terms between languages, because we seek to understand cross-cultural variation above and beyond any relationship between vocabulary size and (inferred) communicative needs (SI Sec. C.3).

While language differences are largely idiosyncratic, we find a small but measurable impact of distance and biogeography on communicative needs (Fig. 5c, Methods: Correlates of cross-cultural differences in communicative need). In particular, increasing the geodesic distance between language communities by a factor of 10 decreases the mean similarity in their communicative needs by a factor of 2.9% ([1.7%, 4.2%] 95% CI), while sharing the same ecoregion increases the mean similarity by a factor of 8.4% ([3.9%, 12.7%] 95% CI). By contrast, we find no significant effect of language genealogy on communicative needs, at least at the coarse scale of language family. Taken together, these results suggest that color vocabularies are adapted to the local context of language communities.

## Discussion

We have inferred language-specific needs to communicate about different colors, using a novel algorithm that applies to any rate-distortion Bregman clustering. Accounting for non-uniform needs substantially improves our ability to predict color vocabularies across 130 languages. Neither our predictions of term maps nor, in contrast to prior work, our inferences of needs use empirical information on the mappings from colors to terms, allowing us to test the compression model of color naming against independent data.

The distribution of communicative needs, averaged across languages, reflects a warm-to-cool gradient, as hypothesized in Gibson et al. [3]; and it is related to object salience more generally, as indicated by the positioning of ripe fruit coloration in regions of highest need. This is true even though the needs *p*(*x*) that we infer by maximum entropy differ from the notion of communicative efficiency, or surprisal, used in prior work (SI Sec. C.6.1). We also document extensive variation across languages in the demands on different regions of color space, correlated with geographic location and the local biogeography of language communities.

Our analysis provides clear support for the compression model of color naming. Whereas prior work has established the role of shared perceptual mechanisms for universal patterns in color naming, our results highlight communicative need as a source of cross-cultural variation that must be included for agreement with empirical measurements. A catalogue of language-specific needs (SI Fig. E11–E27) will enable future study into what drives cultural demands on certain regions of color space, and how they relate to contact rates between linguistic communities, shared cultural history, and local economic and ecological contexts. Our methodology also provides a theoretical framework and inference procedure to study categorization in other cognitive domains, including other perceptual domains of diverse importance worldwide [50], and even in non-human cognitive systems that exhibit categorization (e.g. Zebra finches [51, 52], the songbird *Taeniopygia guttata*).

Several languages have been advanced as possibly invalidating the universality of color categories [53–55]. Languages are known to vary in the degree to which different sensory domains are coded [50, 56, 57], and in Pirahã and Warlpiri the existence of abstract terms for colors has been disputed [58, 59]. Moreover, the color vocabularies in Karajá and Waorani notably lack alignment with the shape of perceptual color space [7]. Once we account for communicative needs, however, we find that the color terms of Karajá and Waorani are well explained by rate-distortion theory. Likewise, while Pirahã may seem exceptional when assuming uniform communicative needs, we recover accurate predictions once accounting for a non-uniform distribution of needs (Fig. 2c, Fig. 3b)^†^.

Nevertheless, several languages show little or no improvement in predicted term maps using inferred versus uniform communicative needs, and Warlpiri is among these cases. Before drawing conclusions about exceptionalism, however, we note that several technical assumptions of our analysis may be violated for these languages. For one, we assumed that basic color terms are used with equal frequency, to first approximation. This is a reasonable assumption given that basic color terms are elicited with roughly equal frequency under a free naming task in e.g. English [36]. Moreover, the inferred distribution of needs for WCS and B&K languages are relatively insensitive to non-uniformity of color term frequency, up to variation by a factor 1.5 (SI Sec. C.2, SI Fig. C2d). Still, this assumption may not be accurate enough for all languages, and the frequencies of color terms requires future empirical study. Another possibility is that the choice of the WCS stimuli themselves, i.e. the set of Munsell chips, 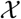, may work well for identifying focal colors of most languages, but may be too restrictive in the languages that show little improvement. Future field and lab work could remedy this by broadening the range of color stimuli used in surveys.

Another limitation of the WCS is variability in chroma across the Munsell color chips used as stimuli, which might bias participants’ choice of focal color positions [25, 61–64]. While there is no relationship between chroma and language-specific communicative needs (see SI Sec. C.6.2; SI Fig. C8b), we do find a small but statistically significant correlation (two-sided Spearman’s *ρ* = 0.13, *p* = 0.019, *n* = 330) between chroma and the inferred distribution of communicative need averaged across WCS languages. However, if this bias dominated the choice of focal colors in the WCS, then we would not expect distributions of need inferred from focal colors to improve predictions of color term maps. The fact that we do see substantial improvement suggests that whatever bias this effect may have, it is evidently not large enough to impact the relationship between focal color positions and color term maps for most languages. Nor would chromatic bias in stimuli explain the cross-cultural variation in communicative needs that we observe, since the set of stimuli was held constant across languages.

Our study has focused on how languages partition the vast space of perceivable colors into discrete terms, and how communicative needs shape this partitioning. Why some languages use more basic color terms than others remains an open topic for cross-cultural study. In principle, the issue of tolerance to imprecision in color communication is orthogonal to the distribution of communicative needs in a community. In practice, the number of color terms has a small impact on the resolution of inferred needs (SI Fig. C3a), which we control for in cross-cultural comparisons (SI Fig. C3b). Nonetheless, languages that have similar vocabulary sizes tend to have more similar communicative needs across colors, and this covariation is greater than any effect of vocabulary size on the resolution of our inferences (SI Fig. C3). These results suggest that causal factors driving vocabulary size may also influence a culture’s communicative demands on colors – a hypothesis for future research.

Future empirical work may begin to unravel why cultures vary in their communicative demands on different regions of color space. It is already known that natural environments vary widely in their color statistics [65, 66] and this variation matters for color salience [67]. The need to reference certain objects, as well as their salience relative to similar backgrounds, may help explain why communities that share environments prioritize similar regions of color space, as we have seen. And so shared environment, physical proximity, and shared linguistic history at a finer scale than language family, are all plausible avenues for future study on the determinants of color demands. Beyond these factors, there remains substantial interest in cultural features that we have not studied here, including religion, agriculture, trade, access to pigments and dies, and different ways of life, that can all shape a community’s needs to refer to different colors, and the resulting language that emerges.

## Methods

### World Color Survey

Berlin & Kay [1] and Kay et al. [4] surveyed color naming in 130 languages around the world using a standardized set of color stimuli. The stimuli (Fig. 1a), a set of Munsell^†^ color chips, were designed to cover the gamut of human perceivable colors at maximum saturation, across a broad range of lightness values. Native speakers were asked to choose among the basic color terms in their language to name each color chip, one at a time, in randomized order. The WCS study surveyed 25 native speakers in each of 110 small, pre-industrial language communities; the B&K study surveyed one native speaker in each of 20 languages from a mixture of both large (e.g. Arabic, English, and Mandarin) and comparatively small (e.g. Ibibio, Pomo, and Tzeltal) pre- and post-industrial societies.

The stimuli provided by the Munsell color chips are a function of the color pigment of the chips and the ambient light illuminating them. The ambient light source was approximately controlled by conducting the survey at noon and outdoors in shade, corresponding to CIE standard illuminant C. To the extent possible, participants were surveyed independently, although preventing the discussion of responses among participants was not always possible (discussed in Regier et al. [60]).

In our treatment of the color naming data, for each language we include all recorded terms that had an associated focal color, was used by at least two surveyed speakers (unless a B&K language, in which case only one speaker was surveyed), and was considered the best choice for at least one WCS color chip.

The 20 B&K languages were included in our analyses where appropriate: comparisons based on focal colors and inferred communicative needs. They were excluded from term map comparisons because the methods of estimating term maps differed methodologically from those in the WCS [68], and they do not provide straightforward estimates of 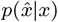. In addition, B&K languages with significant geographic extent, e.g. Arabic and English, were excluded from statistical analysis of the correlates of cross-cultural differences in communicative needs, because estimating geographic distance or local biogeography would make little sense for these languages.

### RMSE of focal color predictions

Language-specific focal color positions were compared to model predictions using the root mean squared error (RMSE) between observation and prediction in units of CIE Lab ΔE^✶^, computed for each WCS language *i* according to

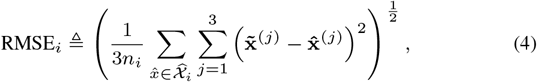

where the superscript (*j*) specifies the coordinate in the CIE Lab color space of position vectors 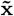 and 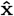, corresponding respectively to the predicted and empirically observed coordinates of the focal color for term 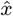 in language *i*’s vocabulary, 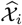. Here 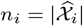 denotes the number of basic color terms in language *i*’s vocabulary.

### Spectral measurements of ripening fruit

Spectral measurements of ripening fruit in the diets of caterrhine primates were obtained from the Cambridge database of natural spectra.^‡^ Reflectance data for fruit taken from the Kibale Forest, Uganda, were converted to CIE XYZ 1931 color space coordinates using CIE standard illuminant C. We then converted points from XYZ to CIE Lab space using the XYZ values for CIE standard illuminant C (2°standard observer model) as the white point, in order to match the WCS construction of CIE Lab color chip coordinates. Calculations were performed in R (v3.6.3) using the package `colorscience` (v1.0.8).

Indicators of fruit ripeness include color, odor, and smell. Therefore, to measure visual salience we considered only fruit that had a discernable (in terms of CIE Lab ΔE^✶^) difference between unripe and ripe measurements (see Fig. C6a for determination of statistical threshold on change in chromaticity). For fruits with detectable changes in chromaticity, we projected their unripe, midripe, and ripe positions onto the WCS color chips such that absolute lightness, L^✶^, and the ratio of a^✶^ to b^✶^ was preserved (Fig. C6b).

### Measuring distance between distributions over colors

We quantified the perceptual difference between any two distributions over the WCS color chips in terms of their Wasserstein distance (used in Fig. 5c), defined as

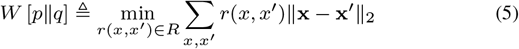

where *R* is the set of joint distributions satisfying 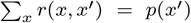 and 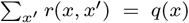. The CIE Lab coordinates of *x* and *x*′ are given by x and x′, respectively, and the Euclidean distance between them approximates their perceptual dissimilarity, by design of the CIE Lab system. Under this measure, a small displacement in CIE Lab space of distributional emphasis is distinguishable from a large displacement. For example, for discrete distribution *p*(*x*) = *α* if 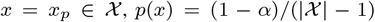 otherwise, let distribution *q*(*x*) be defined identically except substituting 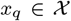 for *x_p_*. Then the Wasserstein distance between *p* and *q* will increase with the Euclidean distance between *x_p_* and *x_q_*, whereas e.g. the Kullback–Leibler divergence between *p* and *q* would remain constant for any *x_p_* ≠ *x_q_*.

We used a generalization of this distance measure to quantify the match between predicted and measured term maps. To make this comparison we find the minimum-CIE ΔE^✶^partial matching between predicted and measured term map categories, 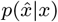, for each term 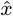 (used in Fig. 3b). To do this we find the minimum cost achievable by any assignment of chips empirically labeled by 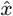 to those predicted to be labeled 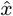, weighted by the measured and predicted 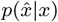. The best partial matching accommodates for the fact that predicted and measured categories can differ in total weight. This measure is known as the Earth mover’s distance [69] (EMD), which has the Wasserstein distance as a special case with matching total weights. Both measures were computed in R (v3.6.3) using the `emdist` (v0.3-1) package.

### Correlates of cross-cultural differences in communicative need

We modeled the pairwise dissimilarity in communicative need between B&K+WCS languages as a log-linear function of the geodesic distance between language communities, shared linguistic family, and shared ecoregion, using a maximum-likelihood population-effects model (MLPE) structure to account for the dependence among pairwise measurements [49]. For languages *j* = 2, …, *n*, *i* = 1, …, *j* – 1, we use a generalized linear mixed effects model with form

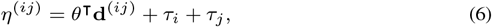

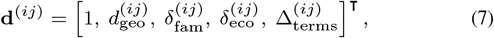

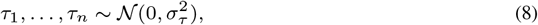

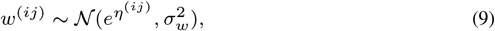

where *w*^(*ij*)^ is the Wasserstein distance between the inferred distributions of communicative need for languages *i* and *j*; 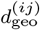 is their estimated geodesic distance (Haversine method) in standardized (normalized by standard deviation) units based on geographic coordinates in *glottolog* (and restricting to languages with small geographic extent); 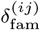 is a binary indicator of being in the same linguistic family or not (1 or 0, respectively); 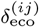 is a binary indicator of being in the same ecoregion or not (1 or 0, respectively); and 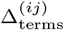 is the difference in their number of color terms, which we include as a control. The random effects *τ*_1_, …, *τ_n_* model the dependence structure of the pairwise measurements. Model diagnostics suggest reasonable behavior of residuals using a log-link function (SI Fig. C7). Fitted coefficients indicate a positive increase in dissimilarity with geodesic distance, and a decrease in dissimilarity with ecoregion, but no significant effect of shared language family (SI Fig. C7). GLMM fits were performed in R (v3.6.3) using the `lme4` (v1.1-21)package, with MLPE structure based on code from `resistanceGA` [70]. Model diagnostics based on simulated residuals were done using package `DHARMa` (v0.2.6).

Pseudo-*R*^2^ measuring overall model fit was computed as 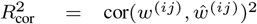, where 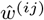 is the model predicted value for *w*^(*ij*)^, based on Zheng & Agresti [71]. For our model, 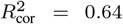. However, there is no standard, single measure of *R*^2^ for models with mixed effects. A recent proposal [72, 73] suggests reporting two separate quantities, a conditional and marginal *R*^2^, which can be interpreted as measuring the variance explained by both fixed and random effects combined 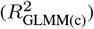, and the variance explained by fixed effects alone 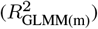. For our model we computed these as 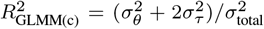 and 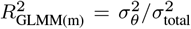, respectively, where 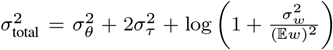 based on Nakagawa et al. [73]. For our model, conditional 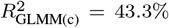 and marginal 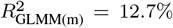. We based the inclusion of fixed effects on AIC (Fig. 5c) following best practices for MLPE models [74].

### Data availability

All data came from pre-existing datasets. Color vocabulary data was sourced from the World Color Survey online repository (http://www1.icsi.berkeley.edu/wcs/data.html). Additional language data was sourced through glottolog v3.4, available online (https://glottolog.org/meta/downloads). Data on biogeographic regions were provided by the World Wildlife Foundation, available online (https://www.worldwildlife.org/publications/terrestrial-ecoregions-of-the-world). Fruit reflectance data came from the Cambridge database of natural spectra, available online (http://vision.psychol.cam.ac.uk/spectra). Salient object data originating from the Microsoft Research Asia (MSRA) dataset are not publicly available, but were kindly provided to us on request by the corresponding authors of Gibson et al. [3]. Data generated by our inference method and used to estimate the average communicative needs across languages (Fig. 2a) and language specific communicative needs (SI Fig. SI E11–E27) will be shared under a creative commons license and made available on github.

### Code availability

Custom code was developed to infer communicative needs using the algorithm derived in this paper (SI Sec. B). All code will be made available as open source and shared via github.

## Supplementary Information

### A Rate-distortion theory

Rate-distortion theory [1, 2] provides a mathematical treatment of the problem of lossy compression, based on information-theoretic quantities. In information theory [1], the entropy of a discrete random variable, *X*, defined

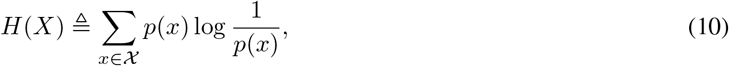

provides a measure of the average length of the shortest description (“amount of information”) needed to specify the outcome of random variable *X* with outcomes in the set 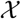 occurring with probability *p*(*x*). The joint entropy of *X* and a second random variable, *Y*, *H*(*X*, *Y*), is defined similarly in terms of the joint distribution of *X* and *Y*, *p*(*x*, *y*), and measures the average length of the shortest description needed to specify the outcomes of both random variables together. When the outcome of *X* is related to the outcome of *Y* in some (possibly nonlinear and stochastic) way, then the shortest description of both *X* and *Y* together may be smaller than the shortest descriptions of each of *X* and *Y* separately. In general, *H*(*X, Y*) ≤ *H*(*X*) + *H*(*Y*), with equality if and only if *X* and *Y* are statistically independent. The mutual information between *X* and *Y*, defined,

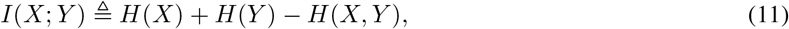

then gives a non-negative measure of the average amount of information *X* and *Y* contain about each other, which is nonzero if and only if *X* and *Y* are not independent.

In the lossy-compression context, for a given source (random variable) *X* and a description of that source, 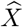, the mutual information 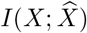 measures the amount of information the description contains about *X*, and it is this quantity we wish to minimize for compression, subject to a loss function, i.e. a measure of distortion. This can be formalized as

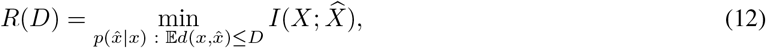

where the loss is measured in terms of an expected distortion, 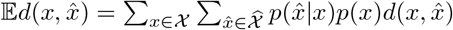, with *p*(*x*) a property of the source, and 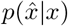 the mapping of *x* to 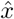 chosen to achieve on average the smallest description size possible, *R*(*D*), for a given allowable average distortion, *D*. Intuitively, the minimum compressed description size, *R*(*D*), increases as the allowable average distortion, *D*, decreases, dependent on the details of the source, *X*, and loss function, *d*.

#### A.1 Bregman clustering

The classical formulation of the rate-distortion tradeoff gives an optimal mapping of *X* to 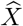 for fixed 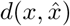. When every *x* and 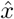 has coordinates in a vector space, denoted x and 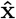, respectively, then for a large family of distortion measures known as Bregman divergences, optimal coordinates for each 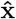 can be found [3] in addition to the optimal mapping between *X* and 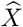. For Bregman divergence 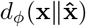, defined

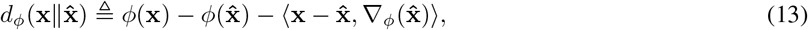

with convex function *ϕ*, gradient 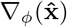 evaluated at 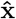, and inner product denoted 〈·, ·〉, the centroid of the mapping from 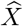 to *X* is the minimizer of the average distortion for 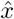, i.e.

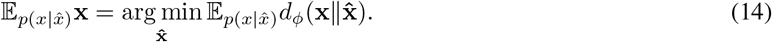

Solutions to rate-distortion Bregman clustering (RDBC) problems have the property that each 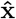 satisfies Eq. 14.

#### A.2 Compression model of color naming

The first RDBC model of color naming appears in work by Yendrikhovskij [4]. Using a perceptual measure of distortion, Yendrikhovskij [4] worked to show that efficient solutions to a tradeoff between average perceptual distortion and vocabulary size account for color categories based on natural image statistics. While the results are likely sensitive to the exact, unreported choices of natural images used to produce the image statistics [5], the conceptual link to a rate-distortion tradeoff has proved significantly productive. Using essentially the same RDBC-based compression model but disregarding scene statistics, instead using the neurophysiological constraints of perceptual discrimination and gamut alone, Regier et al. [6] showed that the compression model of color naming can qualitatively explain many of the typical vocabularies of natural languages in the WCS. Subsequent work by Zaslavsky et al. [7] investigated a “soft” partitioning variant of this same conceptual framework (although with additionally a mixture of Gaussians based measure of distortion derived from a Bayesian listener model of color naming), allowing for uncertainty in the mapping between terms and colors. In all cases, implicitly or explicitly, we can equivalently restate these compression-based accounts of color naming in terms of RDBC.

#### A.3 Focal colors as category centroids

In the World Color Survey, participants were asked to identify among the WCS color chips the “best example” of each basic color term identified in their vocabulary. In the WCS instructions to scientists conducting the field work, this is intended to elicit a response in the participant that identifies a color chip that “…is a good, typical, or ideal…”^†^ example of a given color term. In this work, we hypothesize that focal colors are observations of the centroids defined by Eq. 14. Two objections to this hypothesis immediately arise.

First, past work has shown that empirical measurements of category centroids differ from focal point positions [8–10], which would seem to invalidate our hypothesis. However, the discrepancy can be resolved by understanding how past work measured category centroids. Sturges and Whitfield [9], following earlier work by Boynton and Olson [8], conducted a color naming experiment similar to the WCS but in controlled laboratory conditions (and for English speakers only). Similar to the WCS, participants were asked to name, one by one in randomized sequence, a presented color chip, recording both the response as well as the timing of the response. The chips with shortest response times were considered the focal colors, and despite the difference in method these appear to be in good agreement with the “best example” focal colors recorded by Berlin & Kay for English speakers.

For each participant, the centroid of a category was computed as the average of all the color chips (in a given color space) that the participant named with that category’s color term (e.g. “red,” “green,” etc.). To write this out mathematically, we have a sequence of participant responses, 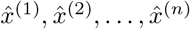, where each response is a color term, i.e. 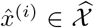, elicited by an experimenter presented color chip, *x*^(1)^, *x*^(2)^, …, *x*^(*n*)^, where 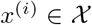. Note that each color chip in 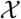 was presented more than once in the sequence of *n* presentations. Then the centroid for category 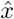 was computed as

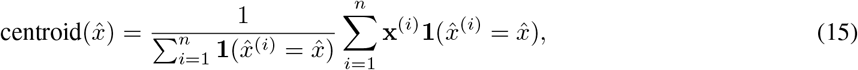

where **1**(·) is the indicator function equal to 1 if its argument is true and 0 otherwise, and x^(*i*)^ gives the coordinates of color *x*^(*i*)^ in color space. Let 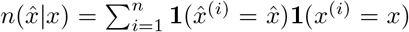 count the occurrences of 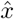 given presentation of color chip *x*, then we have

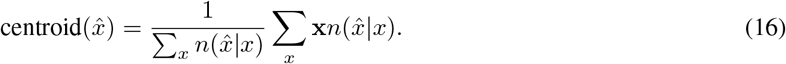

Let 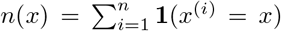 count the occurrences of *x* in the sequence. Then 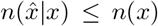, and 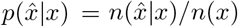 gives the fraction of times 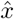 was used to name *x*, out of a total of *n*(*x*) occurrences. Since each color chip was presented the same number of times, we have further that *n*(*x*) = *m*. Then we have equivalently

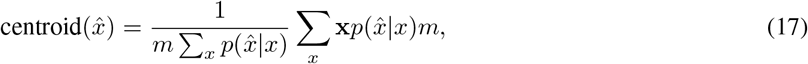

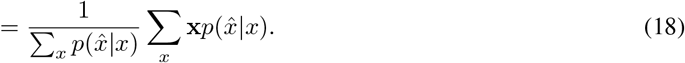

Lastly, note that 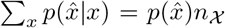, where 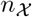 is the total number of color chips used, i.e. the cardinality of 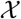, and 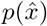 is the fraction of occurrences of 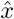 in the sequence. Thus we have

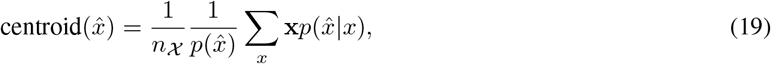

which by Bayes rule is equivalent to our definition of centroid with a uniform distribution of communicative need over the color chips, i.e. 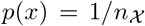. Thus in past work centroids have been shown to differ from focal colors *when a uniform distribution of communicative need over color chips is assumed*. In this paper, by contrast, we show that by inferring and using a non-uniform distribution of communicative need we better predict both empirical color term maps and focal point positions, and that focal point positions coincide with category centroids under this non-uniform distribution of needs.

The second objection stems from work done by Abbott et al. [11] investigating a measure of the “representativeness” of focal colors based on color category extents for the WCS. Representative colors of a given category are not necessarily those with the highest likelihood, i.e. maximizing 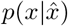, but instead are the most likely relative to their likelihood given any other category, weighted by the prior of that category, i.e. maximizing 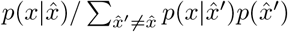. This appears problematic for the hypothesis that category centroids are equivalent to focal points, due to the bijection between Bregman divergences and regular exponential family distributions, and the equivalence between Bregman divergence minimization and maximum likelihood estimation [3, 12]. Again, it is crucial to examine definitions to see that the discrepancy is resolved by the assumption placed on the form of 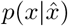. In Abbott et al. [11] 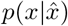 was assumed to be normally distributed. Whereas under the compression hypothesis, the maximum likelihood is taken over the mixture model as a whole, and the form of 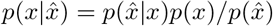 is not normally distributed in general. The broader message of Abbott et al. [11] is that focal color positions reflect a balance between typicality within a category and distinction from other categories; and this interpretation agrees with our identification of focal colors as category centroids when category centroids “compete” to represent different parts of color space, as in the compression model of color naming.

### B Inverse inference of source distribution

In this section we address the general problem of inferring an unknown source distribution, *p*(*x*), from knowledge of its compressed representation (i.e. a representation 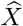 that lies on the rate-distortion curve for some unknown value of the tradeoff parameter, *β*). Concretely, we wish to find the *q*(*x*) that best approximates the unknown distribution *p*(*x*) using only what we know about 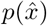 and 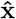 from its compressed representation, with no other assumptions. For fixed marginal distribution 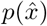 over 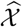, this can naturally be expressed as a problem of finding the conditional distributions 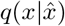 that together maximize the entropy of the marginal distribution 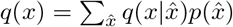 over 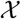, subject to a set of constraints that enforces we recover the known compressed representation, i.e.

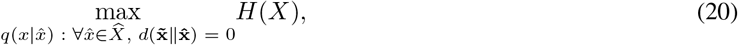

where *H*(*X*) is the Shannon entropy of *X* and 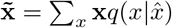.

We show that a numerical solution to this problem can be found via an alternating minimization strategy used by Blahut and Arimoto in their solutions to the channel maximization and rate-distortion problems [13, 14] and later generalized by Csiszár & Tusnády [15]. To do so, we first note that the objective function can be rewritten as

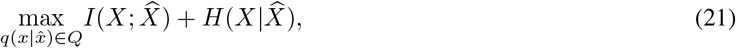

using the fact that 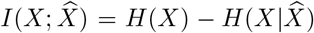. Here *Q* is the set of all conditional probability distributions such that 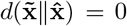 for all 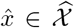. Since the mutual information term can be written as a maximization over 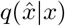, [13, 14] i.e.

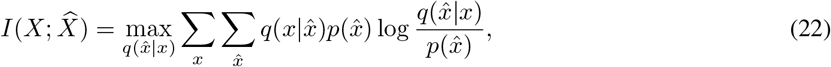

and 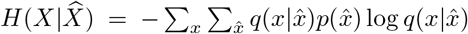 is constant with respect to varying 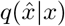, we can rewrite our objective function as a double maximization of the function

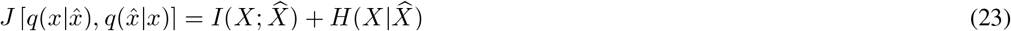

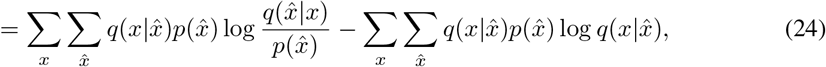

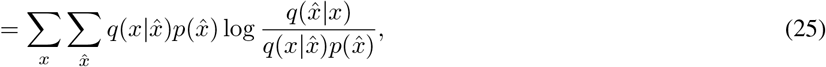

to change the problem into one of alternating maximizations over 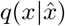 and 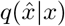, i.e.

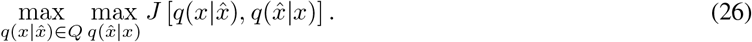

The inner maximization over 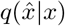 for constant 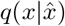 is given by 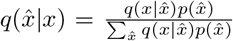, as previously shown by Blahut and Arimoto. The outer maximization over 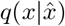 must respect a set of constraints that ensure we recover 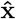 as a minimum distortion representation of x and that we have a valid probability distribution, i.e.

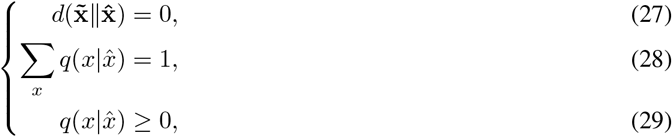

where 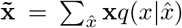. Eq. 27 enforces that there is no difference between the true compressed representation centroids 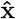 and those generated by the estimated 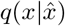, while the remaining two constraints ensure that 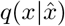 is a proper probability distribution.

Temporarily setting aside the non-negativity constraint (it will be enforced by the form of the solution), the Lagrangian is then

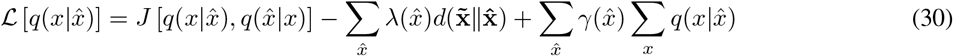

for fixed 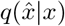. Taking the derivative with respect to 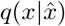 and setting equal to zero, we have

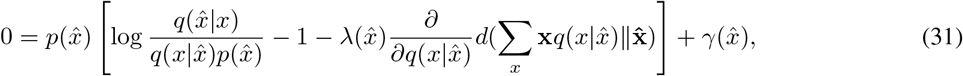

where we absorb a 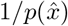 term into each Lagrange multiplier 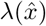. If the function *d* is a Bregman divergence, i.e. it can be written as 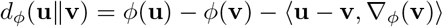 for some convex function *ϕ*, then

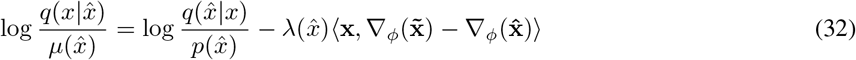

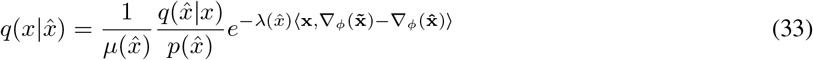

Where 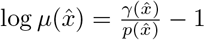.

For the constraint 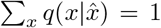 to be true, the Lagrange multipliers, 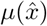, must act as a normalization factor, giving us

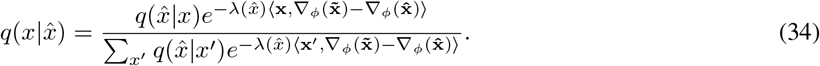

This also satisfies the non-negativity constraint for each 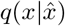, since 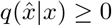, and *e^x^* ≥ 0 for any 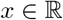. Finally, we can combine the unknown scalar 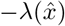 and vector 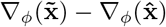 into a single unknown vector 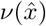, giving

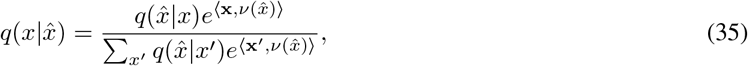

where 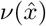 must be chosen such that Eq. 27 is true.

For any Bregman divergence, *d_ϕ_*(**u**‖**v**) = 0 iff **u** = **v** (see Banerjee et al. [3]). Thus to enforce Eq. 27, we need to find 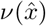 s.t. 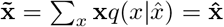. Let 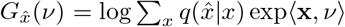. Then the vector of partial derivatives of 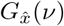 with respect to *ν* are given by

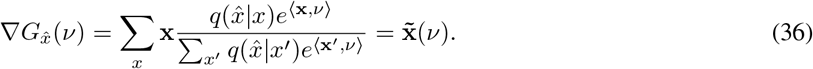

Since 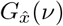 is strictly convex, we have by Legendre transform its convex conjugate dual,

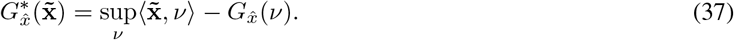

and vector of partial derivatives

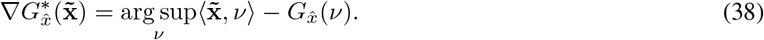

By the strict convexity of 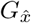 and the definition of the Legendre transform we have that 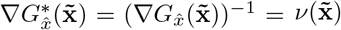, i.e. the unique choice of *ν* for a given value of 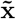. The unique choice of *ν* to guarantee 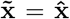 is then simply 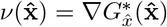, which can be computed numerically via e.g. BFGS.

The alternating maximization algorithm is then to iterate

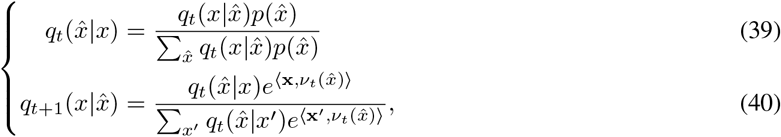

with 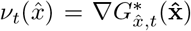, and starting from any initial 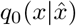. By construction, the choice of 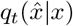 maximizes *J* for fixed 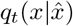, and 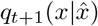 maximizes *J* for fixed 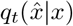, subject to their respective constraints. We thus have a sequence indexed by *t* of non-decreasing values for *J*, which converges whenever the maximum entropy is finite. The solution for the marginal distribution of *X* is then given by 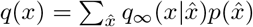.

#### B.1 Convergence to the global optimum

In this section we will show that the alternating minimization algorithm defined by Eq. 39 and Eq. 40 converges to the global maximum of 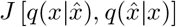 for any initial choice of 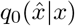. We will do this using a geometric approach developed by Csiszár & Tusnády [15]^†^, which for example can be used to prove convergence to the global optimum for the alternating minimization algorithm proposed by Blahut [14] to find numerical solutions to the rate-distortion problem. First, note that maximizing 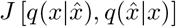 is equivalent to minimizing 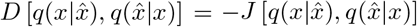. Then by Theorems 1 and 2 of Csiszár & Tusnády [15], to show convergence to the global minimum via alternating minimizations of *D* it is sufficient to show that the “three points property” and “four points property” both hold for *D* and a choice of functional, *δ*.

##### Definition 1.

(From Csiszár & Tusnády [15]) Let *δ*[*p, p*′] be a non-negative valued function on *P* × *P* such that *δ*[*p, p*] = 0 for each *p* ∈ *P*. Given *D* and *δ*, for a *p* ∈ *P* the three points property holds if

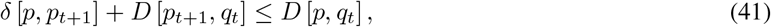

whenever *p*_*t*+1_ = arg min *_p_ D* [*p, q_t_*]. The four points property holds for a *p* ∈ *P* if for every *q* ∈ *Q*

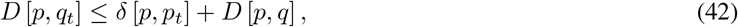

whenever *q_t_* = arg min *_q_ D* [*p_t_, q*].

We will show that the three and four point properties hold for *D* and the following choice of *δ*,

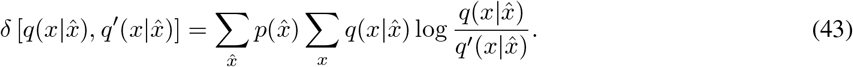

Non-negativity of Eq. 43 follows directly from the non-negativity of the KL-divergence and 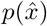, as does equality holding iff 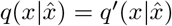.

We will also make use of the fact that we can rewrite both the definition of *δ* given by Eq. 43 and *D* in terms of the following Bregman divergence,

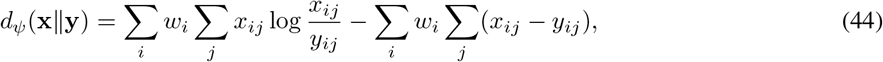

where *w_i_* are constant non-negative weights that sum to one, and *x_ij_, y_ij_* ≥ 0, not necessarily summing to one. In this case *ψ* is the strictly convex function 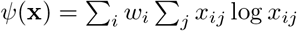. Then with 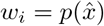, *i* indexing elements of 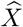, and *j* indexing elements of *X*, we have that

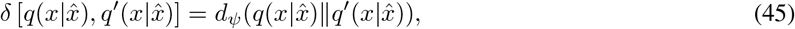

and

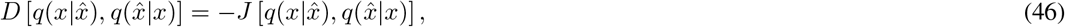

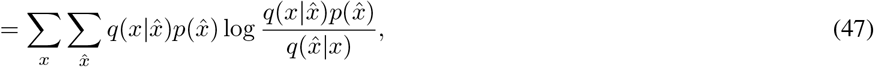

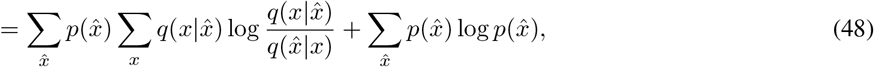

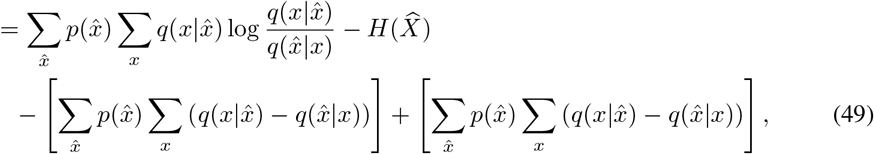

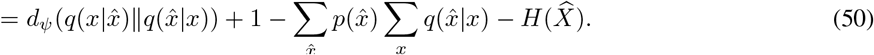

##### Lemma 1.

The three points property, 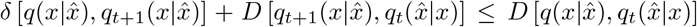, where 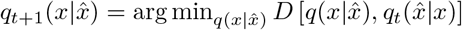, holds.

*Proof*. Rewriting using Eq. 45 and Eq. 50, we must show that

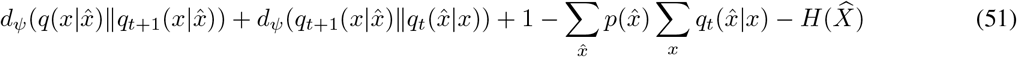

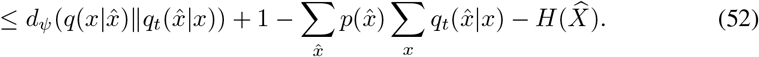

Cancelling, we need to show

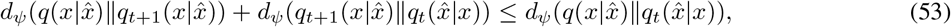

which follows immediately from the Generalized Pythagoras Theorem [3] and the fact that by construction solutions of Eq. 40 maximize *J* for fixed 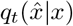, so that

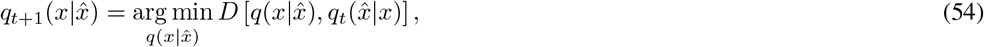

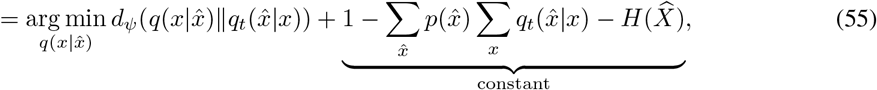

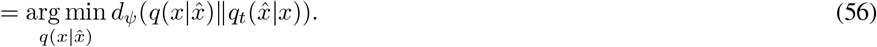

##### Lemma 2.

The four points property, 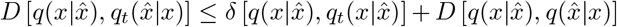, where 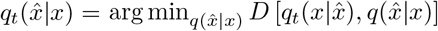, holds.

*Proof*. From the definitions of *D* and *δ*, we must show that

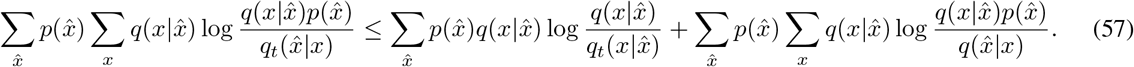

By subtraction, equivalently we must show that

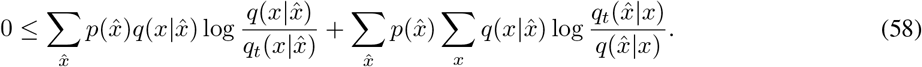

Denoting 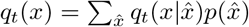, from Eq. 39 we have that 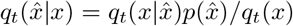. Then by substitution we have

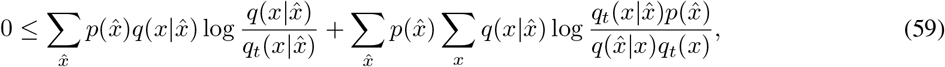

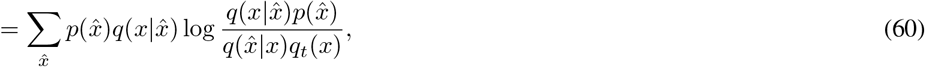

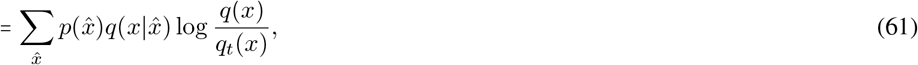

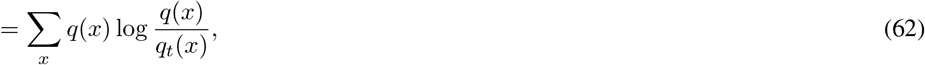

where Eq. 61 follows from the fact that 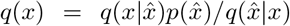, and Eq. 62 from the fact that 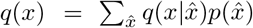. Then this is equivalent to the statement that 0 ≤ *D*_KL_ [*q*(*x*)‖*q_t_*(*x*)], which is true by non-negativity of the KL-divergence.

##### Theorem 1.

The sequence of alternating maximizations defined by Eq. 39 and Eq. 40 converges to the global maximum of 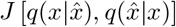 for any initial choice of 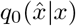.

*Proof*. Proof of Theorem B.1 follows from satisfying the five point property of Csiszár & Tusnády [15], which is implied by satisfying the three and four points properties from Lemma 1 and Lemma 2, respectively.

#### B.2 Uniqueness

In the previous section we showed that the solution found by the alternating maximization algorithm is globally optimal. Here we show that the optimal *q*(*x*) distribution is also unique.

##### Theorem 2.

The distribution 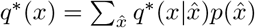 for the 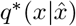 achieving the maximum of 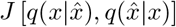 is unique.

*Proof*. Assume *q*\*(*x*) is not unique, and there exists a distinct solution *q*′(*x*) that also achieves the maximum of 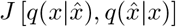 with 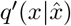. Then two things must be true.

First, since *q*\*(*x*) and *q*′(*x*) are distinct, then 0 < *D*_KL_ [*q*\*(*x*)‖*q*′(*x*)]. From the definition of the KL-divergence and using the fact that 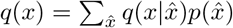, we have that

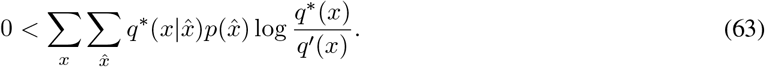

Since 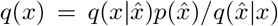 (for any choice of 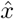), the definition 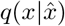 from Eq. 40, and the equivalence of 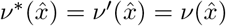, we have

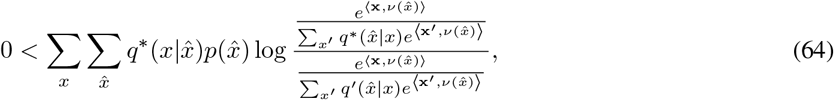

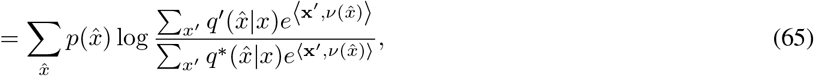

since after cancellation none of the terms depend on *x* except 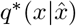, and 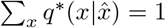.

Second, since both *q*\*(*x*) and *q*′(*x*) achieve the global optimum, we must have that 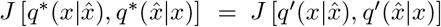. Then after cancelling we have

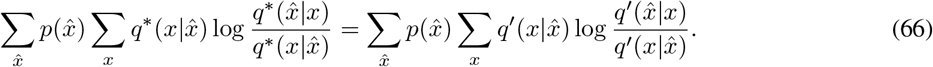

From the definition of 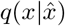 in Eq. 40 and the equivalence of 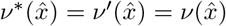,

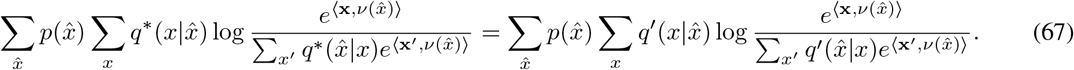

Then, since 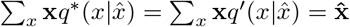, we can cancel the 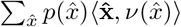 term from both sides, and using the fact that 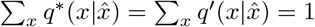, we have

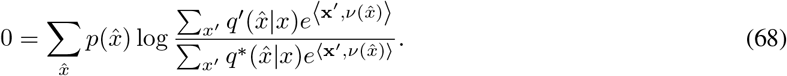

But this contradicts the inequality established by Eq. 65. Thus *q*\* (*x*) must be unique.

#### B.3 Example inference and comparison to prior work

As an illustrative example, we present the results of the inverse inference method above for a known distribution of needs, *p*(*x*). This toy example allows us to study the properties of the inverse inference when the ground truth, *p*(*x*), is known. We also use this example to illustrate the difference between our inference method and inferences based on two prior methods in the literature. Rather than solving for the maximum entropy distribution consistent with a rate-distortion optimal vocabulary, the “capacity achieving prior” (CAP) method [7] assumes instead that the true *p*(*x*) will be one such that, given a vocabulary of term mappings 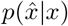, we only ever need to communicate the *x*’s that are maximally unambiguous to specify with that vocabulary. The CAP distribution is the one that achieves the maximum channel capacity for the given term map 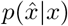 (the specification of the channel from *X* to 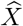), i.e. satisfying

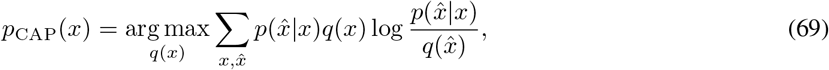

where 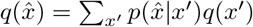. This is a strong assumption in general, and when it is violated, as we will see in this section, the CAP provides a poor approximation of the true distribution *p*(*x*).

We also compare our approach to the word-frequency method (here abbreviated WF) proposed in [17], which asks for the maximum entropy distribution *p*_WF_(*x*) that satisfies the linear constraints 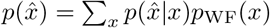. In other words, the WF method solves for the distribution of communicative needs consistent with a given term mapping, 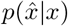, and known word frequencies, 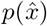. To understand what this does in practice, it is instructive to consider the case of “hard” clusters where 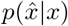 equals either 1 or 0. In this case, we derive an analytical solution for the WF inference (see SI Sec. D.1), which is given by 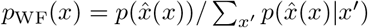, where 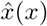 is the unique nonzero 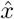 in 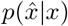; i.e., the 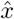 chosen for a given *x*. In effect, then, the WF method approximates communicative need by dividing the frequency of a given word, 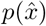, uniformly across its mapped domain. In the “soft” case, solutions behave similarly but with an additional factor accounting for “fuzzy” boundaries between 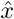 domains. While this gives considerably more reasonable estimates of needs than the CAP approach, it depends on the availability of word frequencies, 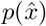, and it again requires knowledge of the term maps 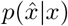; the former is unknown for almost all WCS languages, and the latter introduces circularity when aiming to predict term maps based on language-specific communicative needs.

In our toy example *x* ∈ *X* covers the unit grid (*n* = 100 × 100) with an arbitrary but specified distribution *p*(*x*), as shown in Fig. B1a (ground truth). The figure also shows the RDBC solutions for either 4 or 8 terms (Fig. B1a and B1b, respectively; this example uses squared Euclidean distance as the distortion measure). The ground truth distribution *p*(*x*) was chosen to be nonuniform, with a broad probability gradient from (0, 0) to (1, 1), and a smaller-scale low to high to low to high oscillation in probability along the x-axis. The RDBC centroids and Voronoi (nearest-centroid) regions show a non-uniform division of *X* into clusters (or “terms,” to link this to the terminology of the color naming problem), as a result of using a non-uniform *p*(*x*).

Based on only the positions of the focal terms 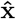 and the term frequencies 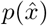, our inverse method produces an estimate of *p*(*x*) that recapitulates the broad-scale features of the ground truth. The inverse inference performs well even with as few as 4 terms (Fig. B1a), with some additional, fine-scale details captured when inferring from 8 terms (Fig. B1b). By contrast, the distributions inferred by the CAP method, which are not based on 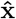 and 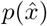 but instead require knowing the full term map 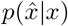, deviate significantly from the ground truth (Fig. B1a and B1b; note different scale).

In Fig. B1c, the entropy of inferred and CAP solutions are shown for a broader range of vocabulary sizes (from 2 to 10). The figure also quantifies the dissimilarity between the ground truth and the estimated distributions, based on their KL-divergence. Successive iterations of the inverse inference algorithm show monotonic convergence to a maximal entropy value that lies between the ground-truth entropy and the unconstrained maximum entropy distribution (uniform over *X*). Note there are only small differences between the maximum entropy values achieved when varying the vocabulary size used (the equivalent of the number of color terms). While not directly constrained by the inverse inference method, since the ground truth distribution is assumed unknown, the inverse method converges to distributions that are very close to the true distribution. Solutions become closer to ground truth as the vocabulary size increases, but even small vocabularies provide inferences that closely approximate the ground truth. By comparison, CAP solutions have entropies that are substantially lower than the maximum or even the ground truth entropy, and they are sensitive to vocabulary size. CAP solutions are orders of magnitude more divergent from ground truth, compared to the results of the inverse inference method we have developed.

### C Application to color categories

We use the inverse inference method of SI Sec. B to find the distributions of communicative need for empirical color vocabularies via the following correspondence (outlined in Fig. 1c). In this application, the source, *X*, denotes the visible colors that need to be communicated, which are the WCS stimuli set. Each WCS stimulus color, *x*, in the set of WCS stimuli, 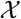, has a position x in CIE Lab, a perceptually uniform color space. The unknown distribution of communicative need we wish to infer is *p*(*x*). Our estimate of *p*(*x*) will be the one that best matches the known position, 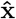, of each “best-example,” or focal, color for each term, 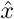, in the language’s color vocabulary, 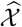, and is otherwise maximally unbiased (maximizes the entropy of the inferred distribution).

**Figure B1.**
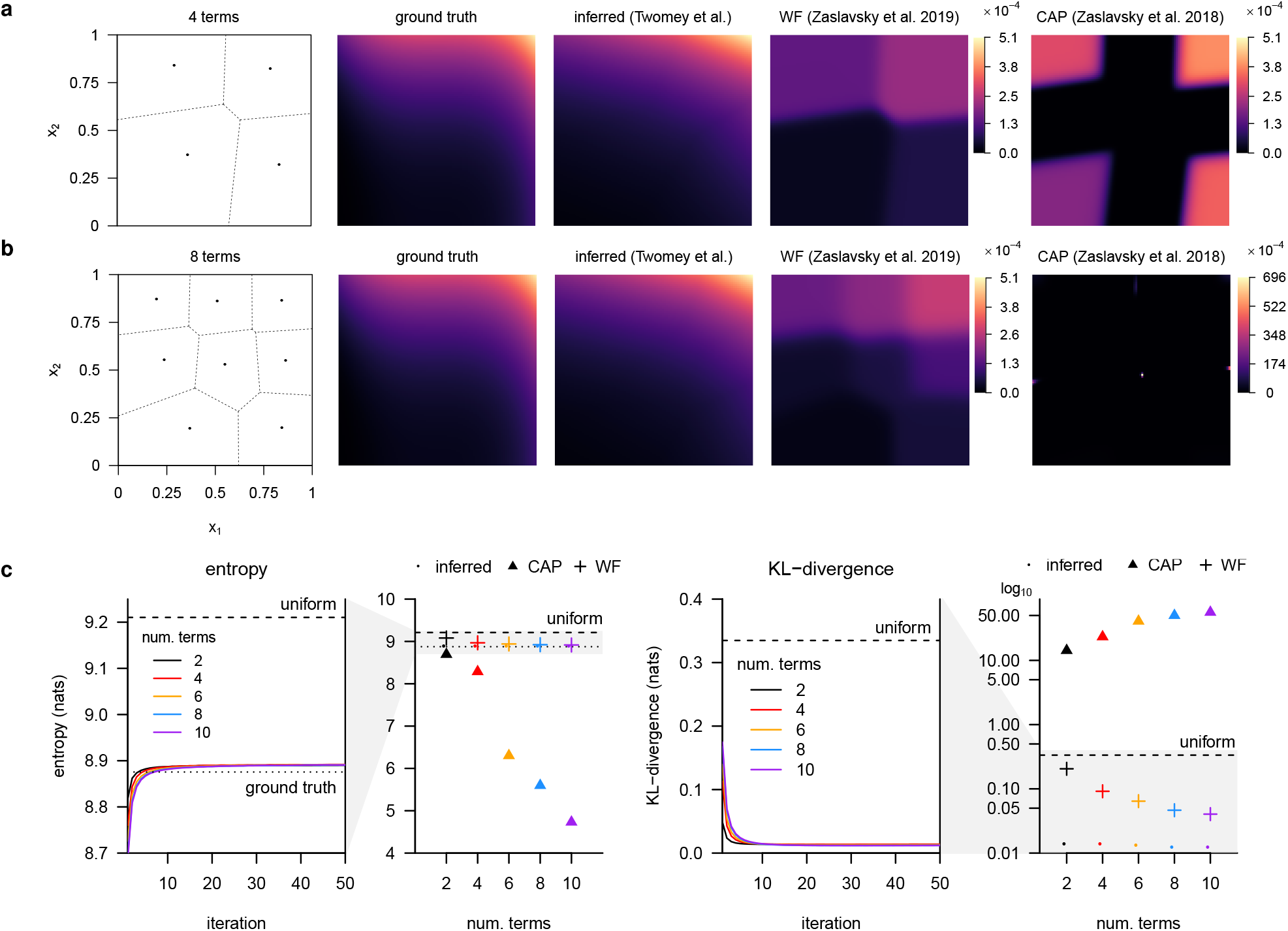
Example inference of the distribution, *p*(*x*), underlying a given rate-distortion solution. **(a)** An example rate-distortion vocabulary (*left*) with 4 terms (black points show position of term centroids, 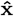) giving a compressed representation of a dense square grid (100 × 100) of elements *X* with distribution *p*(*x*) (*middle-left* “ground truth”). Using the positions, 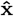, and probabilities, 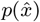, of the vocabulary terms alone, the inverse algorithm infers distribution *p*_inf_(*x*) (*middle-center*), which approximates the main features of the true distribution *p*(*x*) (same color scale). For comparison, we also show the distributions inferred under the word-frequency (WF) method (*middle-right*) and the capacity achieving distribution (CAP, *right*). **(b)** The same example as in (a) but for an 8-term rate-distortion vocabulary (grid and ground truth distributions are identical). **(c)** Evolution of the key quantities in the inference algorithm’s iterative solutions to the example used in (a) & (b), for vocabulary sizes 2, 4, 8, and 16, with comparisons to WF and CAP used in prior work. (*Left*) The entropy of the inferred distribution monotonically increases with each iteration of the inverse inference algorithm, and converges to values between the true entropy (dotted line labeled “ground truth”) and the maximum entropy (dashed line labeled “uniform”) for this example as expected. The entropy is reduced and approaches the true entropy as the number of terms is increased, but only to a small degree compared to CAP. The adjacent figure with expanded *y*-axis shows converged values for inferred, WF, and CAP distributions (circles, crosses, and triangle plotting symbols, respectively). CAP solutions have lower entropy than the true distribution, are more sensitive to the number of vocabulary terms, and become increasingly different as the number of terms increases. (*Right*) The KL-divergence between the inferred distribution and true distribution tends to decrease and converges to small values. This is a consequence of matching the term centroids since the true distribution is not known to the inverse inference algorithm. The adjacent figure compares the inferred, WF, and CAP distributions KL-divergence to the true distribution at convergence (log scale). The distributions inferred by our method are close to the true distribution, and become even closer with increasing vocabulary size; while the CAP distributions are far from the ground truth and become increasingly farther, even more so than uniform. The WF solutions are sensitive to the number of terms available, and at 10 terms give solutions that are nearly a factor 3 further from ground truth than our inferred solutions for 2 terms (0.04 vs. 0.014).

Intuitively, in the inverse inference procedure (SI Sec. B), the vectors 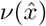 can be thought of as “pulling” on the inferred distribution such that the inferred centroids match the position of the true centroids. In the example shown in SI Sec. B.3, the positions of the true centroids lie in the interior of the boundary of all the *x* positions. To match prior work and the WCS itself, we use the WCS color chips (Fig. 1a) as the support set for the inverse inference. Since WCS participants selected focal colors from this same set, the average focal color position across participants could lie on or near the boundary of the support set if there is high agreement among participants. To match these positions with the given support set, the inverse method would be forced to “pull” with overly large magnitudes towards these remote points, when this is just an artifact of the constraints on participants and the choice of support.

To check if this was the case, and to mitigate any impact it may have, we constrained the maximum magnitude that any 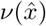 could have, and varied this value as a parameter, λ. At λ = 0 the inverse method makes no attempt to match the language centroids, and we recover only the uniform distribution over the WCS color chips. At λ = ∞, we recover the unconstrained inverse inference method. At intermediate values of λ, pathologically large magnitudes have limited impact on the inference. If indeed there are pathologically large magnitudes at play, then there should be a large difference between the entropy at λ = ∞, where the inferred distribution becomes overly concentrated at the problematic focal point, and at intermediate values of λ for which the nearly the same RMSE between inferred and true focal points is achieved. Fig. C2a shows that this is exactly the case, and suggests that λ ≤ 0.25 is sufficient to achieve RMSE’s close to the unconstrained solutions, while maintaining substantially higher entropies.

Note that RMSE is measured using the empirical focal points and the position of the focal points for the optimal rate-distortion fit using the inferred distribution at a given value of λ. Rate-distortion solutions were found using the standard alternating minimization algorithm (see Banerjee et al. [3]),

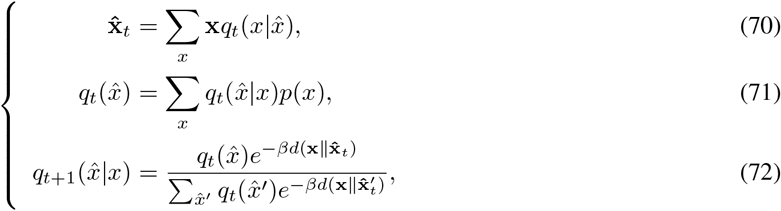

where 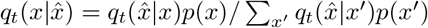, and *β* is a parameter that acts as an “inverse temperature,” controlling the “softness” (low values of *β*) or “hardness” (high values) of the boundaries between terms given by 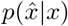. Since RDBC solutions are not unique, we run the algorithm starting from many different initial conditions (initial 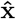 positions drawn uniformly at random from the set of WCS color chips) until convergence (change in 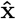 positions between iterations is < 1 × 10^−5^ or the maximum number of iterations is reached; max iterations 1 × 10^4^ used in searches for the optimal value of *β*; max iterations 5 × 10^4^ for calculation of RDBC solution using the optimal value *β*), and keep the solution with lowest mean squared error. We used a standard derivative-free nonlinear optimization method (bound optimization by quadratic approximation [18], via the `nloptr` (v1.2.1) package for R v3.6.3) to search for lowest mean squared error values of *β*.

For each B&K+WCS language, the minimal RMSE for inverse inference with λ ≤ 0.25 is shown in Fig. C2b (y-axis), and compared with the minimal RMSE (same optimization procedure for non-unique RDBC solutions and choice of *β*) for uniform (x-axis). In all cases, use of the inferred distribution reduces RMSE compared to uniform (all points below 1–1 line). As useful references, we quantified the RMSE for within-language variability in focal point positions among participants (via bootstrap resampling of participant responses and measuring their RMSE with respect to the mean focal point positions for that language), as well as the RMSE when all terms are off by one WCS color chip. Most inferred distribution RMSE’s are between the median values of these two reference quantities, which is not the case for uniform.

Similarly, in Fig. C2c we show the absolute improvement in term map predictions for the WCS languages shown in Fig. 3b, comparing the Earth mover’s distance (EMD) between predicted and empirical term maps based on inferred (y-axis) and uniform (x-axis) distributions. WCS languages were used for term map comparisons both for the ability to resample from among speaker responses (the B&K data surveyed only one speaker per language) to assess confidence intervals on improvement in Fig. 3b, and because the B&K study design substantially differed methodologically from the WCS in the color naming task.^†^ In the WCS color naming was assayed for each color chip, whereas in B&K participants selected chips out of the full set of stimuli [19]. While the B&K term maps are related to 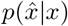, they are not straightforward estimates of 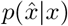 as in the WCS, and behave qualitatively differently. As useful reference points, we computed the EMD between empirical vocabularies and rotations thereof, approximated by cycling WCS columns 2:41. This transform preserves the structure of each vocabulary while increasing the displacement (in hue) between the true and rotated terms, and has been used in prior work on color naming [6]. Here it provides a more meaningful distance scale for the EMD measurements than e.g. chip-wise randomization.

**Figure C2.**
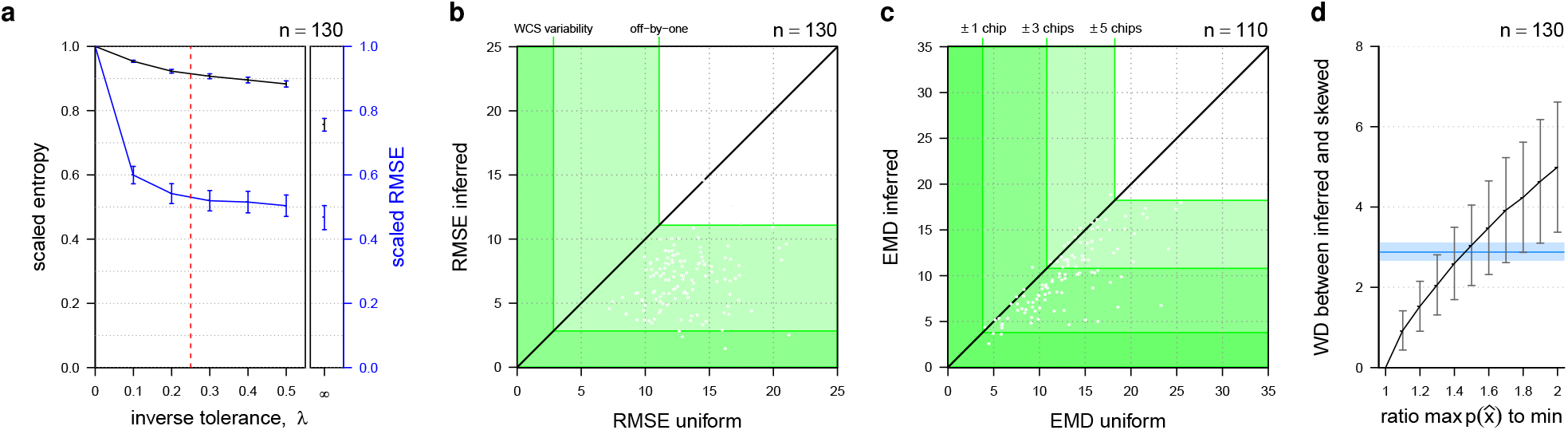
Application of inverse inference algorithm to WCS. **(a)** In the constrained-maximum version of the inverse inference method, small values of the inverse tolerance parameter, λ, (x-axis) can achieve values of RMSE (mean and 95% confidence intervals shown scaled relative to uniform) comparable to the non-relaxed inverse inference (λ = ∞), while maintaining a much higher entropy (shown scaled relative to uniform). **(b)** RMSE between rate-distortion optimal vocabularies under inferred distribution, *p*_inf_(*x*), (y-axis) and empirical ground truth, compared to RMSE under uniform distirbution, *p*_unif_(*x*), (x-axis). All points lie below 1-1 line (black), showing that inferred strictly improves over uniform for matching focal point posisitions. Regions bounded by reference lines for median RMSE from within-language variability in focal point position (“WCS variability”) and median all focal point positions off-by-one WCS chip (“off-by-one”) are shown overlapping in green. **(c)** Average Earth mover’s distance (EMD) between rate-distortion predicted and empirically observed term maps for each WCS language vocabulary under a uniform distribution of communicative need (x-axis) or the language inferred distribution (y-axis). Reference lines at ±1, ±3, and ±5 chips show the median EMD across languages comparing empirically observed languages to themselves ± a rotation in hue (rotation of WCS columns 1:40). **(d)** Sensitivity of inferred language-specific communicative needs to the assumption of uniform term frequencies, 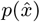. Mean and standard deviation Wasserstein distance is shown between inferred distributions under a uniform 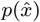 and an asymmetric (“skewed”) distribution constructed with varying ratios of max 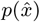 to min 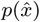 (x-axis). Reference line (blue) shows median Wasserstein distance and 95% CI between inferred distributions derived from language mean focal color position and focal colors resampled from language speaker responses (based on WCS languages).

#### C.1 RMSE reference points

We provide three points of comparison for the RMSE distributions shown in Fig. 2b. First, the “WCS variability” reference line was computed by resampling participant focal point choices by language, recomputing the mean focal point across resampled participants, and measuring the RMSE between the recomputed focal points and the actual language focal points. We used the median computed RMSE as a useful reference point approximating a lower bound on how well predicted focal points might be expected to perform. Second, the “off-by-one” reference line was computed by repeatedly offsetting each focal point by one WCS chip sampled uniformly at random from the neighborhood of WCS color chips in Fig. 1a and measuring the RMSE between the set of perturbed focal points for a language and the actual focal points. The median computed RMSE in this case gives an intermediate point of comparison for predicted focal point RMSE distributions. Third, the “random” reference line was computed by resampling each language’s focal points from the WCS color chips uniformly at random without replacement, then assigning each resampled focal point to the nearest true focal point, and measuring the RMSE of the two sets of focal points under this assignment. This gives an approximate upper bound on how poorly a predicted set of focal points might perform, using the same procedure for assigning predicted focal points under the rate-distortion model to actual language focal points.

#### C.2 Sensitivity of inferred distributions to term frequencies, 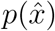

The inverse inference algorithm uses the frequency of vocabulary terms, 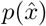, as part of the inference process for determining *p*(*x*). This information is not available for color vocabularies in the B&K and WCS datasets, and further field work would be required to estimate these quantities directly (with the additional caveat that vocabularies may have changed since having been originally surveyed). And so the present work uses the simplifying assumption of a uniform 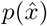. This is a reasonable approximation, given the WCS selection criteria for basic color terms and evidence in English that basic color terms are elicited with approximately equal frequency under a free naming task [10]. Nevertheless, to investigate the sensitivity of inferred distributions to this choice, we compared inferred distributions under increasingly asymmetric (“skewed”) distributions for 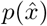, sampled from a linearly increasing set of probabilities between the minimum and maximum 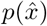. Fig. C2d shows the Wasserstein distance between the inferred distributions under uniform 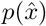 and the skewed distribution as a function of the ratio between the maximum and minimum 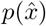. As a useful reference point, we computed the median Wasserstein distance between inferred distributions under the uniform 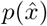 assumption re-sampled from the WCS language speaker populations. Ratios in usage greater than approximately 1.5 would be needed before non-uniformity would begin to have a more significant impact on inferred distributions than the among-speaker variability inherent in the data. While this suggests the choice of a uniform 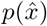 is reasonable to a first approximation, the extent to which this assumption of uniformity may be violated in some languages remains an open, but potentially tractable, question for future field work.

#### C.3 Sensitivity of inferred distributions to vocabulary size, 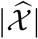

Under the rate distortion hypothesis, color vocabularies optimize the information a listener can infer about the color being referenced, based on the color term chosen by a speaker. Because there are far fewer terms than perceivable colors, there is by necessity some loss of information caused by the compression of colors into terms. As a result, the size of a vocabulary (number of terms) should have some impact on our ability to infer the underlying distribution of communicative needs, *p*(*x*): larger vocabularies should provide more resolution and more detail in the inferred distribution. For the B&K+WCS languages, we can expect that fewer terms will result in the recovery of only broad-scale features of a language’s communicative needs, while more terms allow for additional detail.

This effect is demonstrated in Figure C3a. Here we generated rate-distortion efficient vocabularies for simulated languages, each generated from the same underlying distribution of underlying communicative needs and differing only in the number of color terms. We then inferred the distribution of needs from the simulated focal color positions, using our inverse inference method. As expected, we find that having more color terms allows for more detail to be recovered in the inferred distribution of needs, although the results are qualitatively similar across a range of vocabulary sizes.

We also investigated the relationship between vocabulary size and inferred needs in more systematic detail. To do so, we again generated rate-distortion efficient vocabularies for pairs of languages sharing the same underlying communicative needs and differing only in vocabulary size. We used the B&K+WCS average inferred distribution as a “ground truth,” and the number of terms in each simulated vocabulary was restricted to the range of terms found in the B&K+WCS data. We then inferred the communicative needs for each simulated vocabulary in the pair, and we measured their Wasserstein distance. Figure C3b shows a small but statistically significant impact of differences in vocabulary size on the measured distance between inferred distributions of need – which arises because vocabulary size has an impact on the resolution of the inference. For comparison to these simulations, in which the underlying needs are kept constant, we also plotted the distances between inferred needs measured in the empirical data, for all pairs of B&K+WCS languages. The empirical distances between inferred needs are much larger than can be explained by the simulated data. These results imply that differences in vocabulary size alone cannot explain the large differences observed among B&K+WCS inferred communicative needs. Moreover, the relationship between differences in vocabulary size and differences in inferred needs has substantially greater magnitude in the empirical data than in the simulated languages. This suggests that there may be typical ways in which communicative needs evolve as the vocabularies of languages change in size – which is an interesting hypothesis for future study.

#### C.4 Capacity achieving distributions for individual WCS languages

The capacity achieving distributions, which are referred to as priors (CAP) in the literature, should not in general be expected to approximate the true distribution of communicative need, as shown in SI Sec. B.3. Here we reproduce the average CAP across the WCS languages reported in Zaslavsky et al. [7, 17]. The average CAP differs by several orders of magnitude from the average distribution *p*(*x*) inferred in this paper (Fig. C4a). The language-specific CAP’s for Waorani and Martu-Wangka are shown in Fig. C4b: they each differ radically from the communicative needs we estimate by our inference method. The CAP distributions feature implausible variation in communicative need across nearby colors.

**Figure C3.**
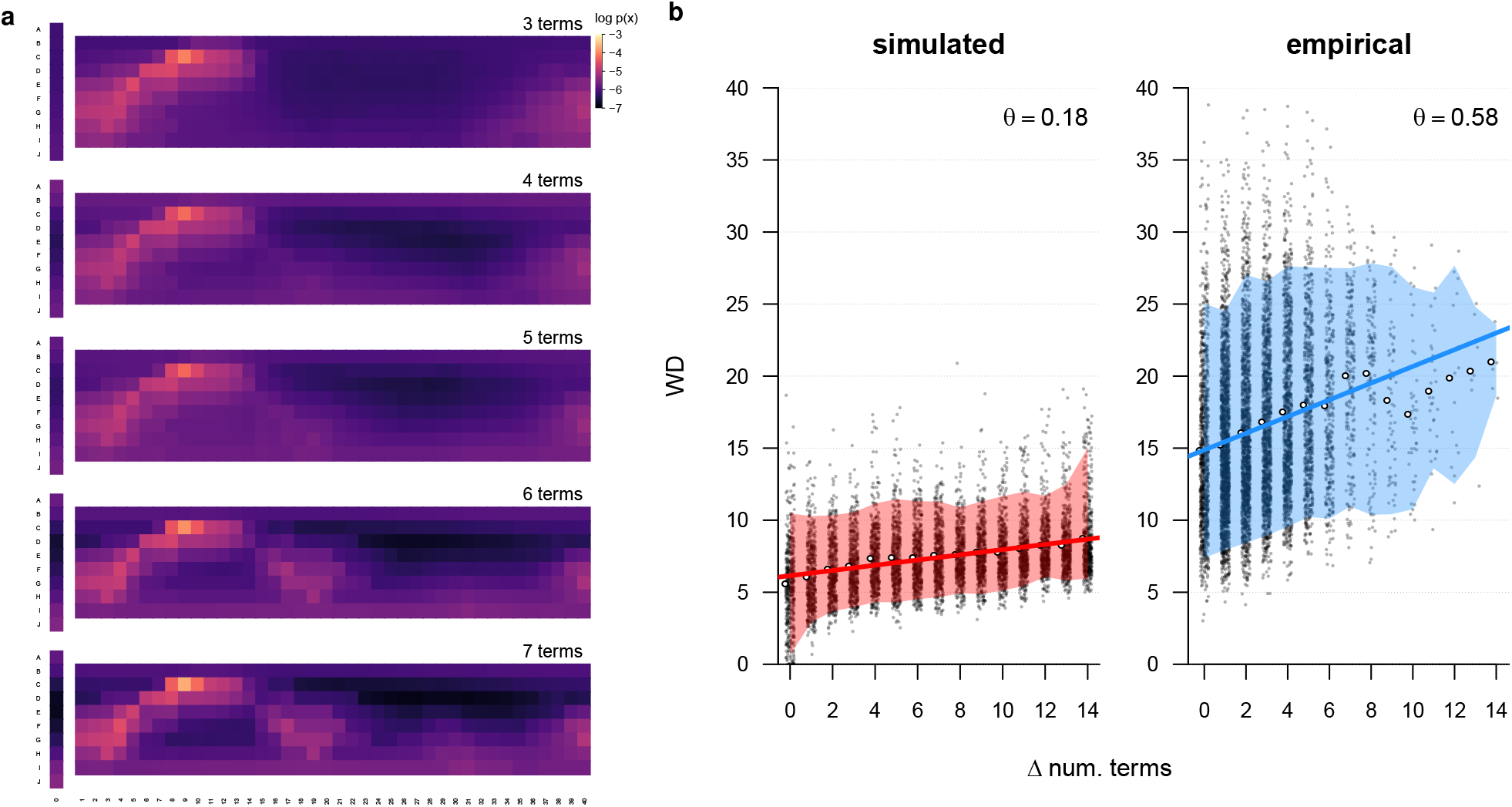
Sensitivity to number of color terms. **(a)** Average inferred distributions of communicative need for rate-distortion optimal vocabularies simulated with different numbers of terms but the same underlying communicative need (the B&K+WCS average inferred distribution). A larger number of terms provides more resolution and detail in the inferred distribution of needs, but the inferred distributions are nonetheless qualitatively the same. **(b)** Simulated (*left*) and empirical (*right*) Wasserstein distances (WD) between inferred distributions of need for pairs of languages, shown as a function of the difference in the number of their terms (Δ num. terms, white points show mean WD for each Δ num. terms). Differences in inferred communicative needs (WD) are substantially smaller in the simulations, which isolate the effects attributable to vocabulary size alone, compared to the differences observed among empirical languages (red and blue bands show 90% extent of simulated and empirical distances, respectively). Also, the relationship between vocabulary size and differences in inferred needs (WD) is substantially smaller (slope of linear regression *θ* = 0.18, red line) in the simulated data with a single shared distribution of needs, compared to the relationship observed in the empirical data (slope *θ* = 0.58, blue line).

#### C.5 Field work variability

Variability in how the field work for the WCS was conducted for different languages does not appear to explain the instances of non-improvement in Fig. 3b term map predictions. For the WCS, native speakers were asked to use only the basic color terms of their language, as previously identified according to a set of specific linguistic criteria. However in some cases it seems that native speakers apparently were not so constrained, either by experimenter or participant choice. Based on the identification of these two modes in the WCS by Gibson et al. [20] in their supplementary materials, there was no apparent relationship between the choice of methodology and a language showing improvement or no improvement under the inferred distribution vs. uniform.

#### C.6 Potential correlates of communicative needs

Here we consider possible correlates of the communicative needs we infer, based on proxy measures and potential confounds suggested by prior work.

**Figure C4.**
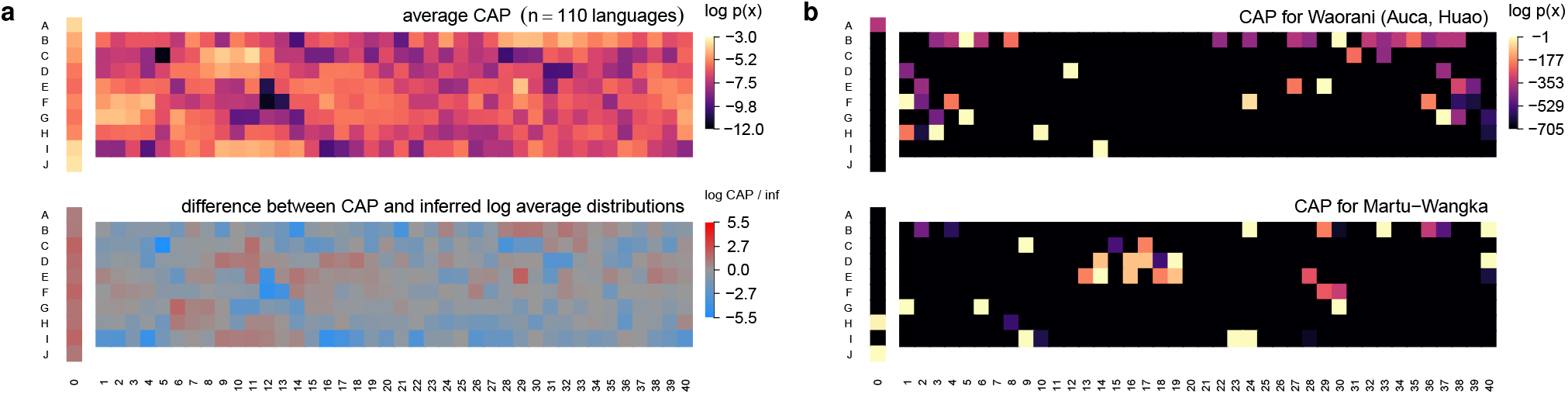
Comparison to the capacity maximizing distributions (“capacity achieving priors”) for WCS languages. **(a)** (Top) Approximate replication of Fig. 3a from Zaslavsky et al. [17] showing capacity achieving prior (CAP) averaged across WCS languages (here we include all WCS languages; some were excluded in Zaslavsky et al. [17]). (Bottom) The difference between the average CAP and average prior we infer (see Fig. 2a) ranges over several orders of magnitude (log scale). **(b)** CAP distributions for two languages used as examples in Fig. 5a (note different scale). Under the CAP inference, two neighboring Munsell color chips may exhibit a 10^300^-fold difference in communicative need.

**Figure C5.**
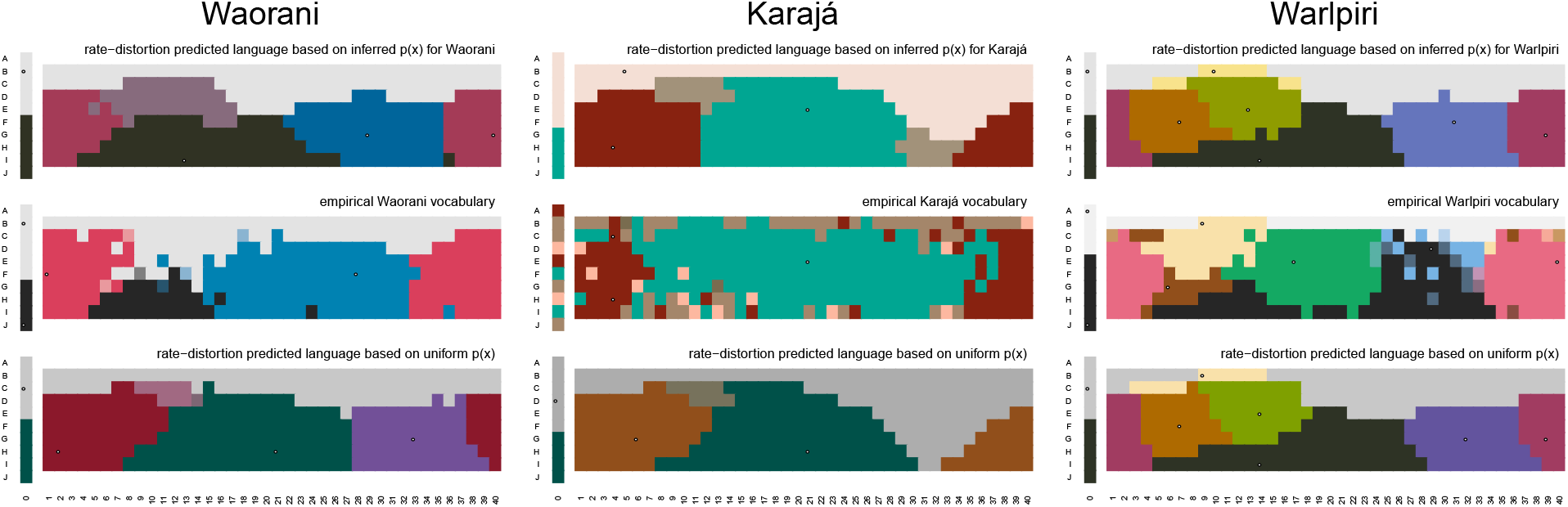
Results for WCS languages previously identified in the literature as possible outliers. RDBC results shown for the languages labeled in Fig. 3b (Pirahá shown in Fig. 2c). Prior work has hypothesized that Pirahá [21], Warlpiri [22], Waorani [23], and Karajá [23], may be exceptions in some way to the broad trends identified in the WCS. All but Warlpiri appear to be substantially improved when we account for language specific communicative needs.

**Figure C6.**
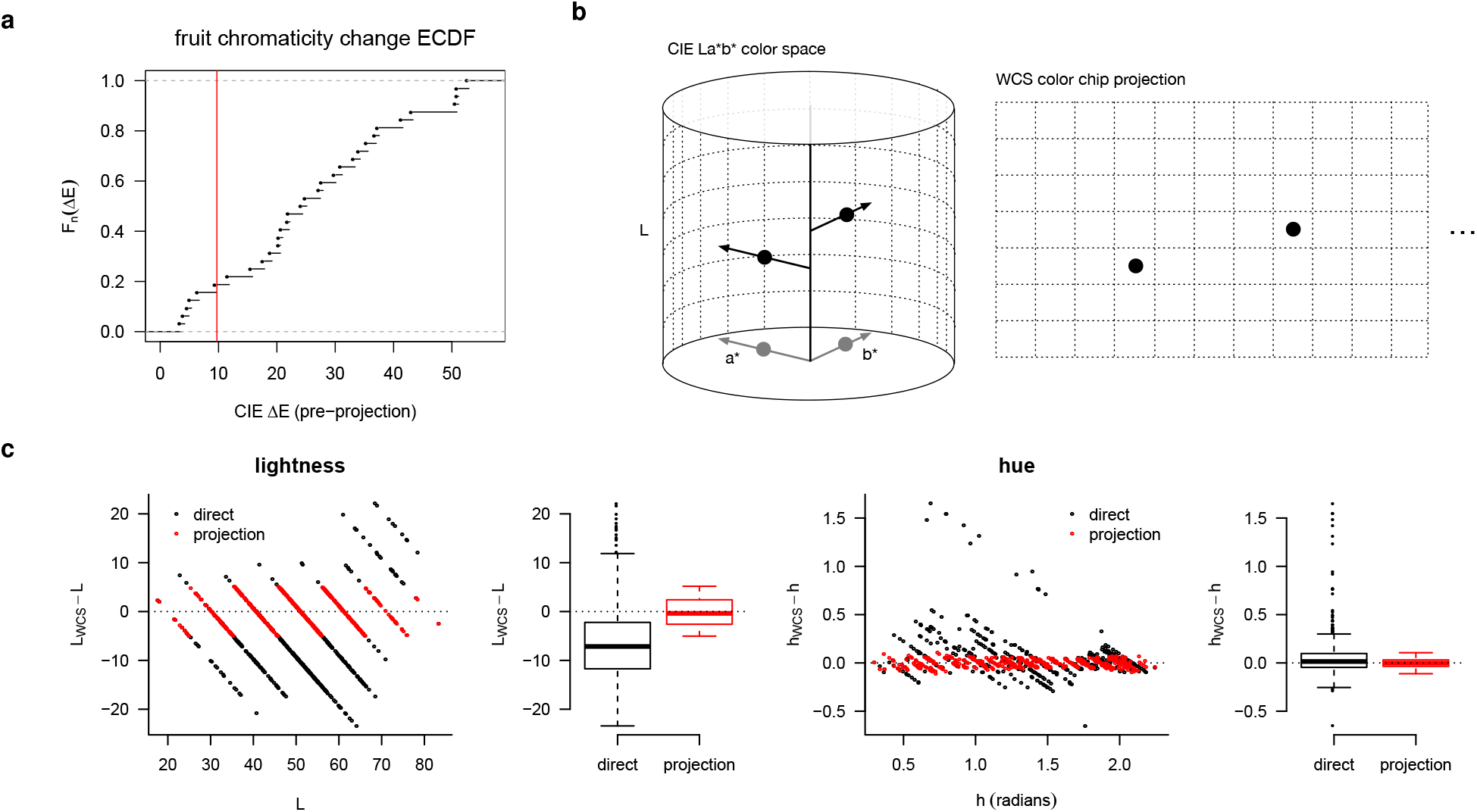
Treatment of Sumner & Mollon fruit chromaticity measurements. **(a)** Empirical cumulative distribution function (ECDF) of the change in fruit chromaticity between unripe and ripe states. Not all fruits signal ripeness by a change in chromaticity [24, 25]; other indicators may include size or smell. For each species collected by Sumner & Mollon having at least one measurement in each of the ‘unripe’ and ‘ripe’ classifications, the species’ chromaticity measurements is included in our analysis (Fig. 4c & d if the CIE Lab difference (ΔE^✶^) between the mean unripe and ripe measurements is greater than a threshold (red vertical line). This threshold is determined by the minimum Δ*E* of a subset of the species measurements for which we could establish a significant change in mean CIE Lab coordinates at the *p* < 0.01 level based on a Hotelling T^2^ test. **(b)** After conversion from spectral measurements to CIE Lab coordinates, the final step is to find the nearest WCS color chip in CIE Lab space. The WCS color chips form a high-saturation outer shell of the Munsell color array, privileging lightness (L) and hue angle over saturation. We adopt this same choice by selecting nearest neighbors based on L and hue angle (i.e. normalizing the (a^✶^, b^✶^) position sub-vector), ignoring saturation. **(c)** The choice of matching by projection rather than directly by ΔE^✶^better constrains the difference in lightness (L) and hue (h) between the matched WCS color chips and the true CIE Lab coordinates, with the tradeoff of a small increase in the overal mean ΔE^✶^(35.5 vs. 44.7). However this tradeoff appears to be necessary to make meaningful comparisons between fruit ripeness categories; without projection there is substantial variation in the residuals as a function of L and h. Box-plots show median and first and third quartiles; whiskers extend to the minimum (maximum) up to 1.5 times the interquartile range, with outliers shown as individual points.

**Figure C7.**
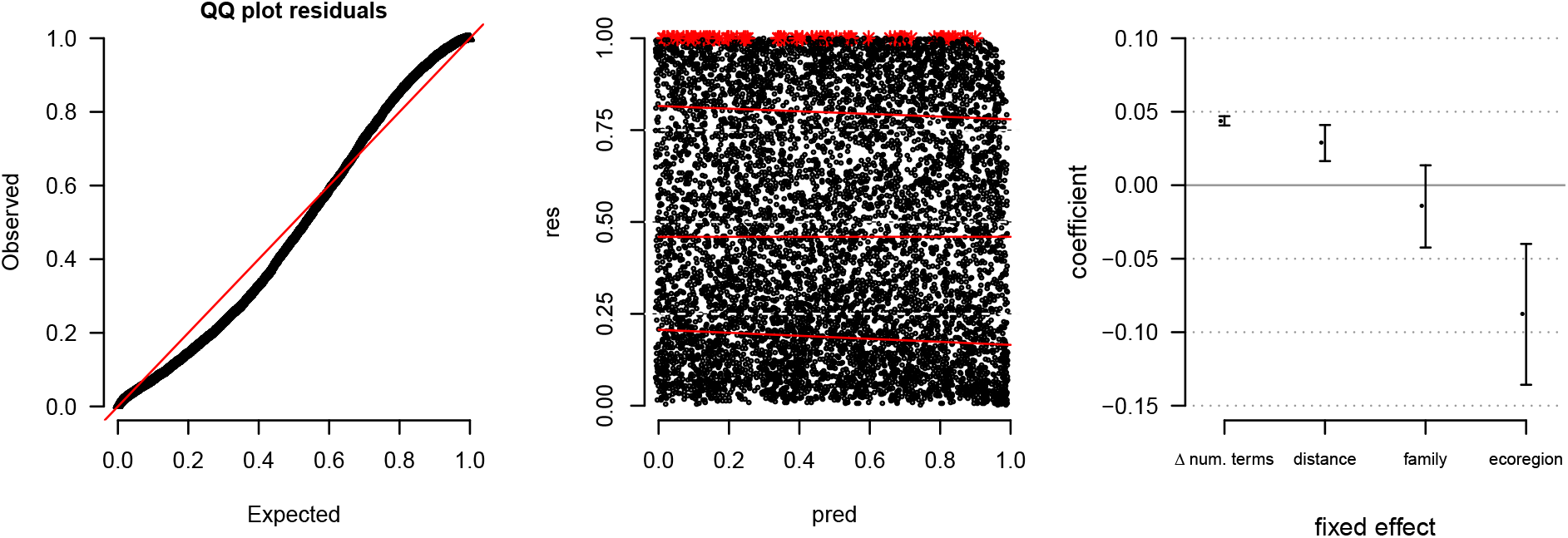
Diagnostics for GLMM of differences in communicative need between languages. (*Left*) Uniform quantile–quantile (QQ) plot of expected vs. observed GLMM model residuals. (*Middle*) Rank transformed model predicted values (pred) vs. residuals (res), with quantile regressions (red lines) compared to theoretical quantiles (dashed white lines at 0.25, 0.50, and 0.75); simulation outliers shown as red stars. (*Right*) Fixed effect coefficients and 95% confidence intervals for geodesic distance (Haversine method; standardized units), shared linguistic family (TRUE=1, FALSE=0), and shared ecoregion (TRUE=1, FALSE=0). Positive coefficients indicate an increase in dissimilarity (increase in Wasserstein distance), while negative coefficients indicate a decrease. Out of *n* = 125^2^ language pairs, 73% and 60% shared the same linguistic family or ecoregion, respectively. Variance inflation factors (VIFs) for distance, family, and ecoregion were 1.291, 1.219, 1.149, respectively. All VIFs are less than 5, showing low multicollinearity.

##### C.6.1 Communicative efficiency (surprisal)

Gibson et al. [20] sought to understand communicative needs by estimating communicative efficiency, or surprisal, defined as

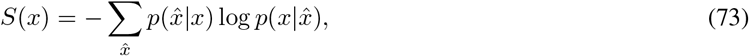

(Eq. 1 of Gibson et al. 2017), where *x* are color stimuli (i.e. the WCS color chips), 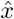 are color terms, and 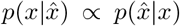 by Bayes rule under the assumption of a uniform *p*(*x*). This is distinct from our work, where by “communicative needs” we in fact mean the quantity *p*(*x*) – namely, the chance that a speaker needs to reference color *x* – which we infer directly for individual languages by maximum entropy (SI Sec. B). Nevertheless, we can ask whether surprisal *S*(*x*) is predictive of the language-specific communicative needs we infer. While we do find a moderate positive correlation (Pearson’s *ρ* = 0.41; *n* = 36,300; see SI Fig. C8a), the vast majority of variance in needs *p*(*x*) remains unexplained (1 – *R*^2^ = 0.83). The substantial differences between these two quantities explains why the needs that we infer have a comparatively weak correlation with the warm-cool trend in colors of salient objects, whereas the warm-cool trend is stronger for the surprisal measure.

##### C.6.2 Color saturation (chroma)

The WCS color chip stimuli were chosen to cover a range of Munsell lightness values and hues at approximately maximal “saturation” (Munsell chroma). Maximal color saturation, or chroma, thus varies as a function of Munsell lightness value and hue for these stimuli. Our inference of communicative needs depends on the positions of color chips used in the WCS. One concern, then, might be that the variation in color saturation, or chroma, in some way directly determines our inferred communicative needs. If this were the case, then we would expect to find a systematic relationship between inferred language-specific communicative needs and Munsell chroma. But we do not find any systematic relationship (SI Fig. C8b), even though the highest values of chroma tend to correspond to high values of *p*(*x*). This relationship for high chroma values may reflect the observation that participants choice of focal colors can be biased towards highly saturated colors [26]. Or it could alternatively, or additionally, reflect a common cause in the determinants of communicative needs and perceptual discrimination.

**Figure C8.**
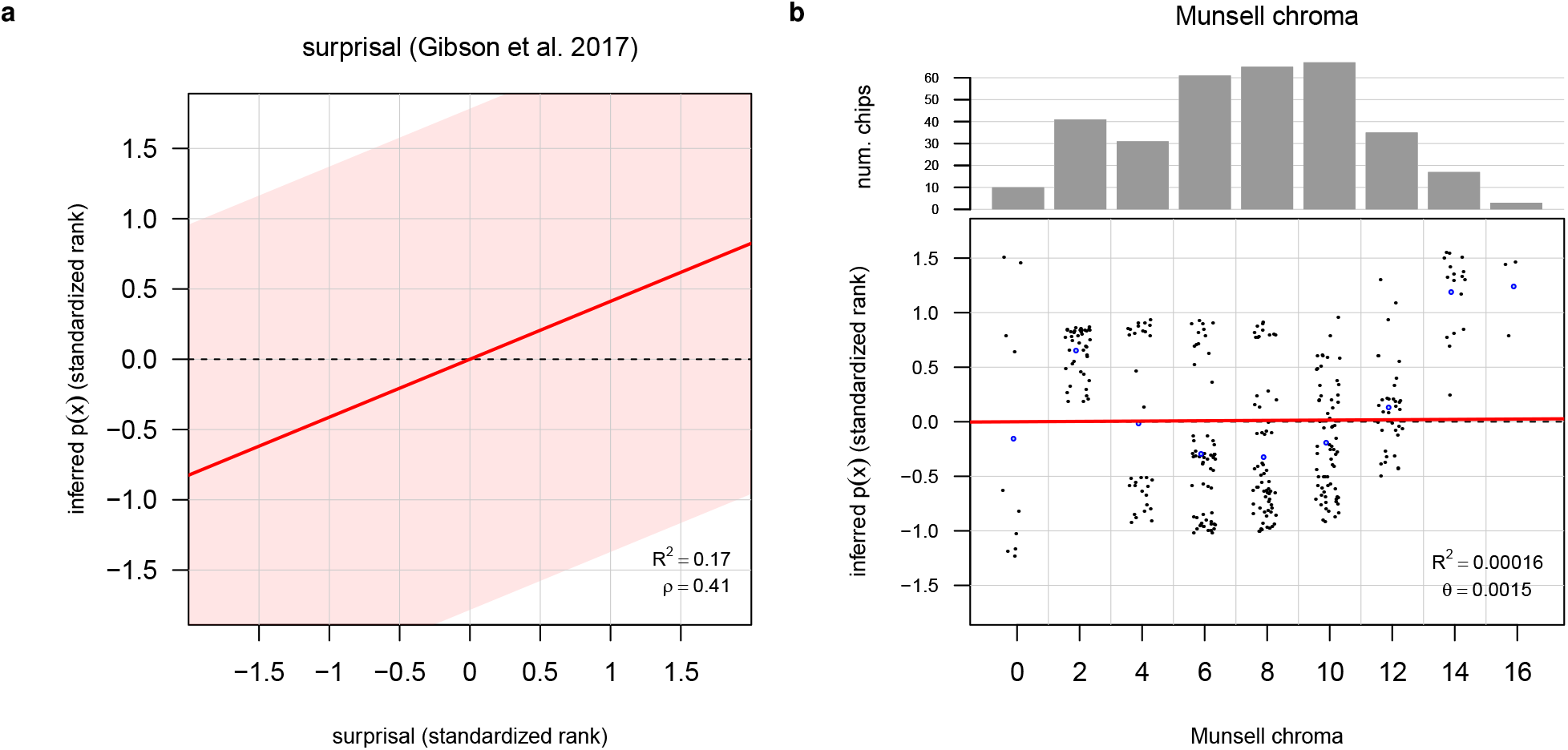
Relationship of inferred language-specific communicative needs to surprisal (Gibson et al. 2017) and color saturation of WCS stimuli (Munsell chroma). **(a)** Surprisal is moderately correlated, but not strongly predictive, of inferred communicative needs. Red line and legend show linear relationship between standardized rank values; light-red area indicates 95% prediction interval. Points show average (over languages) standardized rank inferred communicative needs (y-axis) compared with average (over languages) standardized rank surprisal (x-axis) for each WCS color chip. **(b)** Munsell chroma (color saturation) of WCS stimuli are not predictive of language-specific inferred communicative needs. Red line and legend describe linear relationship based on 110 languages and 330 color chips per language; *n* = 36,300 total comparisons. Points show the Munsell chroma (+ jitter; x-axis) and average (over languages) standardized rank inferred communicative needs for each WCS color chip (y-axis). Standardized ranks are computed per language. The total number of WCS color chips of each Munsell chroma used in the WCS is shown at top. Blue points show expected values of inferred communicative needs by Munsell chroma. Expected values are poor summaries for intermediate chroma values due to multi-modality.

### D Comparison to alternative approaches

In this section we provide additional discussion and comparison of our approach to inferring communicative needs with prior approaches. While past work proposed methods in the context of approximating a single, global distribution of need [7, 17, 27], it is reasonable to consider whether or not those approaches could also be used to estimate language-specific needs. First, for intuition, we provide a brief derivation of an analytical solution to Zaslavsky et al.’s word-frequency (WF) approach [17] for the special case of “hard” category boundaries, i.e. 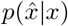 equal to either 1 or 0 only. Second, we give illustrative examples comparing our approach with that of WF [17] and CAP [7] for inference of communicative needs and prediction of color naming maps. Finally, we provide a systematic comparison of estimation methods for communicative needs when word frequencies are unknown (which is the case for almost all languages in the WCS).

#### D.1 Analytical solution for a special case of the WF method

The WF method [17] approximates communicative needs by finding the maximum entropy distribution over colors, *x*, such that the marginalization of the joint distribution formed by measured term maps, 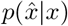, and estimated *p*_WF_(*x*), matches the measured word frequencies, 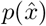; i.e., such that 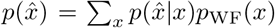. The method assumes that word frequencies 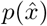 are known. By Lagrange multipliers, one can show that solutions (for “hard” or “soft” conditions), must have the form 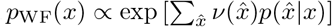, where the constant of proportionality normalizes *p*_WF_(*x*), and 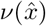 are Lagrange multipliers that need to be chosen to enforce the constraint that 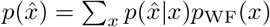.

Let *μ* be the constant of proportionality; in the case of hard clustering, let 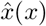 denote the one nonzero 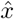 for a given *x*’s 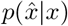 distribution; and with a slight abuse of notation, let 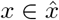 indicate the set of all *x* s.t. 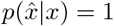. Then for hard clustering we can decompose the constraint into two parts,

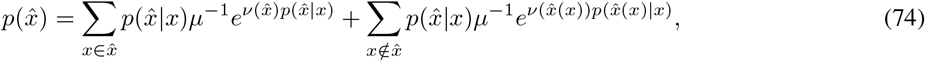

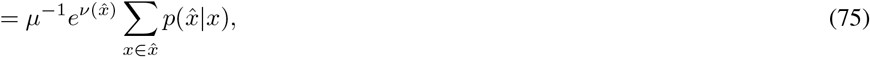

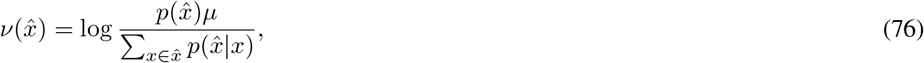

where the second step follows because (for hard clustering) 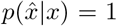 for any 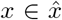 and = 0 for any 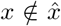. Then after plugging this into the form of the solution and cancelling out *μ*, we have

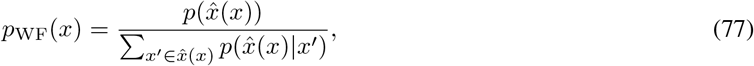

where the denominator simplifies to 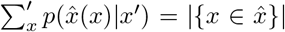. In words, the maximum entropy solution for hard clustering under the WF constraints simply apportions the frequency of a word, 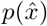, evenly across its domain as given by 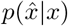. This analytical solution for the hard clustering case gives a helpful intuition for understanding the performance of WF more generally, as we describe in the subsequent two sections.

#### D.2 Illustrative comparison of approaches for language-specific needs

Here, we consider two illustrative examples of our inference method and model predictions in comparison to past work, where the ground truth is known. For each example, we generated a simple, arbitrary ground truth distribution of communicative needs over a unit square domain. (We consider a comparison across a more diverse set of systematically generated examples in the subsequent section, SI Section D.3). In the first example, shown in SI Fig. D9a, the true distribution of communicative needs has a maximum near the top right, and a minimum near the bottom left. The ground truth RDBC for this example provides just three categories (centroids and largest-likelihood 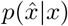 shown with unique colors). Without using knowledge of the term map, 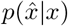, our inference method is able to recover the coarse features of the true distribution of needs. The WF method approximately evenly divides the frequency of each term, 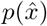, across its mapped domain, 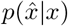, similar to the “hard” partitioning case solved analytically in SI Section D.1. The CAP method exponentially concentrates probability mass away from the boundaries between terms (all distributions in SI Fig. D9 are shown on a log scale).

**Figure D9.**
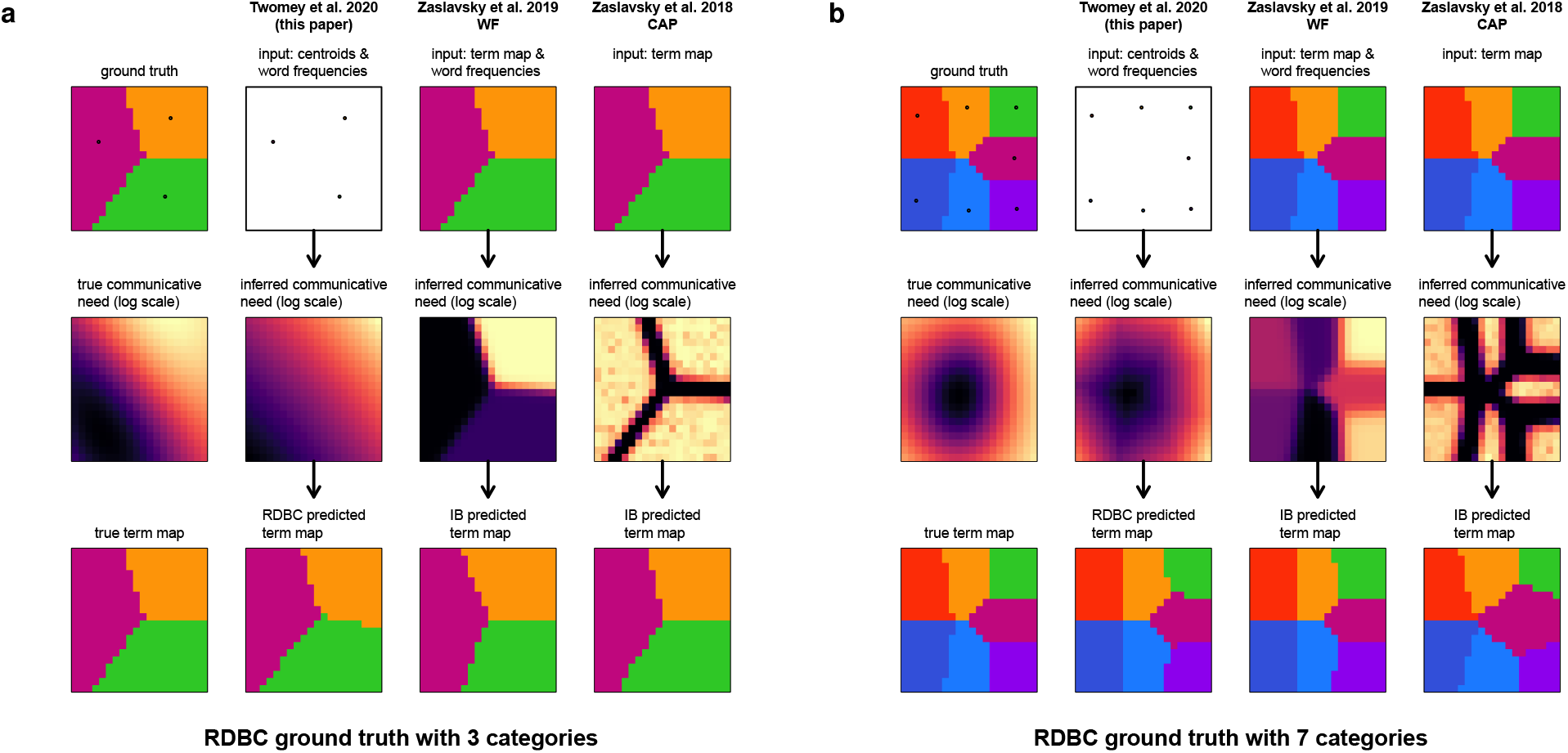
Comparison of methods for inference of communicative needs and prediction of term maps, when the ground truth is known. **(a)** A rate-distortion optimal “vocabulary” with three terms for the unit square is shown at top-left, with category centroids (points) and best-choice term maps (colored regions). The true distribution of communicative needs, *p*(*x*), is shown in the middle row (ground truth). Our inference method (second column) takes as input the category centroids and, if available, their frequencies, 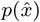, and produces an estimate of communicative need (middle row, same column). Our model of color naming then predicts term maps (bottom row, same column) using the inferred communicative need. Inferences based on the word-frequency based approach of Zaslavsky et al. [17] (third column), and the CAP approach of Zaslavsky et al. [7] (fourth column) do not accurately reconstruct the true distribution of needs. These two methods each use the IB model for prediction of term maps. Note that for language-specific communicative needs, term maps would necessarily be inputs for both the WF and CAP methods, leading to circularity in prediction of term maps. **(b)** A seven-term vocabulary example.

In these examples, the ground truth rate-distortion Bregman clustering (RDBC) is based on a squared-error measure of distortion. It then seems surprising that predictions of term maps using the CAP distribution, which does not well approximate the ground truth communicative need distribution, nonetheless closely resemble the true term mapping, 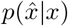, when using with the information bottleneck (IB) model proposed in Zaslavsky et al. [7]. Under the IB model, categories are not characterized by a centroid at a single point in space, but by a distribution over all points in space, which gives a large degree of additional flexibility. When coupled with language-specific inferences based on e.g. CAP, this evidently allows for recovery of the ground truth term maps despite the difference between the true generative process (based on centroids at single points in space with distortion measured by squared-error between points) in this example and the process specified by the IB model (based on mixtures of Gaussians over all points in space, with distortion measured by the KL-divergence between Gaussian mixtures). More critically, the requirement of empirical term maps, 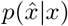, as inputs for both the CAP and WF approaches necessarily leads to circularity if these methods are then used for predicting language term maps based on language-specific inferences of communicative need.

#### D.3 Systematic comparison of approaches for language-specific needs

The examples shown in SI Fig. D9 and earlier in SI Fig. B1 illustrate the relative performance of our method, the WF approach, and CAP, under a few ideal test conditions. Next, we conduct a systematic investigation of these three methods controlling for the level of spatial detail in the ground truth distributions of communicative need, and withholding information about 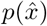, which is unknown for virtually all WCS languages, from all inference methods. Ground truth distributions were generated on a log scale from sinusoids with random phase and amplitude below a given spatial frequency. Distributions were scaled such that entropy was held constant across all generated examples for all spatial frequency cutoffs. This scaling fixes the KL-divergence between each generated ground truth distribution and uniformity at a constant, providing a consistent scale across examples.

SI Fig. D10 shows the KL-divergence between the distributions of communicative need recovered by each method and the generated ground truth distribution, as a percentage of the divergence between uniformity and ground truth, averaged over 500 examples at each spatial frequency cutoff. Our method (SI Fig. D10a) recovers the low-frequency features of the ground truth distribution even when 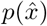 is unknown (assumed uniform) and in a manner that is relatively insensitive to the number of categories (number of terms) of the RDBC on which it is based. High-frequency information is lost, as expected, though recovery of low-frequency information across all spatial frequency cutoffs always improves estimates over uniform.

**Figure D10.**
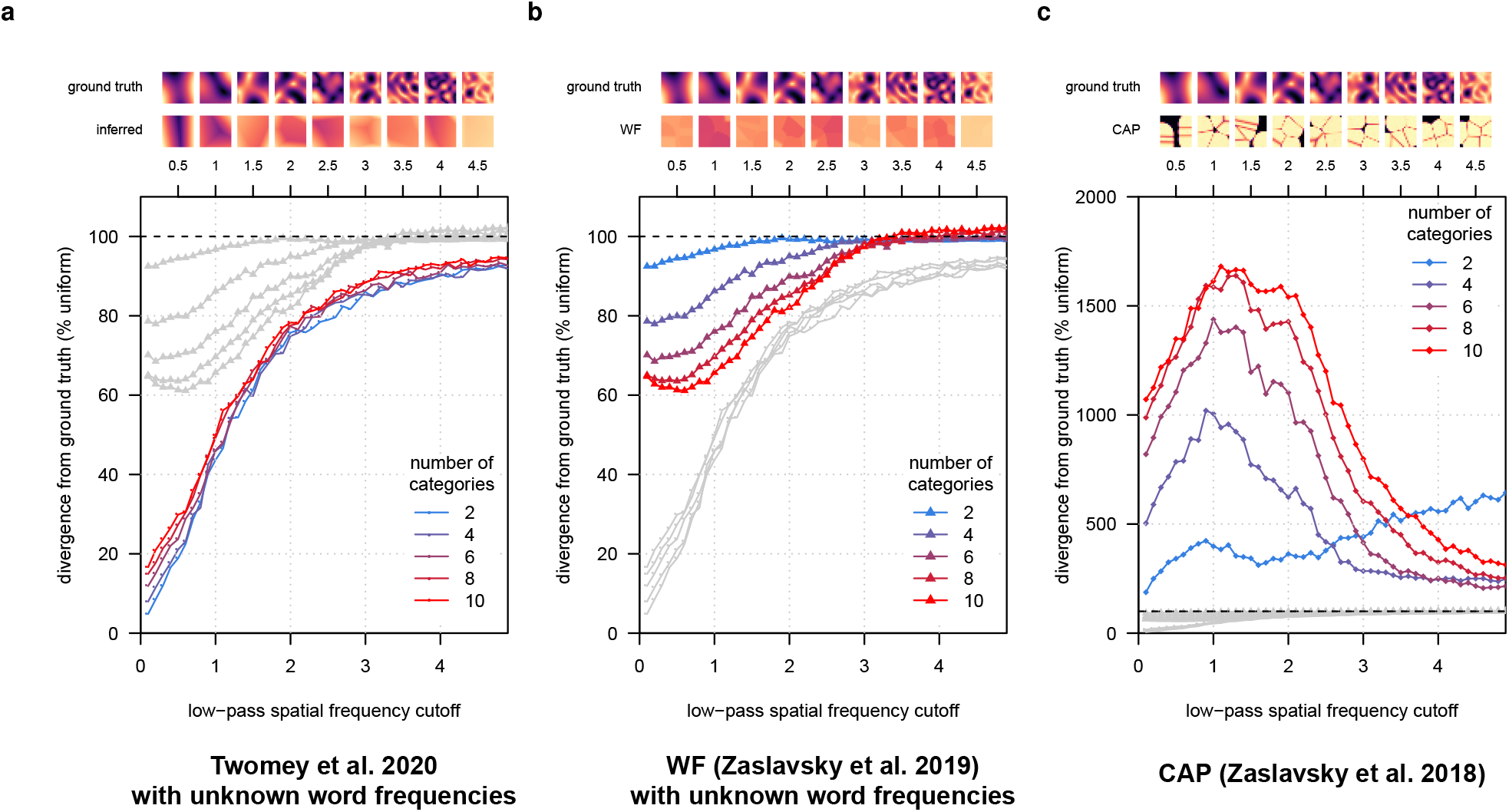
Systematic comparison of inference methods for language-specific communicative needs when word frequencies are unknown (e.g. in the WCS). Ground truth distributions of communicative need were generated with spatial variation up to a given cutoff spatial frequency (x-axis), and scaled such that the entropy was held constant across all generated examples. Accuracy of inference (y-axis; lower is better) is measured as the KL-divergence between inferred versus ground truth communicative needs, for each example, expressed as a percentage of the KL-divergence between uniform and ground truth (dashed line at 100%). Curves are shown for inferences based on rate-distortion vocabularies for 2, 4, 6, 8, and 10 “words” (number of categories), when word-frequencies are unknown. **(a)** Inferences using the method proposed in this paper (Twomey et al); **(b)** based on the word-frequency (WF) method of Zaslavsky et al. [17]; and **(c)** based on the CAP approach of Zaslavsky et al. [7]. For language-specific inferences, our method achieves high accuracy for recovering low-frequency information about the true communicative need, and it does so in a manner that is invariant to the number of available “words” (categories).

By contrast, the WF method (SI Fig. D10b) is highly sensitive to the number of categories available to it through 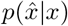, and it requires a large number of categories to improve by even 40% over uniform when 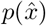 is unknown (assumed uniform). High spatial frequency information is lost, and when word frequencies are unknown this apparently inhibits recovery of low-frequency information as well. Performance at intermediate scales can approach our method for large enough vocabularies (number of categories), but even at these scales our method provides consistently better performance across vocabulary sizes. Inferences by CAP (SI Fig. D10c) never improve over uniform and they are highly sensitive to vocabulary size (number of categories).

## E Language-specific inferences of communicative needs

In the following figures (SI Fig. E11–E27), we show each of the 130 language-specific distributions of communicative need we inferred using our method, for all languages recorded in the WCS and B&K survey data.

**Figure E11.**
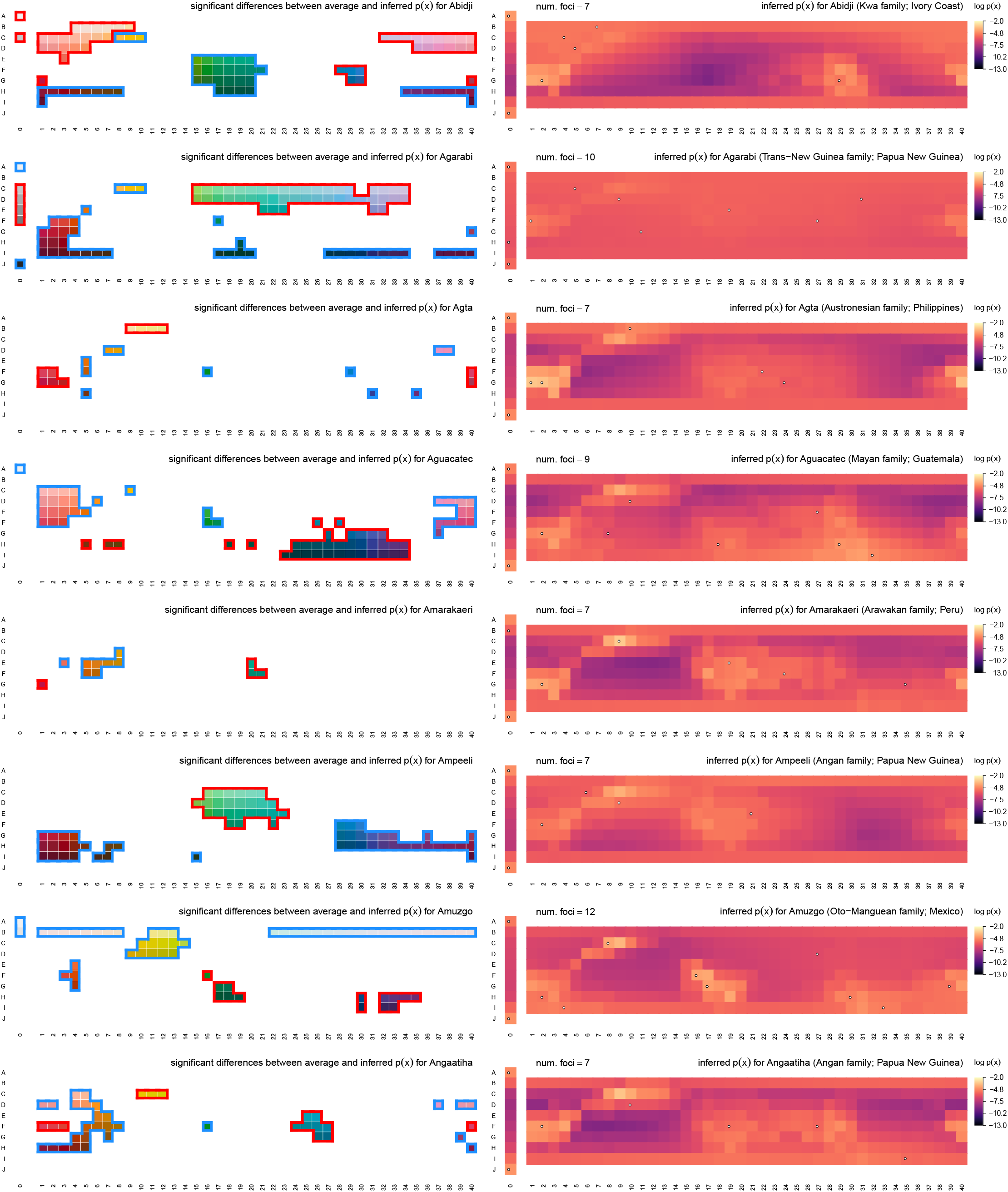
Inferred communicative needs for 130 languages on a common scale. Each row corresponds to a language in the combined WCS+B&K survey data. (*Left column*) Significant differences between language-specific and across-language average communicative needs, shown as in Fig. 5. Deviations that exceed *σ*/2 with 95% confidence are highlighted in red (elevated) or blue (suppressed). (*Right column*) Language-specific communicative needs (log scale) shown with language focal color positions projected on to the WCS color chips (white points). Focal points may overlap on the same color chip.

**Figure E12.**
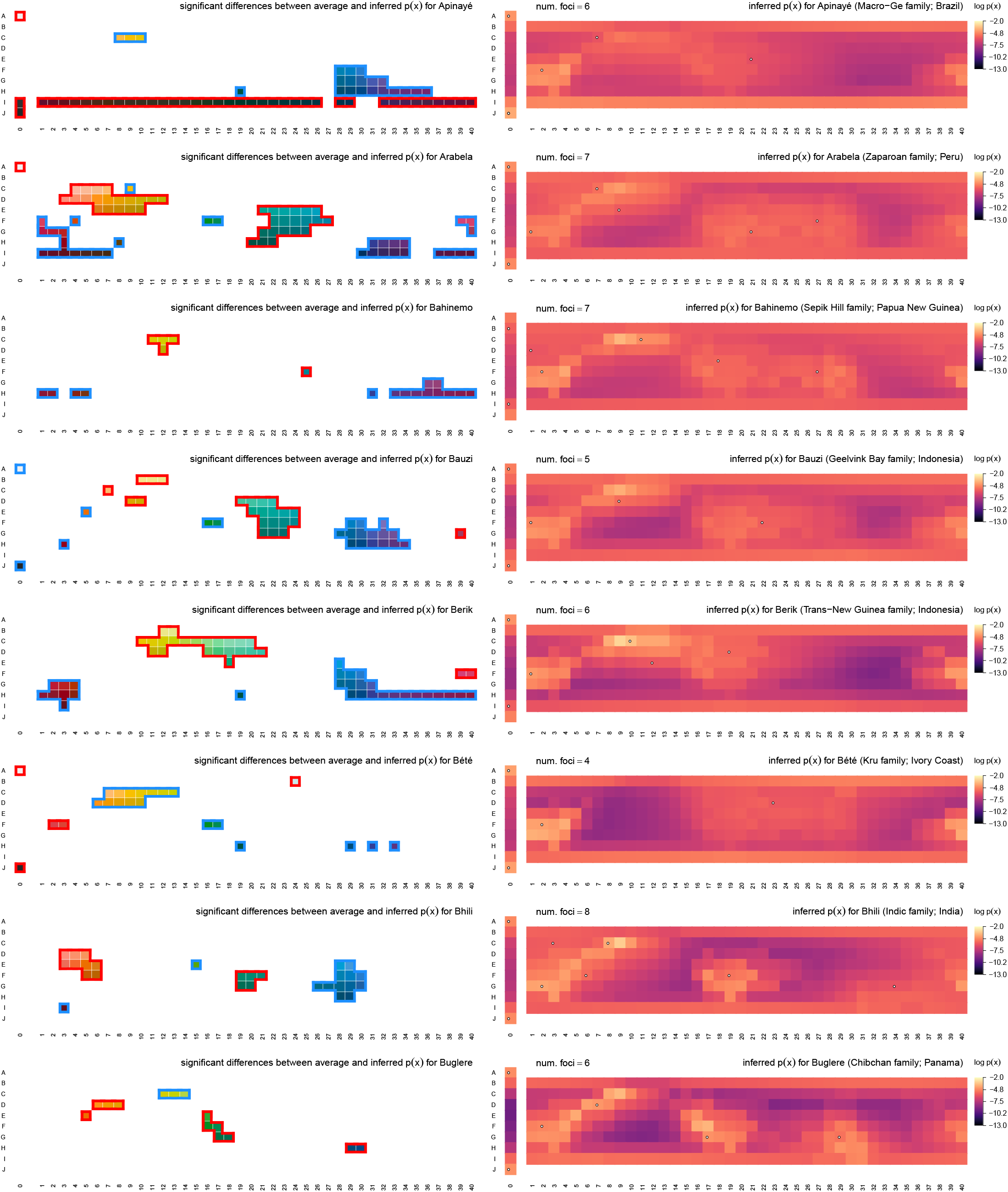
Inferred communicative needs for 130 languages on a common scale (continued).

**Figure E13.**
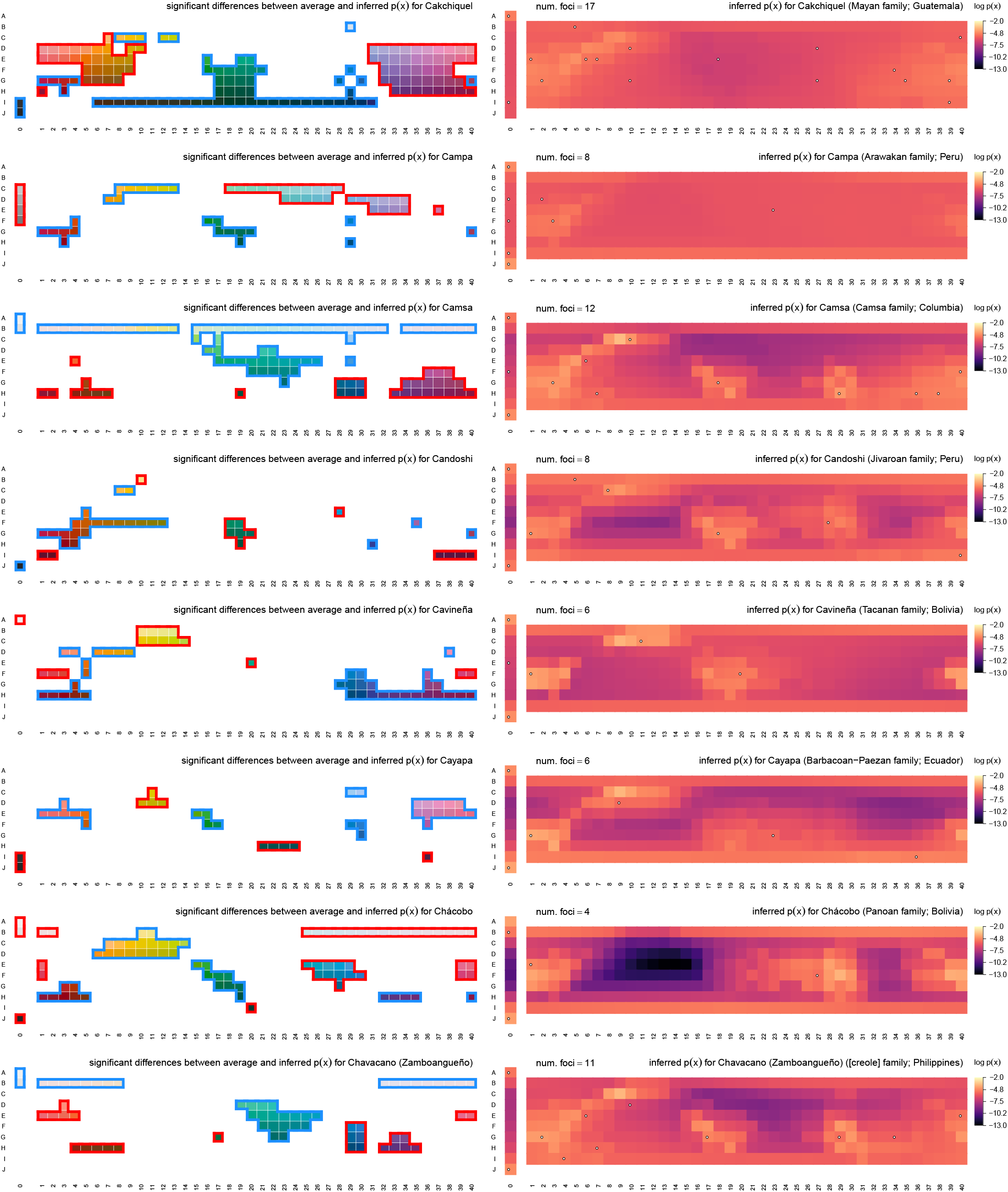
Inferred communicative needs for 130 languages on a common scale (continued).

**Figure E14.**
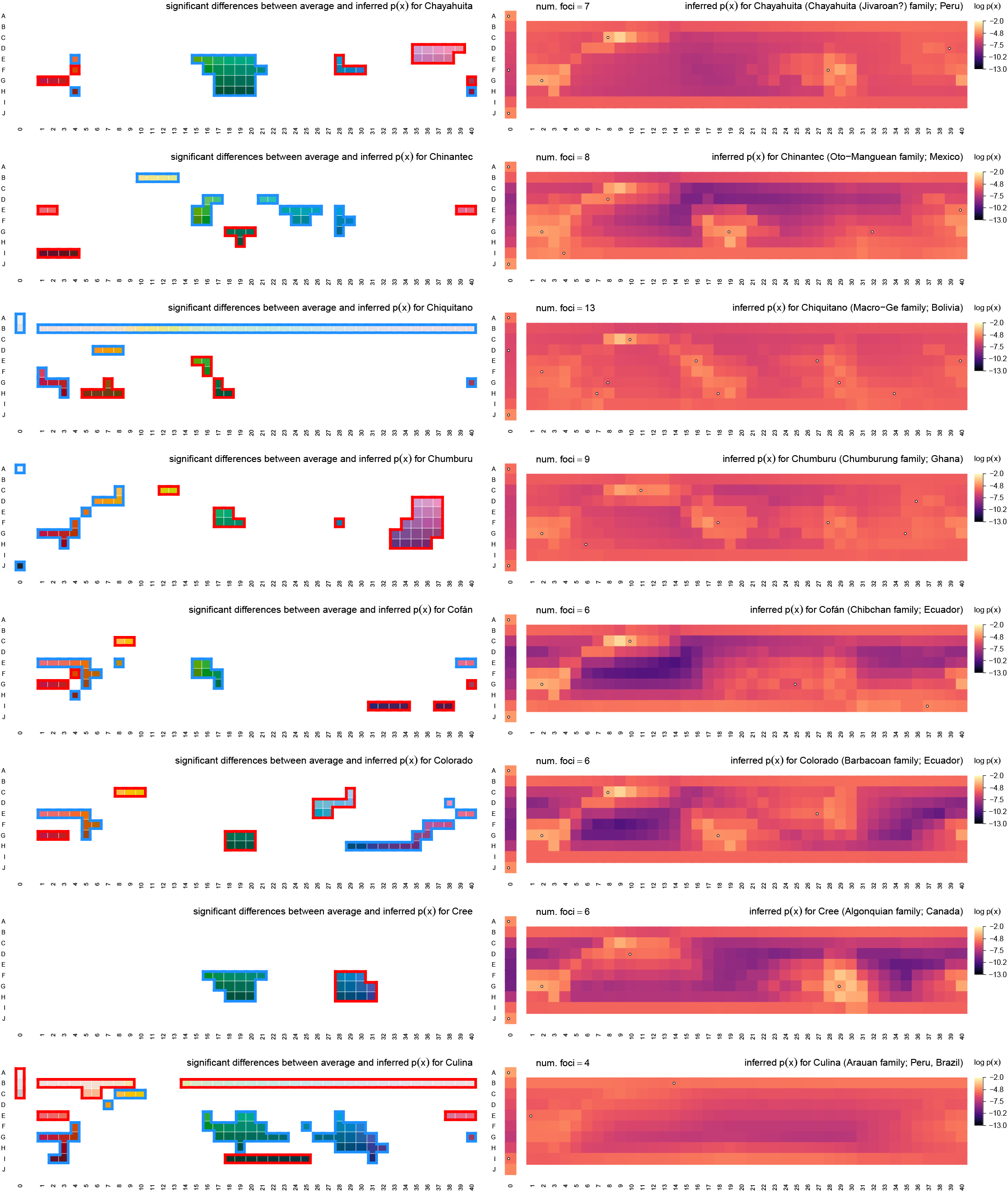
Inferred communicative needs for 130 languages on a common scale (continued).

**Figure E15.**
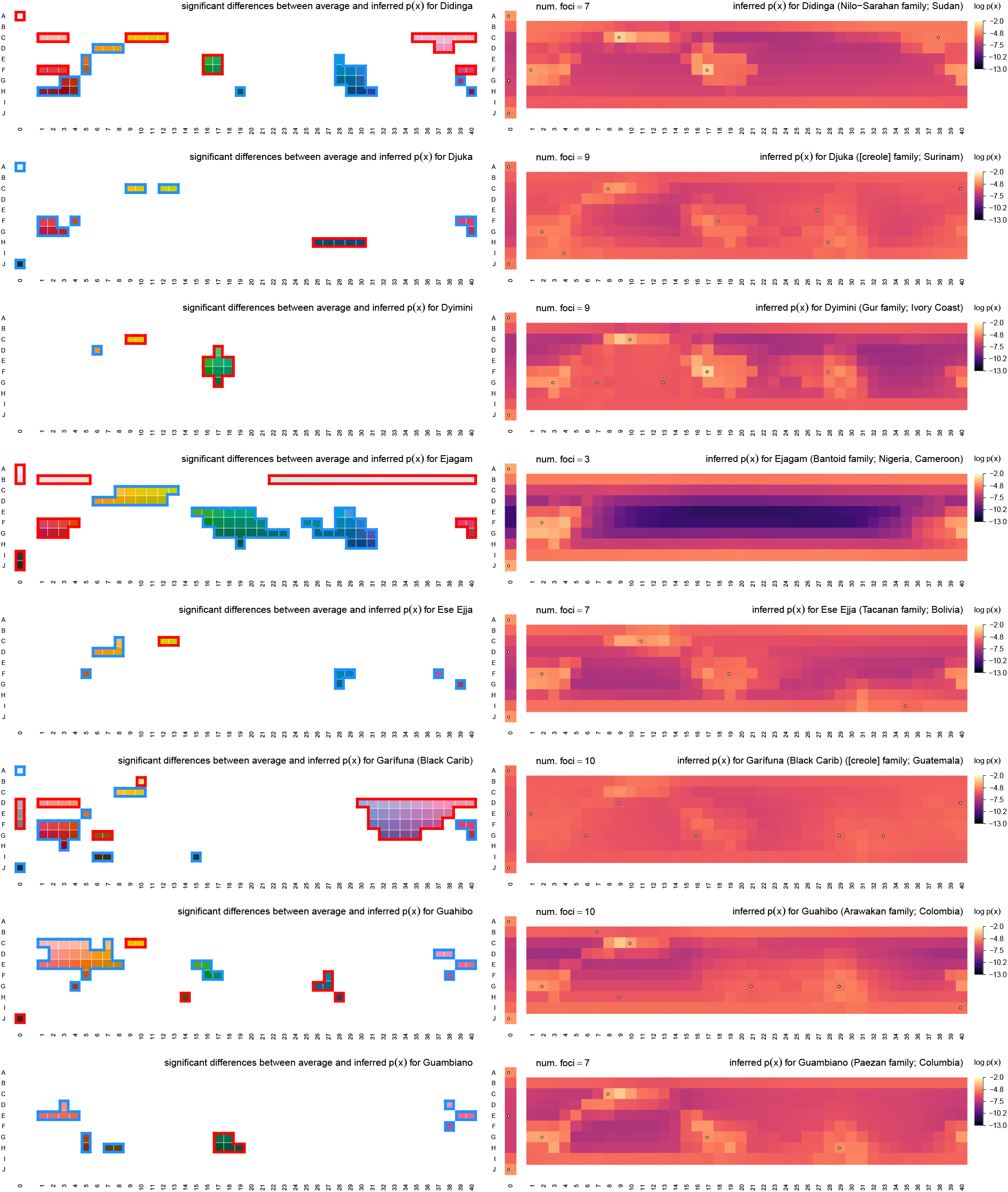
Inferred communicative needs for 130 languages on a common scale (continued).

**Figure E16.**
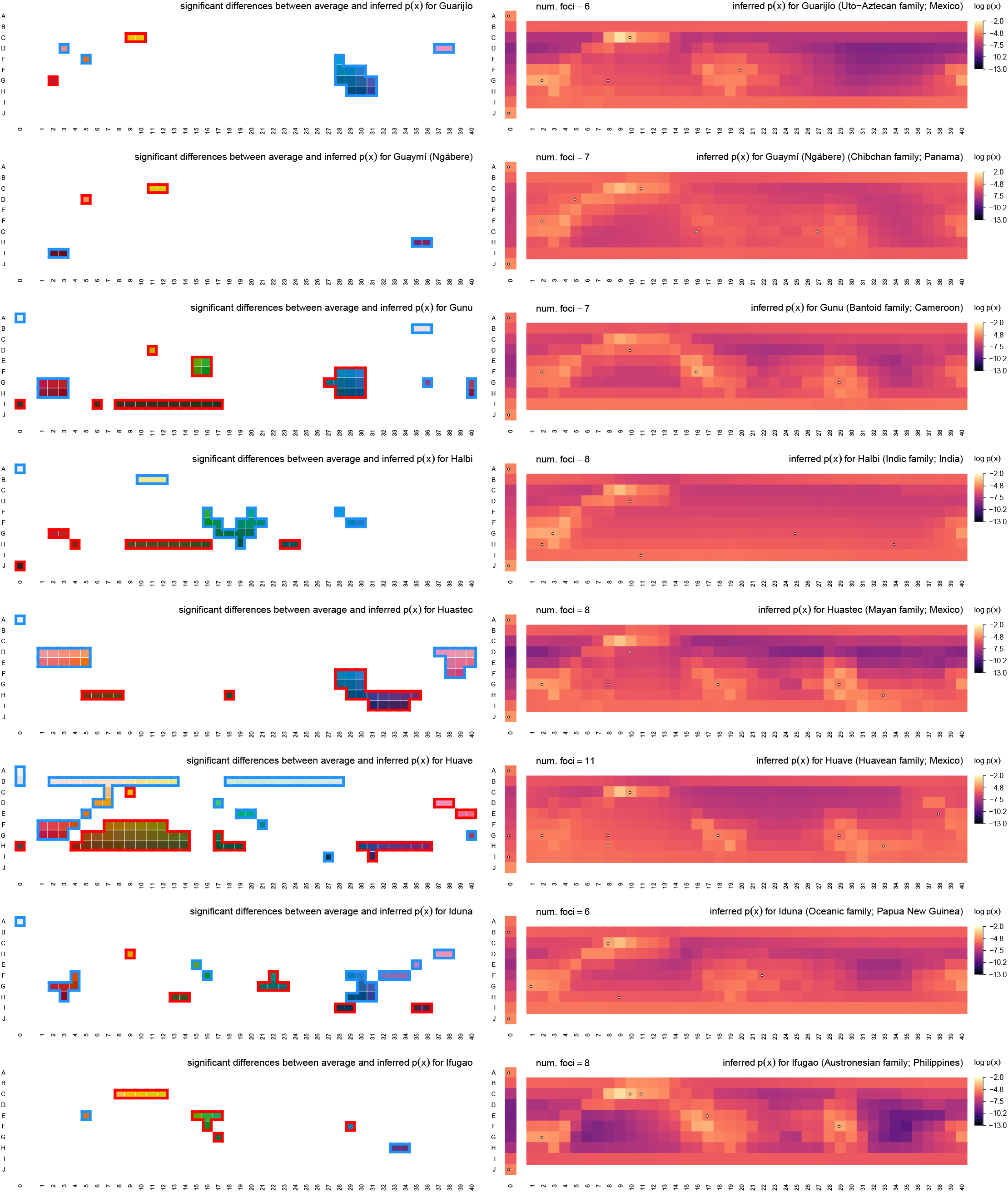
Inferred communicative needs for 130 languages on a common scale (continued).

**Figure E17.**
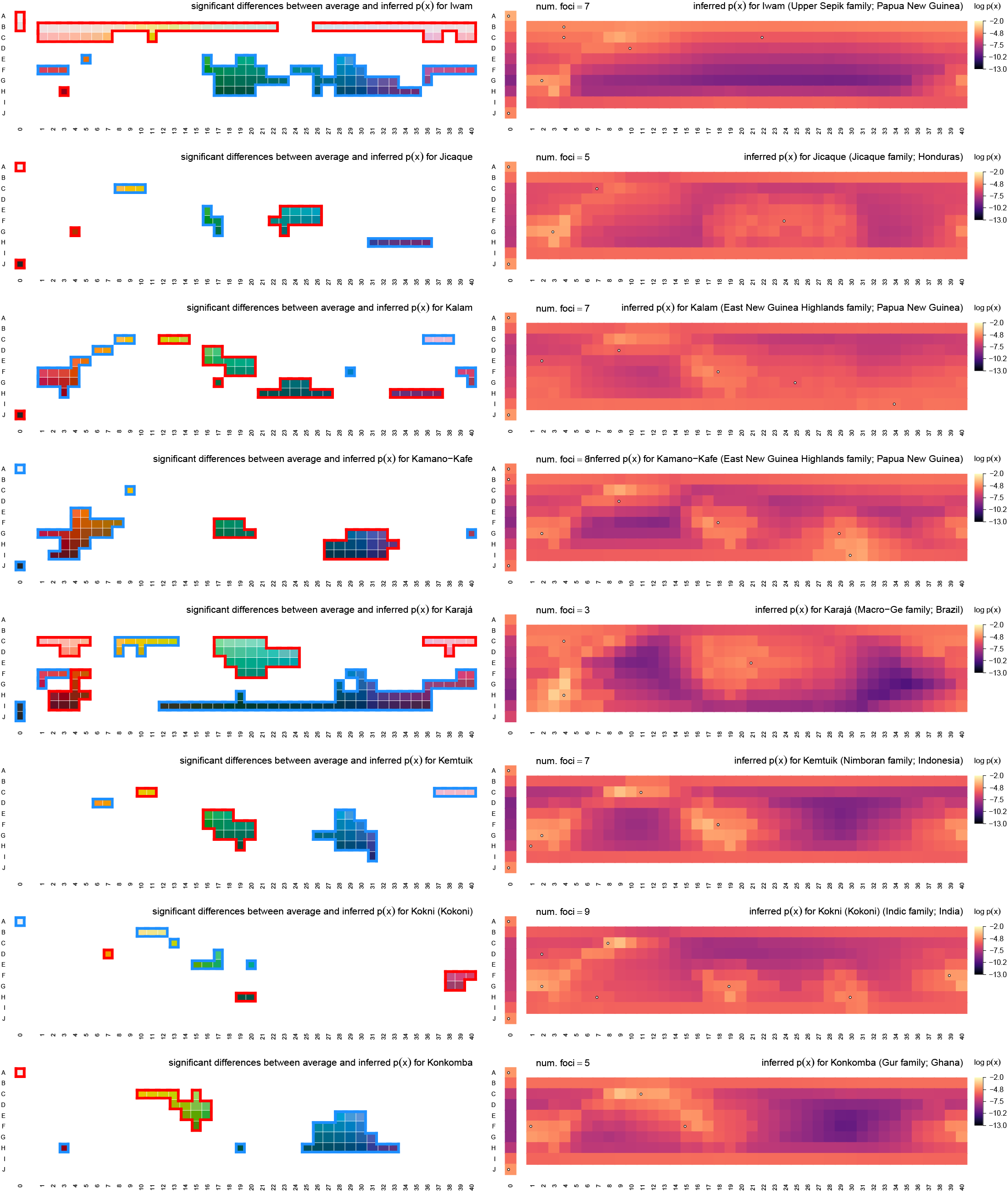
Inferred communicative needs for 130 languages on a common scale (continued).

**Figure E18.**
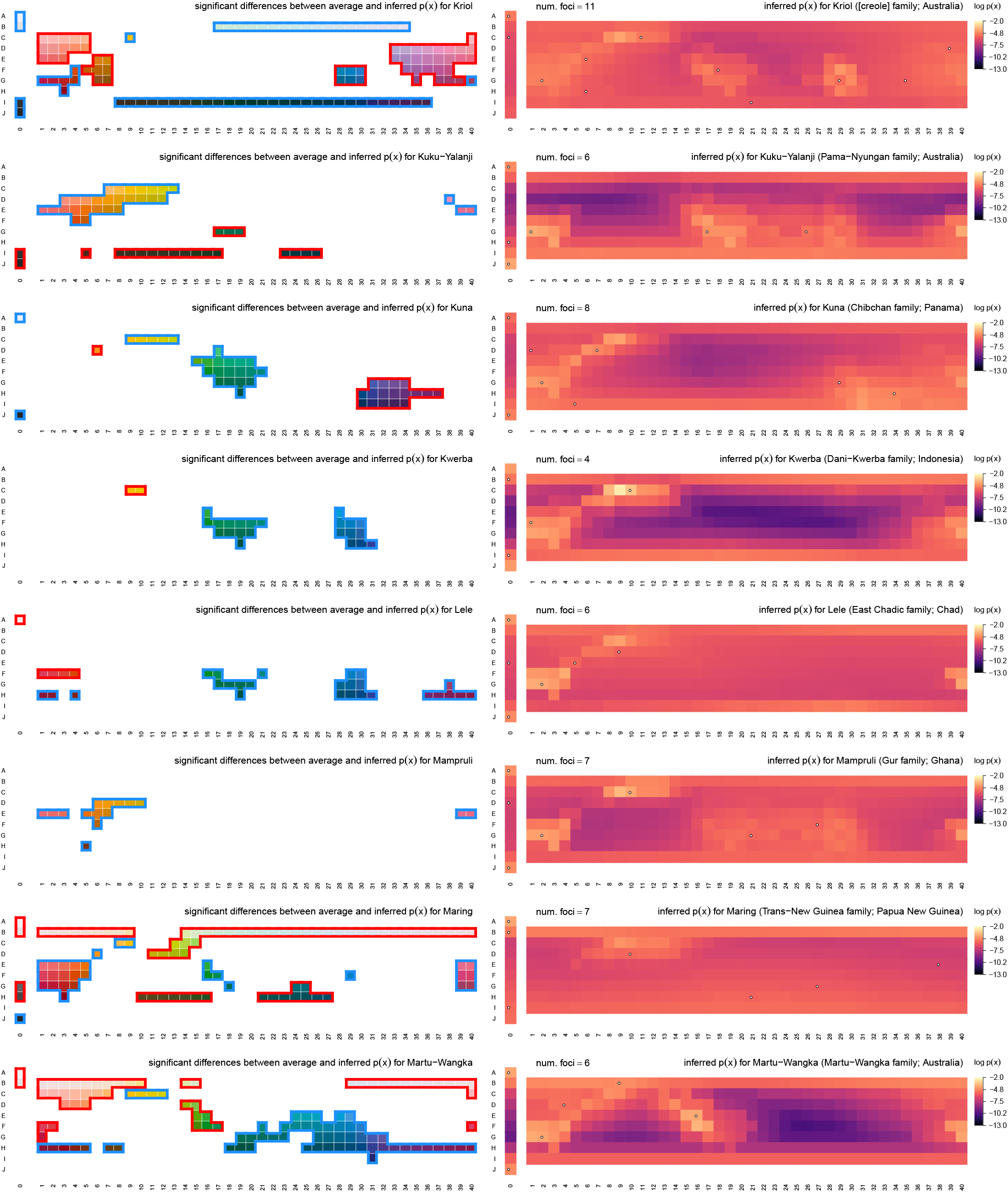
Inferred communicative needs for 130 languages on a common scale (continued).

**Figure E19.**
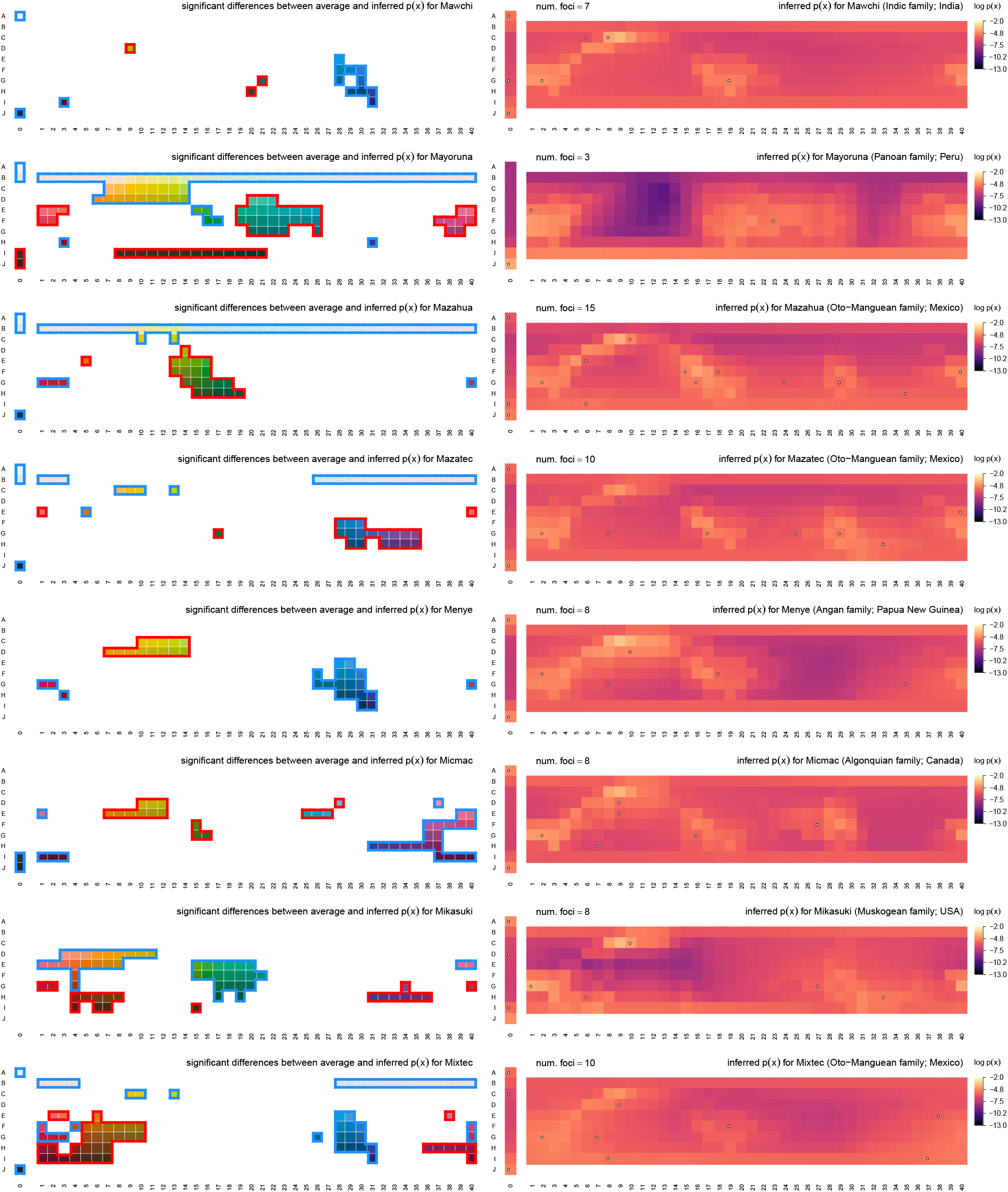
Inferred communicative needs for 130 languages on a common scale (continued).

**Figure E20.**
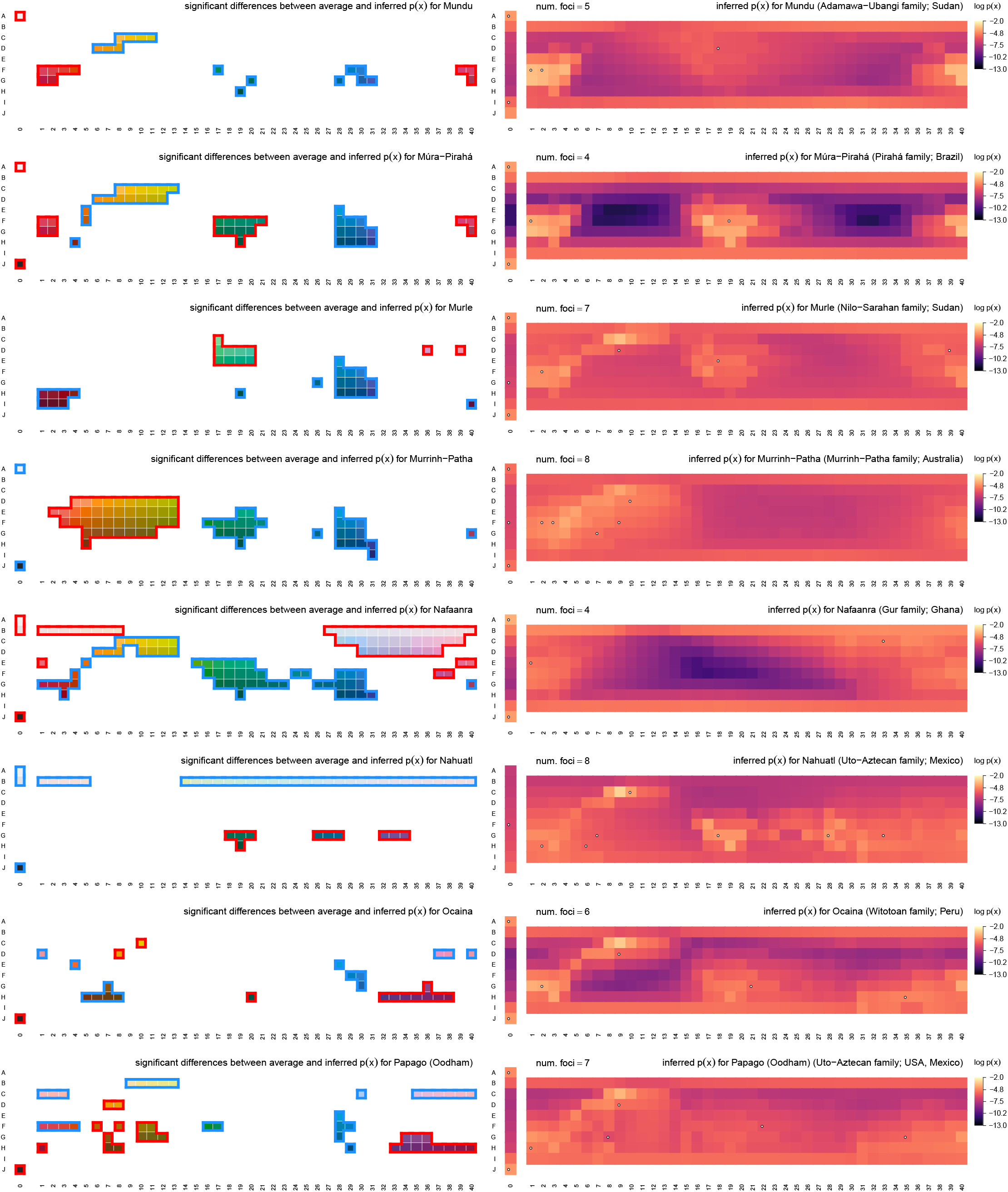
Inferred communicative needs for 130 languages on a common scale (continued).

**Figure E21.**
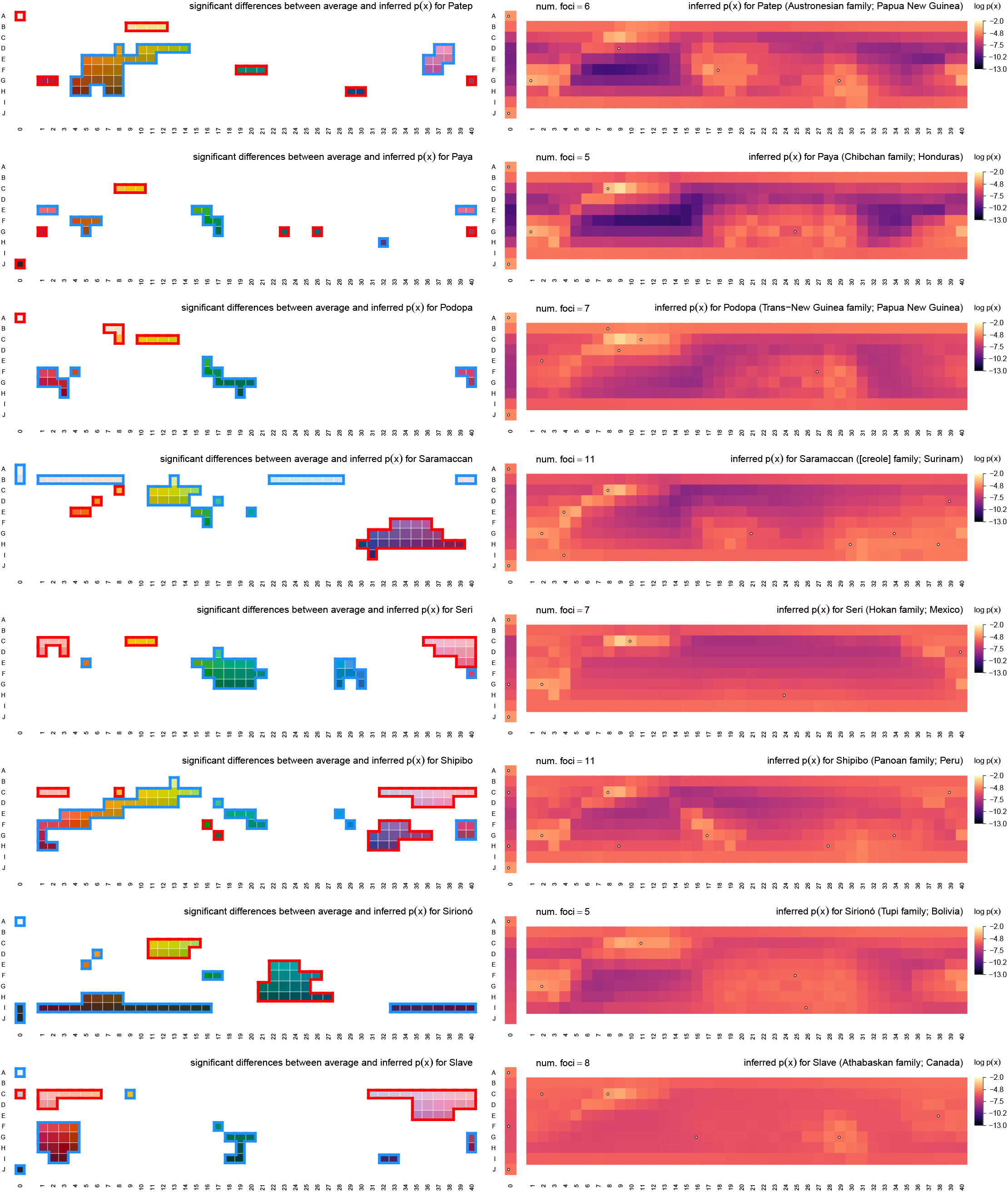
Inferred communicative needs for 130 languages on a common scale (continued).

**Figure E22.**
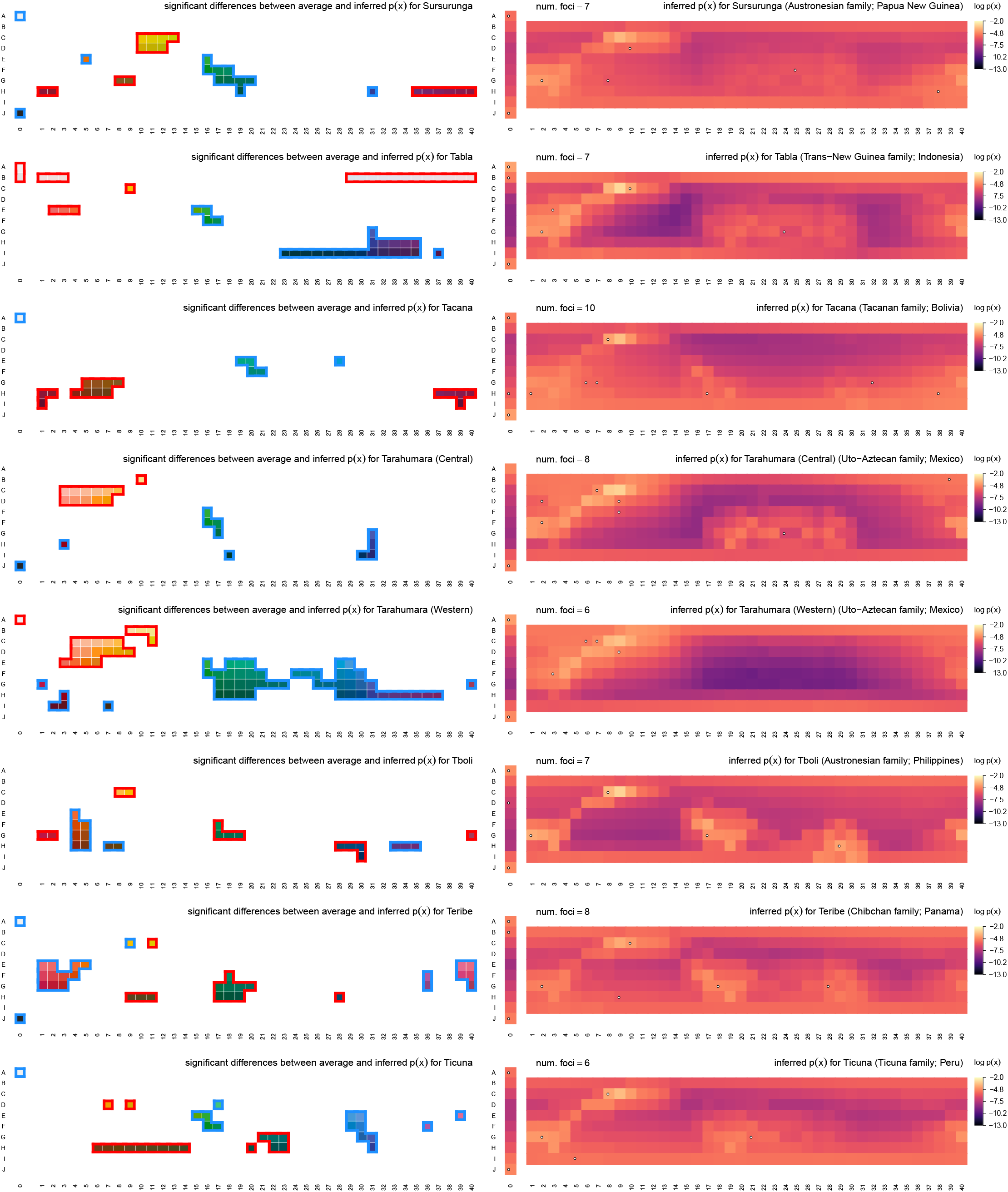
Inferred communicative needs for 130 languages on a common scale (continued).

**Figure E23.**
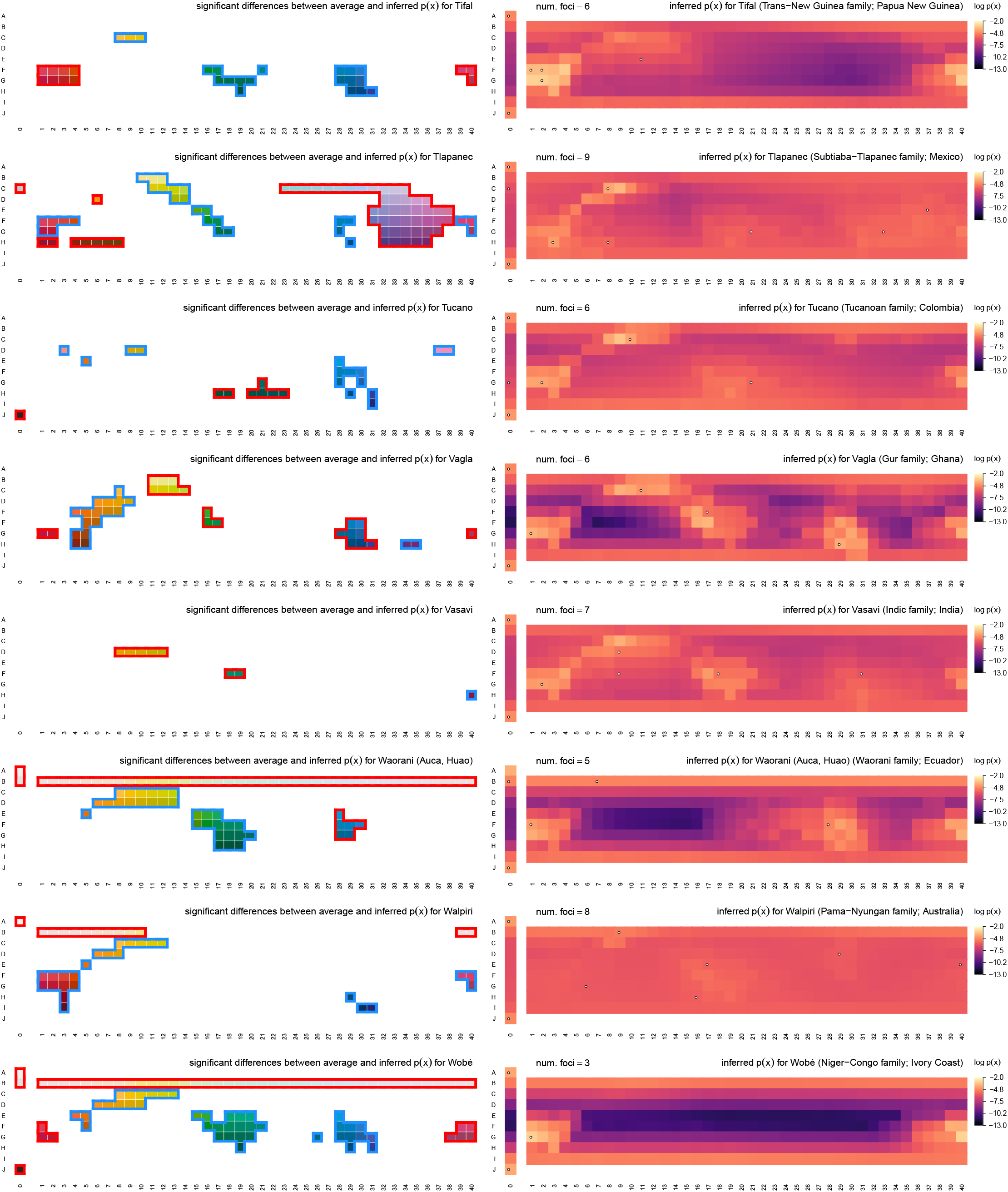
Inferred communicative needs for 130 languages on a common scale (continued).

**Figure E24.**
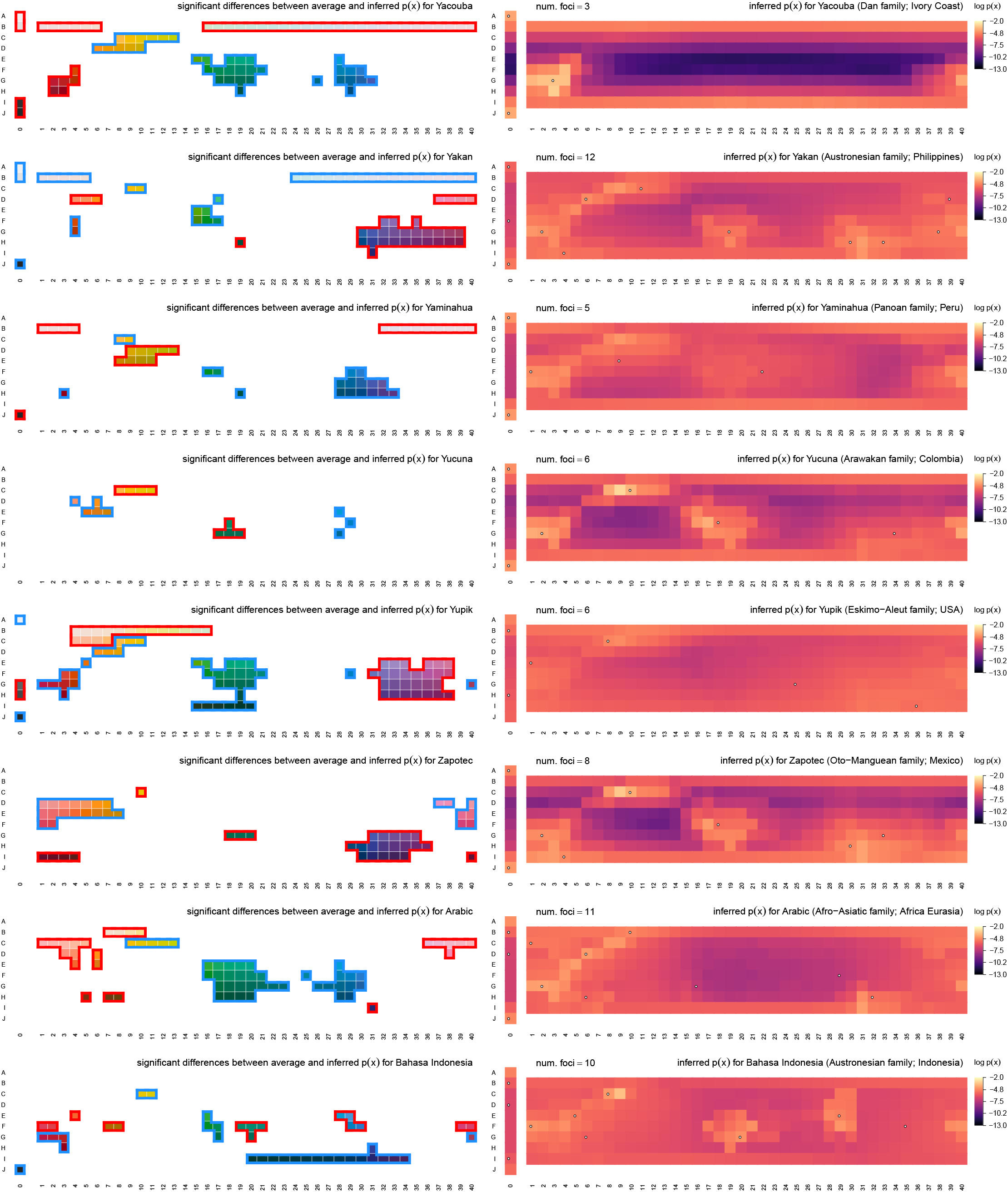
Inferred communicative needs for 130 languages on a common scale (continued).

**Figure E25.**
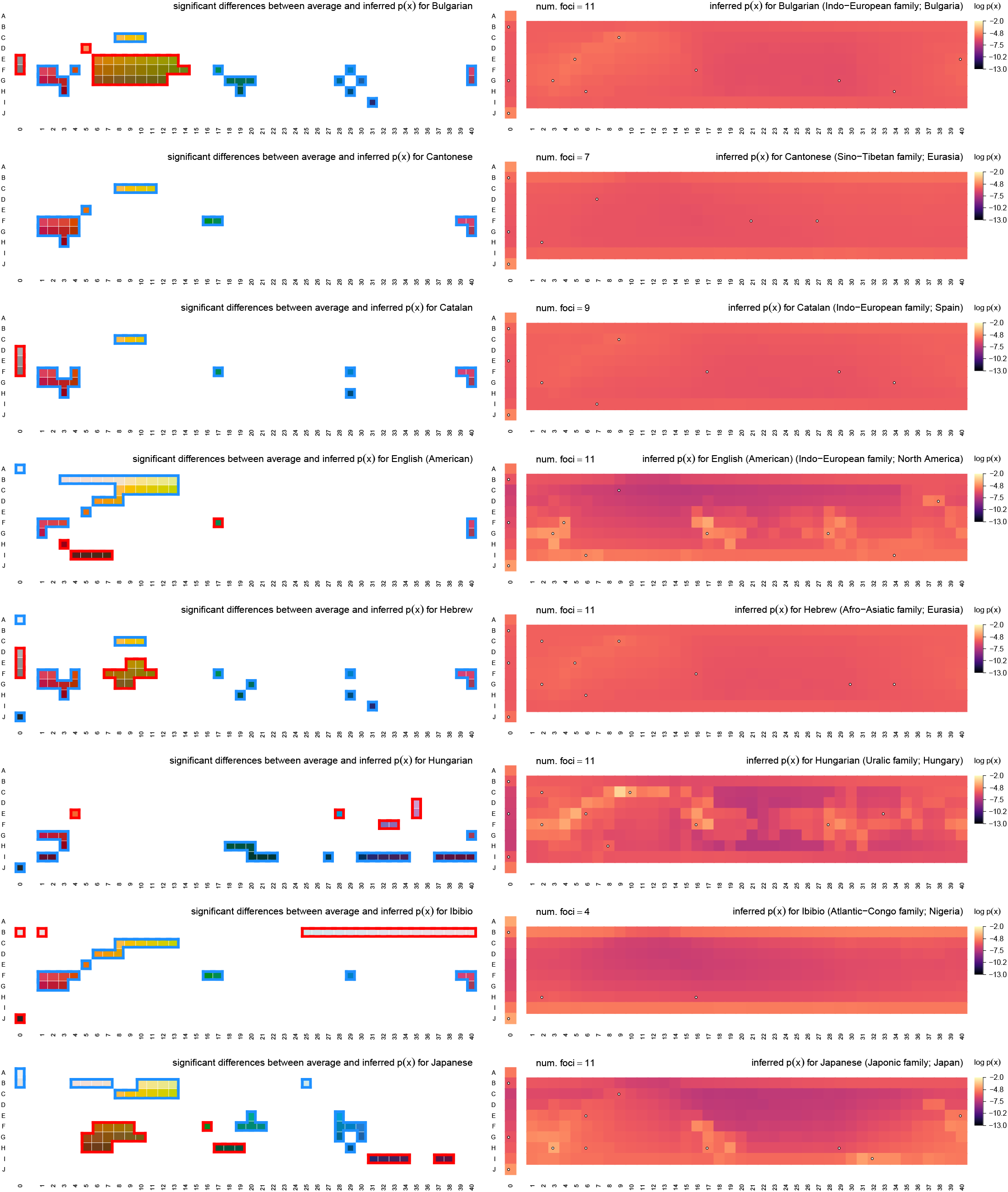
Inferred communicative needs for 130 languages on a common scale (continued).

**Figure E26.**
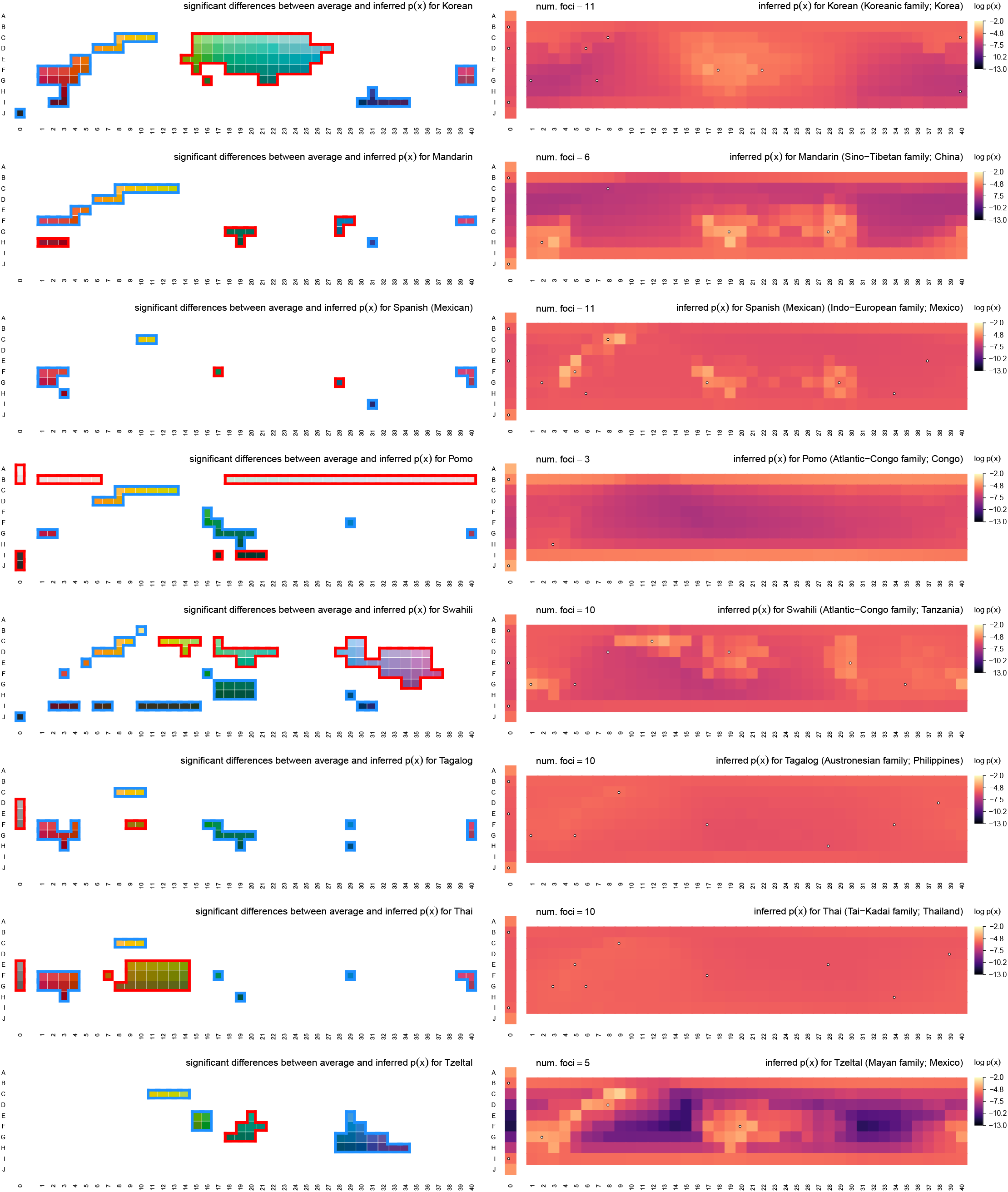
Inferred communicative needs for 130 languages on a common scale (continued).

**Figure E27.**
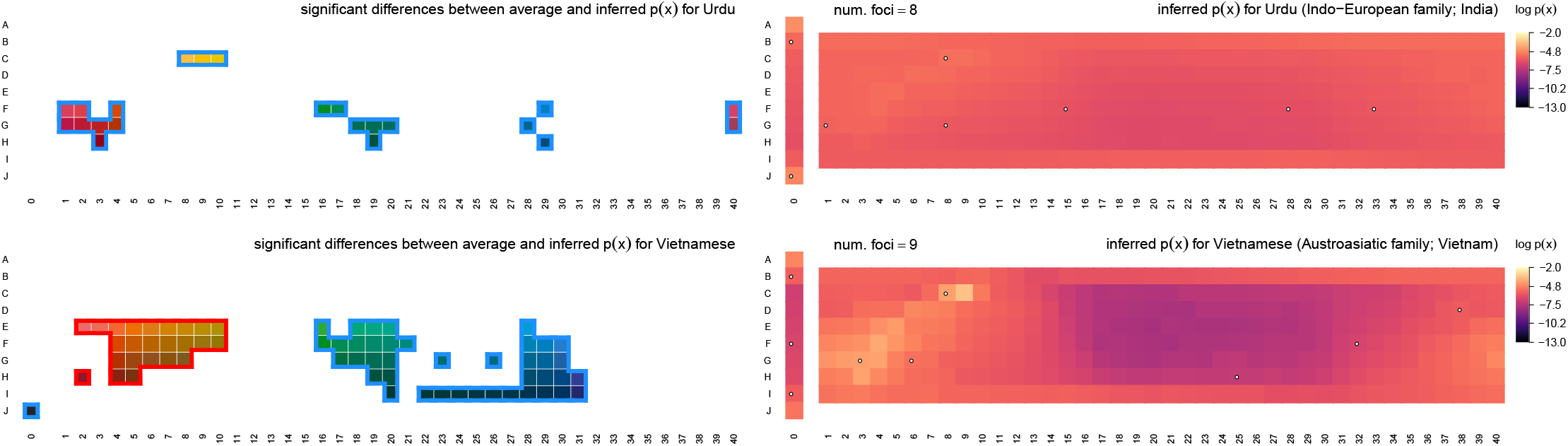
Inferred communicative needs for 130 languages on a common scale (continued).

## Footnotes

† Or by a mixture of the best choice focal colors when there was more than one best choice.

† More precisely, we propose the measured focal colors are the best approximation to the true centroid among the set of WCS color stimuli.

† Nor do we use empirical term maps for selection among the small set of nonunique rate-distortion optimal solutions. In this study, selection is based on focal points alone. See SI Sec. C.

† Note that these results do not imply shared communicative needs are determined by the need to name fruit specifically.

† Although this does not imply that color terms in Pirahã are abstract necessarily; see Regier et al. [60].

† The Munsell color system was created as a means to index human perceivable color by hue, value, and chroma, at empirically measured perceptually uniform intervals along each dimension. In the WCS notation, rows correspond to equally spaced Munsell values, and columns 1–40 correspond to equally spaced Munsell hues. For column 0 Munsell chroma is 0; for all other columns Munsell chroma was chosen as the maximum for the given hue and value.

‡ http://vision.psychol.cam.ac.uk/spectra

† *http://www1.icsi.berkeley.edu/wcs/data.html*

† See also Byrne [16] as a helpful reference.

† The focal color assays of B&K and the WCS were essentially the same, however. Hence the inclusion of both data sets in other analyses where only focal color estimates are necessary.

## References

1. Berlin, B. & Kay, P. (1969) Basic Color Terms: Their Universality and Evolution. Univ. of California Press, Berkeley.

2. Regier, T., Kay, P., & Khetarpal, N. (2007) Color naming reflects optimal partitions of color space. PNAS, 104(4):1436–1441.

3. Gibson, E., Futrell, R., Jara-Ettinger, J., Mahowald, K., Bergen, L., Ratnasingam, S., Gibson, M., Piantadosi, S. T., & Conway, B. R. (2017) Color naming across languages reflects color use. PNAS, 114(40):10785–10790.

4. Kay, P., Berlin, B., Maaffi, L., Merrifield, W., & Cook, R. (2009) The World Color Survey. CLSI, Standford. ISBN 9781575864150.

5. Kay, P. & Regier, T. (2003) Resolving the question of color naming universals. PNAS, 100(15):9085–9089.

6. Kay, P. (2005) Color categories are not arbitrary. Cross-Cult. Res., 39(1):39–55.

7. Regier, T., Kay, P., & Cook, R. (2005) Focal colors are universal after all. PNAS, 102(23):8386–8391.

8. Lindsey, D. T. & Brown, A. M. (2009) World color survey color naming reveals universal motifs and their within-language diversity. PNAS, 106(47):19785–19790.

9. Yendrikhovskij, S. N. (2001) Computing color categories from statistics of natural images. J. Imaging Sci. Technol., 45(5):409–417.

10. Komarova, N. L., Jameson, K. A., & Narens, L. (2007) Evolutionary models of color categorization based on discrimination. J. Math. Psychol., 51(6):359–382.

11. Zaslavsky, N., Kemp, C., Regier, T., & Tishby, N. (2018) Efficient compression in color naming and its evolution. PNAS, 115(31):7937–7942.

12. Webster, M. A., Miyahara, E., Malkoc, G., & Raker, V. E. (2000) Variations in normal color vision. ii. unique hues. J. Opt. Soc. Am. A, 17:1545–1555.

13. Schefrin, B. E. & Werner, J. S. (1990) Loci of spectral unique hues throughout the lifespan. J. Opt. Soc. Am. A, 7(2):305–311.

14. Brainard, D. H., Roorda, A., Yamauchi, Y., Calderone, J. B., Metha, A., Neitz, M., Neitz, J., Williams, D. R., & Jacobs, G. H. (2000) Functional consequences of the relative number of l and m cones. J. Opt. Soc. Am. A, 17:607–614.

15. Neitz, J., Carroll, J., Yamauchi, Y., Neitz, M., & Williams, D. (2002) Color perception is mediated by a plastic neural mechanism that is adjustable in adults. Neuron, 35(4):783–792.

16. Kay, P. & McDaniel, C. K. (1978) The linguistic significance of the meanings of basic color terms. Language, 54(3):610–646.

17. Heider-Rosch, E. (1972) Universals in color naming and memory. J. Exp. Psychol. Gen., 93(1):10–20.

18. Rosch, E. (1973) Natural categories. Cog. Psychol., 4(3):328–350.

19. Sun, R. K. (1983) Perceptual distances and the basic color term encoding sequence. Am. Anthropol., 85(2):387–391.

20. MacLaury, R. E. (1987) Color-category evolution and shuswap yellow-with-green. Am. Anthropol., 89(1):107–124.

21. MacLaury, R. E. (1992) From brightness to hue: an explanatory model of color-category evolution. Curr. Anthropol., 33(2):137–186.

22. Jameson, K. & D’Andrade, R. G. (1997) It’s not really red, green, yellow, blue: an inquiry into perceptual color space. In Hardin, C. L. & Maffi, L., editors, Color Categories in Thought and Language. Cambridge University Press, Cambridge, UK.

23. Webster, M. A. & Kay, P. Individual and population differences in focal colors. In MacLaury, R. E., Paramei, G. V., & Dedrick, D., editors, Anthropology of Color: Interdisciplinary multilevel modeling, pages 29–53, Amsterdam / Philadelphia, (2007). John Benjamins Publishing Company.

24. Gibson, E., Futrell, R., Piantadosi, S. P., Dautriche, I., Mahowald, K., Bergen, L., & Levy, R. (2019) How efficiency shapes human language. Trends Cogn. Sci., 23(5):389–407.

25. Conway, B. R., Ratnasingam, S., Jara-Ettinger, J., Futrell, R., & Gibson, E. (2020) Communication efficiency of color naming across languages provides a new framework for the evolution of color terms. Cognition, 195:104086.

26. Shepard, R. N. (1992) The perceptual organization of colors: an adaptation to regularities of the terrestrial world? In Barkow, J., Cosmides, L., & Tooby, J., editors, Adapted Minds, pages 495–532. Oxford University Press, Oxford, UK.

27. Zaslavsky, N., Kemp, C., Tishby, N., & Regier, T. (2018) Color naming reflects both perceptual structure and communicative need. Cog. Sci., 11(1):207–219.

28. Zaslavsky, N., Kemp, C., Tishby, N., & Regier, T. (2019) Communicative need in colour naming. Cogn. Neuropsychol., 37(5-6):312–324.

29. Shannon, C. E. (1948) A mathematical theory of communication. Bell System Technical Journal, 27(3):379–423.

30. Shannon, C. E. (1959) Coding theorems for a discrete source with a fidelity criterion. IRE National Convention Record, 7(4):142–163.

31. Cover, T. M. & Thomas, J. A. (2006) Elements of information theory. Wiley-Interscience, 2nd edition.

32. Sims, C. R. (2016) Rate-distortion theory and human perception. Cognition, 152:181–198.

33. Kemp, C., Xu, Y., & Regier, T. (2018) Semantic typology and efficient communication. Annu. Rev. Linguist., 4(1):109–128.

34. Boynton, R. M. & Olson, C. X. (1987) Locating basic colors in the osa space. Color Res. Appl., 12(2):94–105.

35. Sturges, J. & Whitfield, T. W. A. (1995) Locating basic colours in the munsell space. Color Res. Appl., 20(6):364–376.

36. Lindsey, D. T. & Brown, A. M. (2014) The color lexicon of american english. J. Vision, 14(2):17, 1–25.

37. Liu, T., Sun, J., Zhen, N.-N., Tang, X., & Shum, H.-Y. Learning to detect a salient object. In IEEE Conference on Computer Vision and Pattern Recognition, Minneapolis, MN, USA, (2007). doi: 10.1109/CVPR.2007.383047.

38. Sumner, P. & Mollon, J. D. (2000) Catarrhine photopigments are optimized for detecting targets against a foliage background. J. Exp. Biol., 203(13):1963–1986.

39. Sumner, P. & Mollon, J. D. (2000) Chromaticity as a signal of ripeness in fruits taken by primates. J. Exp. Biol., 203(13):1987–2000.

40. Lomáscolo, S. B., Levey, D. J., Kimball, R. T., Bolker, B. M., & Alborn, H. T. (2010) Dispersers shape fruit diversity in *Ficus* (moraceae). PNAS, 107(33):14668–14672.

41. Nevo, O., Valenta, K., Razafimandimby, D., Melin, A. D., Ayasse, M., & Chapman, C. A. (2018) Frugivores and the evolution of fruit colour. Biol. Lett., 14(9). doi: 10.1098/rsbl.2018.0377.

42. Valenta, K. & Nevo, O. (2020) The dispersal syndrome hypothesis: how animals shaped fruit traits, and how they did not. Funct. Ecol. doi: 10.1111/1365-2435.13564.

43. Regan, B. C., Julliot, C., Simmen, B., Viénot, F., Charles-Dominique, P., & Mollon, J. D. (1998) Frugivory and colour vision in *Alouatta seniculus*, a trichromatic platyrrhine monkey. Vision Res., 38(21):3321–3327.

44. Regan, B. C., Julliot, C., Simmen, B., Viénot, F., Charles-Dominique, P., & Mollon, J. D. (2001) Fruits, foliage and the evolution of primate colour vision. Phil. Trans. R. Soc. Lond. B, 356(1407):229–283.

45. Onstein, R. E., Vink, D. N., Veen, J., Barratt, C. D., Flantua, S. G. A., Wich, S. A., & Kissling, W. D. (2020) Palm fruit colours are linked to the broad-scale distribution and diversification of primate colour vision systems. Proc. R. Soc. B, 287(1921). doi: 10.1098/rspb.2019.2731.

46. Hammarström, H., Forkel, R., Haspelmath, M., & Bank, S. Glottolog 4.2.1. Max Planck Institute for the Science of Human History, (2020). URL https://glottolog.org.

47. Olson, D. M., Dinerstein, E., Wikramanayake, E. D., Burgess, N. D., Powell, G. V. N., Underwood, E. C., D’Amico, J. A., Itoua, I., Strand, H. E., Morrison, J. C., Loucks, C. J., Allnutt, T. F., Ricketts, T. H., Kura, Y., Lamoreaux, J. F., Wettengel, W. W., Hedao, P., & Kassem, K. R. (2001) Terrestrial ecoregions of the world: a new map of life on earth. Bioscience, 51(11):933–938.

48. Smith, J. R., Letten, A. D., Ke, P.-J., B., A. C., Hendershot, J. N., Dhami, M. K., Dlott, G. A., Grainger, T. N., Howard, M. E., Morrison, B. M., Routh, D., San Juan, P. A., Mooney, H. A., Mordecai, E. A., Crowther, T. W., & Daily, G. C. (2018) A global test of ecoregions. Nat. Ecol. Evol., 2:1889–1896.

49. Clarke, R. T., Rothery, P., & Raybould, A. F. (2002) Confidence limits for regression relationships between distance matrices: estimating gene flow with distance. J. Agric. Biol. Envir. S., 7(3):361–372.

50. Majid, A., Roberts, S. G., Cilissen, L., Emmorey, K., Nicodemus, B., O’Grady, L., Woll, B., LeLan, B., de Sousa, H., Cansler, B. L., Shayan, S., de Vos, C., Senft, G., Enfield, N. J., Razak, R. A., Fedden, S., Tufvesson, S., Dingemanse, M., Ozturk, O., Brown, P., Hill, C., Le Guen, O., Hirtzel, V., van Gijn R., Sicoli, M. A., & Levinson, S. C. (2018) Differential coding of perception in the world’s languages. PNAS, 115(45):11369–11376.

51. Caves, E. M., Green, P. A., Zipple, M. N., Peters, S., Johnsen, S., & Nowicki, S. (2018) Categorical perception of colour signals in a songbird. Nature, 560(7718):365–367.

52. Zipple, M. N., Caves, E. M., Green, P. A., Peters, S., Johnsen, S., & Nowicki, S. (2019) Categorical colour perception occurs in both signalling and non-signalling colour ranges in a songbird. Proc. R. Soc. B, 286(1903). doi: 10.1098/rspb.2019.0524.

53. Davidoff, J., Davies, I., & Roberson, D. (1999) Colour categories in a stone-age tribe. Nature, 398(6724):203–204.

54. Roberson, D. & Davidoff, J. (2000) Color categories are not universal: replications and new evidence from a stone–age culture. J. Exp. Psychol., 129(3):369–398.

55. Roberson, D., Davidoff, J., Davies, I. R. L., & Shapiro, L. R. (2005) Color categories: evidence for the cultural relativity hypothesis. Cogn. Psychol., 50(4):378–411.

56. Majid, A. & Kruspe, N. (2018) Hunter-gatherer olfaction is special. Curr. Biol., 28(3):409–413.

57. Majid, A., Burenhult, N., Stensmyr, M., de Valk, J., & Hansson, B. S. (2018) Olfactory language and abstraction across cultures. Phil. Trans. R. Soc. B, 373(1752):20170139.

58. Everett, D. L. (2005) Cultural constraints on grammar and cognition in pirahã: another look at the design features of human language. Curr. Anthropol., 46(4):621–646.

59. Wierzbicka, A. (2008) Why there are no ‘colour universals’ in language and thought. J. R. Anthropol. Inst., 14(2):407–425.

60. Regier, T., Kay, P., & Khetarpal, N. (2009) Color naming and the shape of color space. Language, 85(4):884–892.

61. Lindsey, D. T., Brown, A. M., Brainard, D. H., & Apicella, C. L. (2015) Hunter-gatherer color naming provides new insight into the evolution of color terms. Curr. Biol., 25(18):2441–2446.

62. Witzel, C. (2016) New insights into the evolution of color terms or an effect of saturation? i-Perception, 7(5). doi: 10.1177/2041669516662040.

63. Lindsey, D. T., Brown, A. M., Brainard, D. H., & Apicella, C. L. (2016) Hadza color terms are sparse, diverse, and distributed, and presage the universal color categories found in other world languages. i-Perception, 7(6). doi: 10.1177/2041669516681807.

64. Witzel, C. (2019) Variation of saturation across hue affects unique and typical hue choices. i-Perception, 10(5). doi: 10.1177/2041669519872226.

65. Webster, M. A. & Mollon, J. D. (1997) Adaptation and the color statistics of natural images. Vision Res., 37:3283–3298.

66. Webster, M. A., Mizokami, Y., & Webster, S. A. (2007) Seasonal variations in the color statistics of natural images. Network-Comp. Neural., 18(3):213–233.

67. McDermott, K. C., Malkoc, G., Mulligan, J. B., & Webster, M. A. (2010) Adaptation and visual salience. J. Vis., 10(13). doi: 10.1167/10.13.17.

68. Kay, P., Berlin, B., Maffi, L., & Merrifield, W. (1997) Color naming across languages. In Hardin, C. L. & Maffi, L., editors, Color categories in thought and language, pages 21–56. Cambridge University Press, Cambridge. ISBN 9780521498005.

69. Rubner, Y., Tomasi, C., & Guibas, L. J. A metric for distributions with applications to image databases. In IEEE Sixth International Conference on Computer Vision, pages 59–66, Bombay, India, (1998).

70. Peterman, W. E. (2018) ResistanceGA: An R package for the optimization of resistance surfaces using genetic algorithms. Methods Ecol. Evol., 9(6):1638–1647.

71. Zheng, B. & Agresti, A. (2000) Summarizing the predictive power of a generalized linear model. Statist. Med., 19(13):1771–1781.

72. Nakagawa, S. & Schielzeth, H. (2013) The coefficient of determination *R*^2^ and intra-class correlation coefficient from generalized linear mixed-effects models revisited and expanded. Methods Ecol. Evol., 4(2):133–142.

73. Nakagawa, S., Johnson, P. C. D., & Schielzeth, H. (2017) The coefficient of determination *R*^2^ and intra-class correlation coefficient from generalized linear mixed-effects models revisited and expanded. J. R. Soc. Interface, 14(134). doi: 10.1098/rsif.2017.0213.

74. Row, J. R., Knick, S. T., Oyler-McCance, S. J., Lougheed, S. C., & Fedy, B. C. (2017) Developing approaches for linear mixed modeling in landscape genetics through landscape-directed dispersal simulations. Ecol. Evol., 7(11):3751–3761.

## References

1. Shannon, C. E. (1948) A mathematical theory of communication. Bell System Technical Journal, 27(3):379–423.

2. Shannon, C. E. (1959) Coding theorems for a discrete source with a fidelity criterion. IRE National Convention Record, 7(4):142–163.

3. Banerjee, A., Merugu, S., Dhillon, I. S., & Ghosh, J. (2005) Clustering with bregman divergences. J. Mach. Learn. Res., 6:1705–1749.

4. Yendrikhovskij, S. N. (2001) Computing color categories from statistics of natural images. J. Imaging Sci. Technol., 45(5):409–417.

5. Steels, L. & Belpaeme, T. (2005) Coordinating perceptually grounded categories through language: a case study for colour. Behav. Brain Sci., 28(4):469–489.

6. Regier, T., Kay, P., & Khetarpal, N. (2007) Color naming reflects optimal partitions of color space. PNAS, 104(4):1436–1441.

7. Zaslavsky, N., Kemp, C., Regier, T., & Tishby, N. (2018) Efficient compression in color naming and its evolution. PNAS, 115(31):7937–7942.

8. Boynton, R. M. & Olson, C. X. (1987) Locating basic colors in the osa space. Color Res. Appl., 12(2):94–105.

9. Sturges, J. & Whitfield, T. W. A. (1995) Locating basic colours in the munsell space. Color Res. Appl., 20(6):364–376.

10. Lindsey, D. T. & Brown, A. M. (2014) The color lexicon of american english. J. Vision, 14(2):17, 1–25.

11. Abbott, J. T., Griffiths, T. L., & Regier, T. (2016) Focal colors across languages are representative members of color categories. PNAS, 113(40):11178–11183.

12. Agarwal, A. & Daume III, H. (2010) A geometric view of conjugate priors. Mach Learn, 81:99–113.

13. Arimoto, S. (1972) An algorithm for computing the capacity of arbitrary discrete memoryless channels. IEEE Trans. Inf. Theory, 18(1):14–20.

14. Blahut, R. (1972) Computation of channel capacity and rate-distortion function. IEEE Trans. Inf. Theory, 18(4):460–473.

15. Csiszár, I. & Tusnády, G. (1984) Information geometry and alternating minimization procedures. Statistics and Decisions, Supplement Issue 1:205–237.

16. Byrne, C. L. (2014) Iterative optimization in inverse problems. CRC Press, Boca Raton, FL.

17. Zaslavsky, N., Kemp, C., Tishby, N., & Regier, T. (2019) Communicative need in colour naming. Cogn. Neuropsychol., 37(5-6):312–324.

18. Powell, M. J. D. The BOBYQA algorithm for bound constrained optimization without derivatives. Technical report, Department of Applied Mathematics and Theoretical Physics, Cambridge University, UK, (2009).

19. Kay, P., Berlin, B., Maffi, L., & Merrifield, W. (1997) Color naming across languages. In Hardin, C. L. & Maffi, L., editors, Color categories in thought and language, pages 21–56. Cambridge University Press, Cambridge. ISBN 9780521498005.

20. Gibson, E., Futrell, R., Jara-Ettinger, J., Mahowald, K., Bergen, L., Ratnasingam, S., Gibson, M., Piantadosi, S. T., & Conway, B. R. (2017) Color naming across languages reflects color use. PNAS, 114(40):10785–10790.

21. Everett, D. L. (2005) Cultural constraints on grammar and cognition in pirahã: another look at the design features of human language. Curr. Anthropol., 46(4):621–646.

22. Wierzbicka, A. (2008) Why there are no ‘colour universals’ in language and thought. J. R. Anthropol. Inst., 14(2):407–425.

23. Regier, T., Kay, P., & Khetarpal, N. (2009) Color naming and the shape of color space. Language, 85(4):884–892.

24. Sumner, P. & Mollon, J. D. (2000) Catarrhine photopigments are optimized for detecting targets against a foliage background. J. Exp. Biol., 203(13):1963–1986.

25. Sumner, P. & Mollon, J. D. (2000) Chromaticity as a signal of ripeness in fruits taken by primates. J. Exp. Biol., 203(13):1987–2000.

26. Witzel, C. (2019) Variation of saturation across hue affects unique and typical hue choices. i-Perception, 10(5). doi: 10.1177/2041669519872226.

